# Intermediary role of lung alveolar type 1 cells in epithelial repair upon Sendai virus infection

**DOI:** 10.1101/2021.08.04.455124

**Authors:** Belinda J Hernandez, Margo P Cain, Jose R Flores, Michael J Tuvim, Burton F Dickey, Jichao Chen

## Abstract

The lung epithelium forms the first barrier against respiratory pathogens and noxious chemicals; however, little is known about how >90% of this barrier – made of alveolar type 1 (AT1) cells – responds to injury, in contrast to our accumulating knowledge of epithelial progenitor and stem cells whose importance lies in their ability to restore the barrier. Using Sendai virus to model natural infection in mice, we combine 3D imaging, lineage-tracing, and single-cell genomics to show that AT1 cells have an intermediary role by persisting in areas depleted of alveolar type 2 (AT2) cells, mounting an interferon response, and receding from invading airway cells. Sendai virus infection mobilizes airway cells to form alveolar SOX2+ clusters without differentiating into AT1 or AT2 cells, as shown in influenza models. Intriguingly, large AT2-cell-depleted areas remain covered by AT1 cells, which we name “AT2-less regions”, and are replaced by SOX2+ clusters spreading both basally and luminally around AT1 cell extensions. AT2 cell proliferation and differentiation are largely confined to topologically distal regions – the end of airspace that could be in the periphery or middle of the lung – and form de novo alveolar surface, with limited contribution to in situ repair of AT2-less regions. Time course single-cell RNA-seq and AT1-cell interactome analyses suggest enhanced recognition of AT1 cells by immune cells and altered growth signals. Our comprehensive spatiotemporal and genome-wide study highlights the hitherto unappreciated role of AT1 cells during Sendai virus infection and possibly other injury-repair processes.

## INTRODUCTION

Individual cell types in a multicellular organ must act in concert to fulfill physiological functions and react likewise to injuries. This holistic reaction is expected to reach its full extent in response to naturally-occurring pathogens, possibly due to evolutionary pressure, as opposed to engineered cell ablation using diphtheria toxin or adapted pathogens from noncognate species. Sendai virus (SeV), also known as murine parainfluenza virus type 1, is used for gene delivery and to model chronic lung diseases, but also provides a natural injury-repair model (Faisca and Desmecht, 2007; Holtzman et al., 2005). A comprehensive understanding of spatiotemporal and cell-type-specific responses to SeV, which is the focus of this study, will not only better establish a disease model to test therapeutics but also shed light on natural infections in humans, such as influenza and COVID-19.

As the first barrier against microbial and chemical agents, the otherwise quiescent lung epithelium has evolved robust repair mechanisms that, in the alveolar region, activate a facultative stem cell population – alveolar type 2 (AT2) cells, which have attracted most attention in the field (Barkauskas et al., 2013; Desai et al., 2014; Hogan et al., 2014). In contrast, alveolar type 1 (AT1) cells, constituting 95% of the alveolar surface, are often considered a passive structural component to be injured and then replaced by AT2 cells, but are likely a key component of the lung’s holistic reaction to injury due to their sheer mass (Chen, 2017; Vila Ellis and Chen, 2020). Our limited knowledge of AT1 cell biology in injury-repair stems partly from their expansive and ultrathin morphology that spatially uncouples the nucleus from cellular extensions such that cytokeratin, junctional, or nuclear staining alone is inadequate to study AT1 cells (Yang et al., 2016). Furthermore, the dense 3D packing of alveolar tissues obscures the connection between topologically distal and proximal regions – important spatial attributes also missed in biochemical assays of whole organs. Conceptually, the alveolar epithelium can be unfurled into a 2D surface – an imagery evoked when analogizing the human lung to a half tennis court – that is predominantly AT1 cell surface and sprinkled with AT2 cells (Weibel, 2009). Accordingly, injury-repair could be (1) in situ healing similar to wound healing as a result of cell expansion, proliferation, and migration – a process also artificially invoked during re-epithelialization of decellularized lungs (Uriarte et al., 2018), or (2) de novo growth around the border of the tennis court – a likely process during post-pneumonectomy compensatory growth where there is no direct injury to the existing surface (Lee and Rawlins, 2018; Vila Ellis and Chen, 2020). Regardless, it is integral to our understanding of lung injury-repair to delineate how the majority of the epithelial surface, or essentially the AT1 cell, responds and contributes.

In this study, we notice that upon SeV infection in mice, AT1 cells persist in large alveolar areas depleted of AT2 cells, providing an opportunity to ascertain the role of AT1 cells in injury-repair. Applying our knowledge of and tools for AT1 cells in the uninjured lung (Yang et al., 2016), we show that the surviving AT1 cells mount a robust interferon response, comparable to that by AT2 cells, and are replaced by dysplastic SOX2+ airway cells. In contrast, AT2 cells proliferate and differentiate into AT1 cells mostly in topologically distal regions, and contribute minimally to in situ healing of damaged areas. Therefore AT1 cells have an intermediary role during SeV infection by temporarily covering the tissue surface and coordinating an immune response. The resulting transcriptomic profiling and cellular analysis of AT1 cells and their relationship with other epithelial and non-epithelial cells fills in a gap in our knowledge of lung injury-repair.

## RESULTS

### Airway cells are mobilized to populate damaged alveolar regions, with minimal alveolar differentiation

Expecting SeV infection to induce an asynchronous, spatially heterogeneous response of injury-repair that needed an organ-level depiction, we used optical projection tomography to visualize whole lung lobes immunostained for the airway transcription factor SOX2 (Fig. 1A). At day 9 post infection (9 dpi), SOX2 staining diminished in lobar and segmental airways but remained in terminal bronchioles, consistent with cell loss from viral infection that spread distally. The most proximal airways near the lobe entrance had normal SOX2 staining and thus possibly had limited infection or had recovered (Fig. 1A). By 28 dpi, all airways regained SOX2 staining and, strikingly, formed cauliflower-like SOX2+ clusters that extended beyond terminal bronchioles (Fig. 1A). These outgrowing clusters were often well into the alveolar region and even reached the most distal mesothelium, although our 3D imaging showed that they were continuous with and likely originated from the airways (Fig. 1A), as further tested by lineage-tracing below. The SOX2+ clusters were visible in freshly-dissected lungs using a *Sox2*^*GFP*^ allele (Arnold et al., 2011) and were mutually exclusive with AT2 cells that were genetically labeled prior to infection – an initial, to-be-explored clue for a limited contribution of AT2 cells to restoring these damaged regions (Fig. 1A).

**Figure 1:**
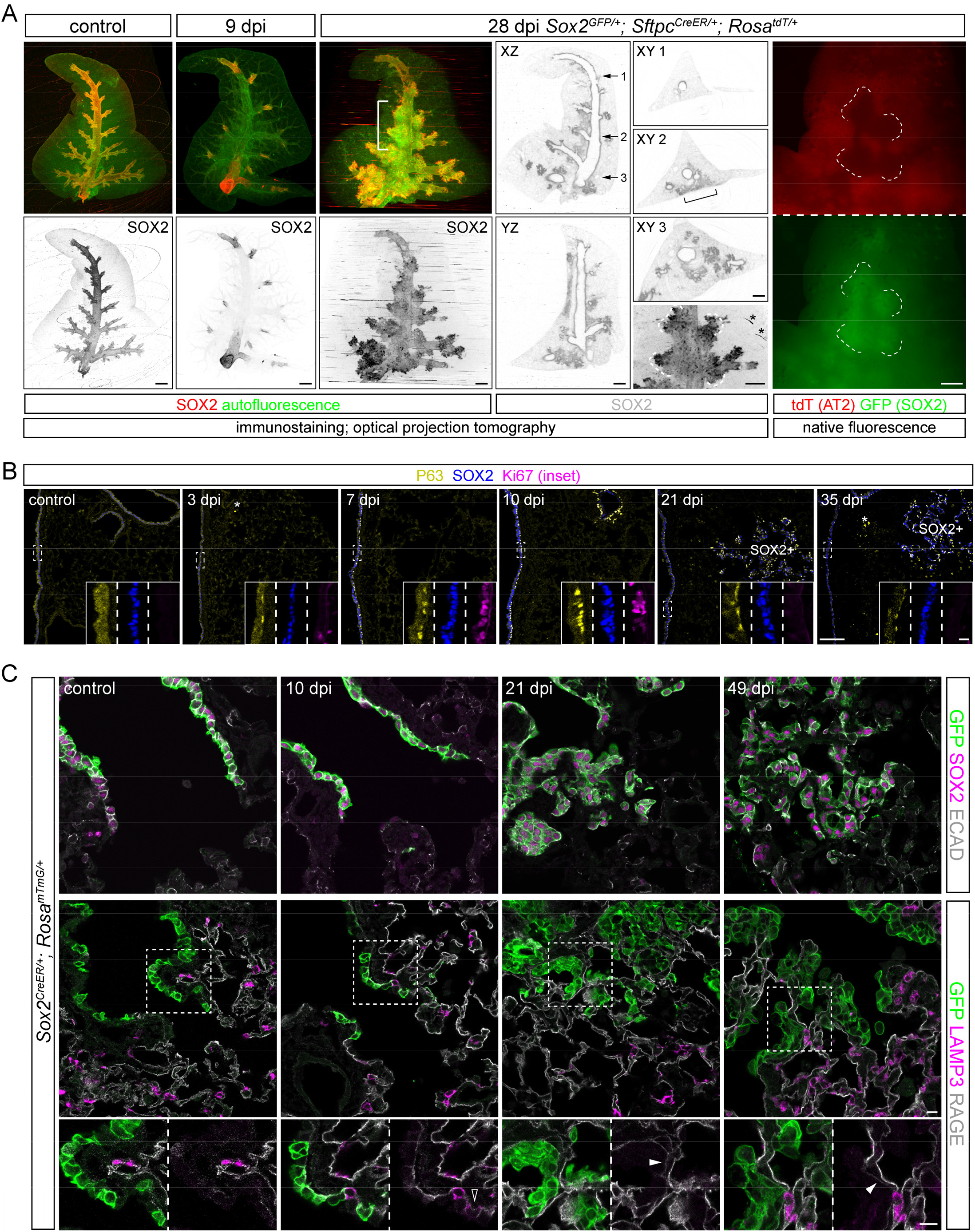
Airway cells are mobilized to populate damaged alveolar regions, with minimal alveolar differentiation. **(A)** Optical projection tomography (OPT) images of wholemount immunostained accessory lobes from control and SeV infected (dpi, day post infection) adult mouse lungs. Tissue autofluorescence highlights airways and macrovasculature, and is more noticeable at 9 dpi when SOX2 expression is lost from most airways. The recovery and cauliflower-like alveolar clusters of SOX2 expression at 28 dpi are apparent on XZ, YZ, and XY (at the numbered levels) optical planes. Alveolar SOX2 clusters can be found at lobe edges but are still continuous with the airways (bracket). The 28 dpi lung is from a *Sox2*^*GFP/+*^; *Sftpc*^*CreER/+*^; *Rosa*^*tdT/+*^ mouse and received 2 mg tamoxifen 3 weeks prior to infection. Native fluorescence images before immunostaining in the bracketed region (rightmost column) show mutual exclusion of airway cells marked by *Sox2*^*GFP*^ (the same contours outlined by dashes as those in the OPT image; asterisk: technical ring artifacts) and AT2 cells marked by *Sftpc*^*CreER*^ and *Rosa*^*tdT*^. Scale: 500 um. Images are representative of at least 3 lungs (same in subsequent figures). **(B)** Section immunostaining images showing SeV-induced proliferation (Ki67 in insets for boxed regions), expansion and persistence of P63+ basal-like cells as a subset of SOX2+ cells, in airways and alveolar SOX2 clusters (SOX2+). Asterisk: non-specific macrophage fluorescence. Scale: 100 um (10 um for insets). **(C)** Section immunostaining images of control and SeV infected adult *Sox2*^*CreER/+*^; *Rosa*^*mTmG/+*^ lungs with 2 mg tamoxifen administered 3 weeks prior to infection. All lineage-labeled cells (GFP+) are positive for the airway marker SOX2 and cuboidal/columnar as outlined by E-Cadherin (ECAD; top row), and negative for an AT2 cell marker (LAMP3) except for rare cells transiently expressing LAMP3 (open arrowhead at 10 dpi) (bottom row). Filled arrowhead: weak AT1 cell marker (RAGE) under GFP+ cells, which will be further examined later. Scale: 10 um.

The SOX2+ clusters were reminiscent of “pods” fueled by rare basal-like cells upon severe influenza infection (Kanegai et al., 2016; Kumar et al., 2011; Xi et al., 2017). Indeed, basally located, proliferative P63+ cells first appeared 3 dpi within airways and were readily found at 7 and 10 dpi (Fig. 1B). Although no longer expressing Ki67, these airway P63+ cells persisted at least 35 dpi (Fig. 1B). Mirroring the observed pattern and kinetics of airway repair (Fig. 1A), the wave of P63 expression extended to alveolar SOX2+ clusters and also persisted (Fig. 1B). The aberrant persistence of p63+ cells in bronchioles and alveoli suggested either a mimicry of larger airways that were normally populated by P63+ cells or a defect in terminating the injury-repair response, possibly contributing to the chronic abnormalities (Walter et al., 2002).

To ascertain the fate of alveolar SOX2+ cells, we used *Sox2*^*CreER*^ (Arnold et al., 2011) to label airway cells prior to infection and found most, if not all, lineage-labeled cells including those aberrantly present in the alveolar region remained SOX2+ and did not express an AT1 marker RAGE nor an AT2 marker LAMP3 (Fig. 1C). The dysplastic SOX2+ clusters often overlay a surface with weak RAGE staining (Fig. 1C), suggesting replacement of AT1 cells as examined in details later. Our whole-lobe and cellular imaging, together with lineage-tracing, captured mobilization of P63/SOX2 airway cells to repair damaged airway and alveolar regions without noticeable differentiation into alveolar cells.

### AT2-less regions are depleted of AT2 cells, but covered by AT1 cells

The rapidly growing, dysplastic SOX2+ clusters prompted us to search for damaged alveolar regions that were presumably in need of repair. A time-course survey of AT1 (marked by RAGE) and AT2 (marked by SFTPC) cells revealed regions depleted of AT2 cells at 7 dpi and spanning hundreds of microns – comparable in size to future SOX2+ clusters (Fig. 2A). Strikingly, these AT2-depleted regions were indistinguishable from the rest of the lung based on RAGE staining, suggesting continued coverage with AT1 cells (Fig. 2A). This uncoupling between AT1 and AT2 cells was confirmed on sequential sections with another set of AT1 and AT2 markers – PDPN and LAMP3, although PDPN additionally marked P63+ cells at 14 dpi as expected for a basal cell marker (Kanner et al., 2010) (Fig. 2A). Besides their cell membranes, AT1 cell nuclei also persisted, as marked by the lung lineage transcription factor NKX2-1 without surrounding SFTPC (Little et al., 2019; Little et al., 2021) (Fig. 2B). To confirm that RAGE staining was from AT1 cells instead of aberrant gene expression upon infection, we used our recently characterized AT1 cell driver *Rtkn2*^*CreER*^ (Little et al., 2021) and labeled AT1 cells prior to infection. Indeed, regions depleted of AT2 cells were as fully covered with lineage-labelled AT1 cells as uninvolved regions (Fig. 2C). Accordingly, we referred to these AT2-cell depleted but AT1-cell covered areas “AT2-less regions”.

**Figure 2:**
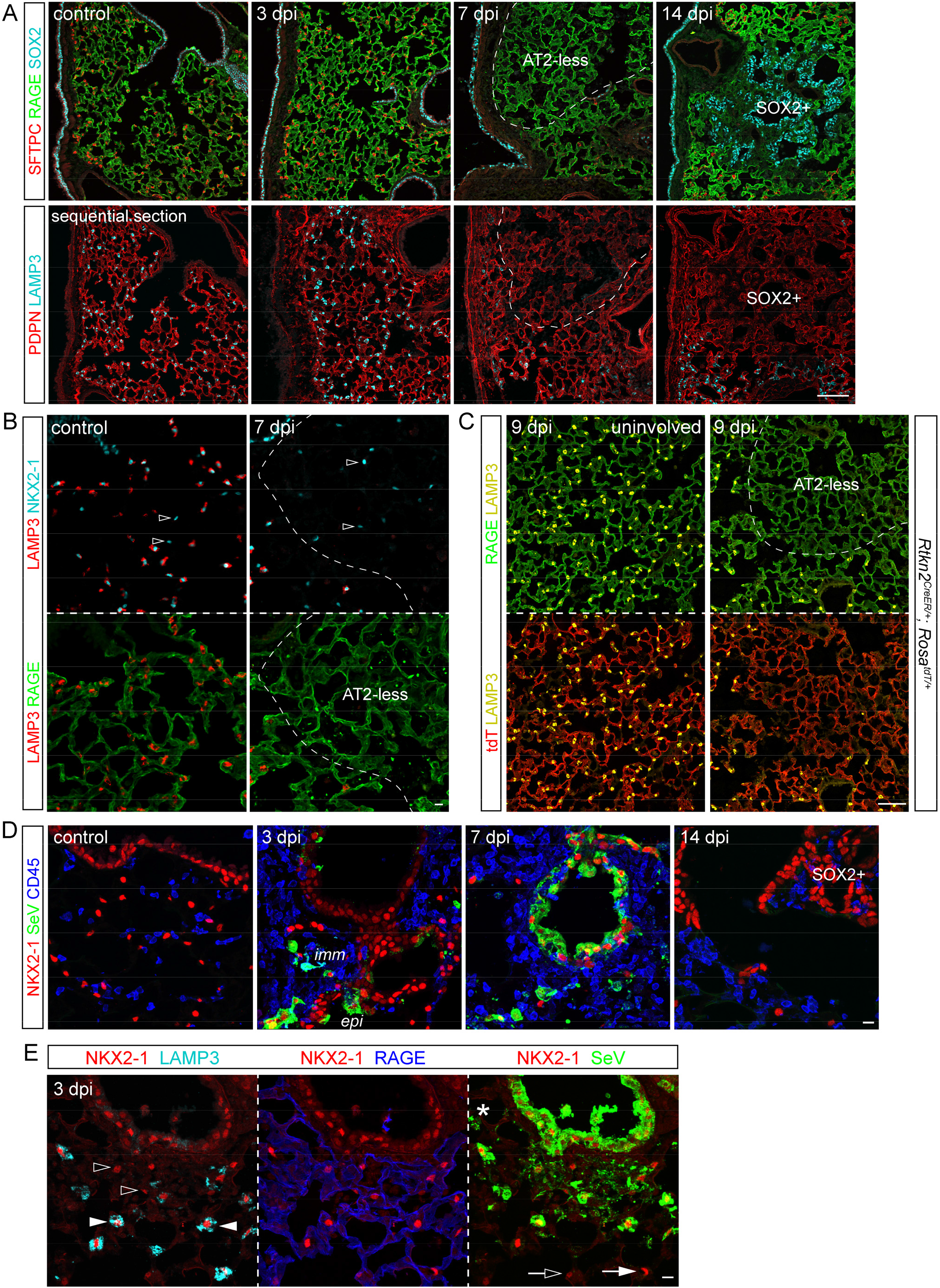
AT2-less regions are depleted of AT2 cells, but covered by AT1 cells. **(A)** Sequential section immunostaining images showing SeV infection depletes dashed areas of AT2 cell markers (SFTPC and LAMP3) without affecting AT1 cell markers (RAGE and PDPN) at 7 dpi. These AT2 cell depleted (AT2-less) regions are similar in size and location to SOX2+ clusters at 14 dpi. PDPN, a basal cell marker, also marks P63+ basal-like cells. Scale: 100 um. **(B)** Section immunostaining images showing AT1 cell nuclei (NKX2-1+ LAMP3-; open arrowheads) are intermingled with AT2 cell nuclei (NKX2-1+ LAMP3+) in the control lung, but are the only nuclei present in AT2-less regions (dashes). Scale: 10 um. **(C)** Section immunostaining images of *Rtkn2*^*CreER/+*^; *Rosa*^*tdT/+*^ lungs with recombination induced at P5 with 500 ug tamoxifen to efficiently label AT1 cells. At 9 dpi, both uninvolved (left) and AT2-less (right; depleted of LAMP3) regions are covered by tdT-labelled RAGE+ AT1 cells. Scale: 100 um. **(D)** Section immunostaining images showing SeV viral proteins (SeV) in immune cells (imm; CD45+) and epithelial cells (epi; NKX2-1+) at 3 and 7 dpi, and cleared by 14 dpi. Scale: 10 um. **(E)** Section immunostaining images showing both thin AT1 cells (open arrowheads; RAGE+ NKX2-1+) and cuboidal AT2 cells (filled arrowheads; LAMP3+ NKX2-1+) are filled with SeV viral proteins and thus infected, in comparison with their uninfected neighbors (open and filled arrows). Asterisk, an infected macrophage. Scale: 10 um.

A possible explanation for the preferential depletion of AT2 cells was differential viral infection or clearance of infected cells, although the receptor for SeV is sialic acid and widely present (Markwell and Paulson, 1980). Examining cell-type infection in vivo was challenging and an underestimate due to the transient nature of infected/dying cells compounded by the likely concomitant decrease of cellular markers. Nevertheless, abundant SeV proteins were readily detected in NKX2-1+ epithelial cells and CD45+ immune cells at 3 dpi and spread to adjacent cells at 7 dpi before largely disappearing at 14 dpi (Fig. 2D). Infected airway and AT2 cells were identified based on their anatomic location, cell shape as filled by SeV proteins, and the AT2 marker LAMP3 (Fig. 2E). Although infrequent, infected AT1 cells were identifiable when the SeV protein staining outlined both the NKX2-1+ nucleus and RAGE+ cellular extensions (Fig. 2E). Therefore, although AT1 cells had a larger surface area accessible to SeV and could be infected, AT2 cells either were more easily infected or survived less, leading to AT2-less regions; the possibility of AT2-cell depletion via their differentiation into AT1 cells was tested and excluded below. As a result, the exposed tissue surface from dying AT1 and AT2 cells was expected to be resealed by remaining AT1 cells, as supported by the continuous RAGE staining (Fig. 2).

### AT2 cells differentiate into AT1 cells in topologically distal regions, but contribute minimally to in situ repair of damaged alveolar surface

As stem cells, AT2 cells were known to proliferate and differentiate into AT1 cells, and were often assumed to do so to repair damaged regions (Hogan et al., 2014). However, as introduced earlier, AT2 cells could be also activated upon pneumonectomy without direct injury and such activation was more apparent near the tissue edge, which was topologically most distal and had more space to expand (Filipovic et al., 2013; Liu et al., 2016). The AT2-less regions in the SeV model provided a landmark to locate activated AT2 cells. At 14 dpi, expanding SOX2+ clusters were still bordered by regions without AT2 cells, let alone activated ones (Fig. 3A). Instead, clusters of AT2 cells formed further away – near the organ edge or major blood vessels and airways, both of which were the most topologically distal part of the respiratory tree even when embedded in the center of the lung (Fig. 3A). These aberrant AT2 cell clusters were intermingled with small, LAMP3-negative AT1 cells (Fig. 3A), an arrangement reminiscent of that in nascent alveolar sacs during development (Yang et al., 2016), and were still present at 42 dpi when the AT2-less regions had disappeared (Fig. 3A). This AT2 cell depletion followed by hyperplasia was evident when tracking the ratio of AT2 to AT1 cells over time and was specific to damaged regions (Fig. 3B). Therefore, AT2 cells clustered de novo near vessels, airways, and the edges of the tissue, but at a distance from AT2-less regions (Fig. 3C).

**Figure 3:**
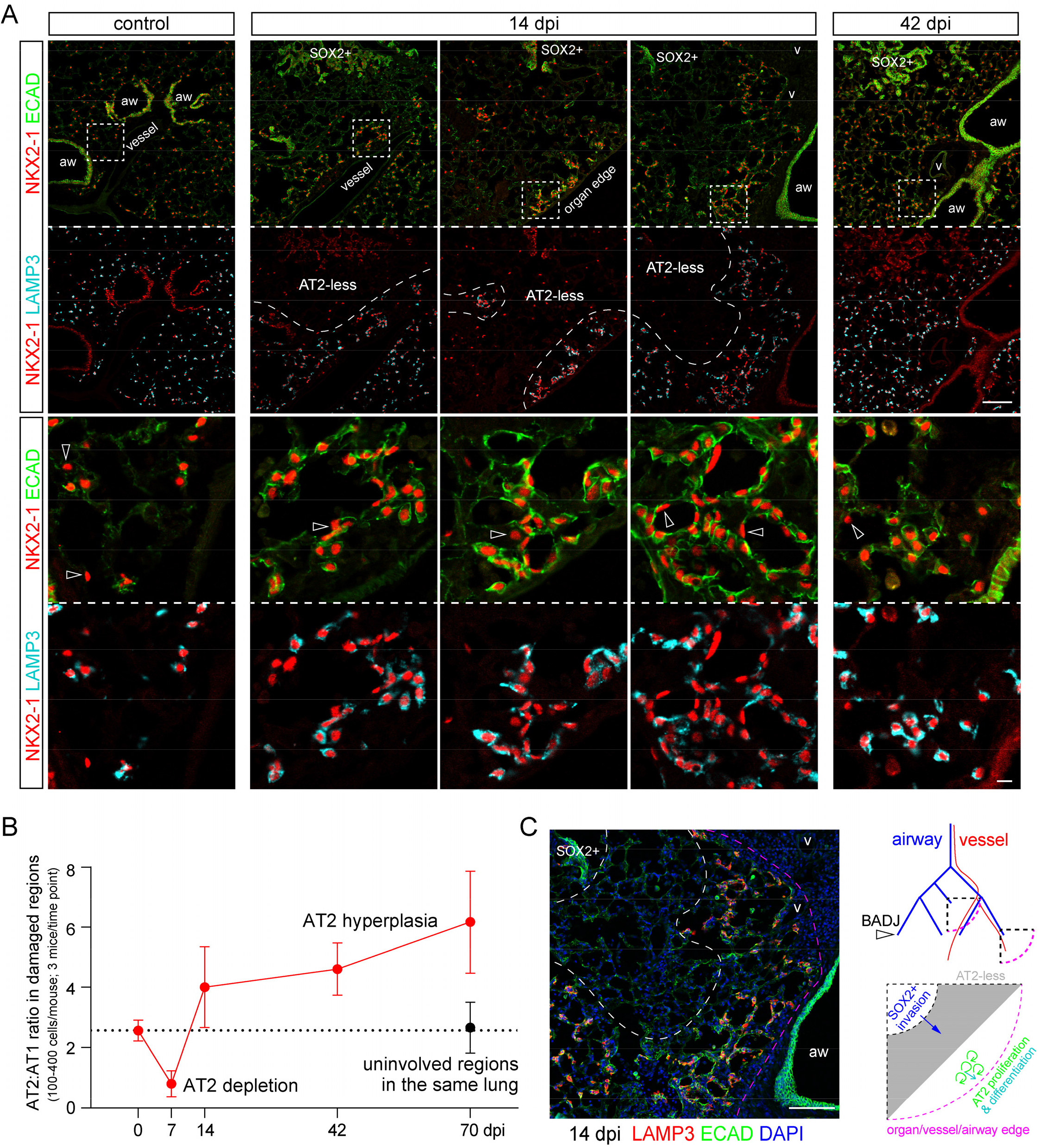
AT2 cells form clusters in topologically distal regions near the organ edge or major blood vessels and airways. **(A)** Section immunostaining images with boxed topologically distal regions enlarged. AT2 cells are LAMP3+ and cuboidal as outlined by E-Cadherin (ECAD), as opposed to AT1 cells (open arrowheads). Three examples of 14 dpi lung sections show that AT2 cells cluster around a nascent airspace lumen away from AT2-less regions and SOX2 clusters, but near the organ edge or major blood vessels (v) and airways (aw), a pattern only partially resolved at 42 dpi. Scale: 10 um. **(B)** Quantification of the ratio of AT2 to AT1 cells supports AT2 cell depletion at 7 dpi and subsequent hyperplasia and clustering. Uninvolved regions in infected lungs have a normal cell ratio. Error bars correspond to standard deviations. **(C)** An image from a 14 dpi lung in (**A**) overlaid with DAPI to demonstrate topologically distal regions beyond a SOX2 cluster and an AT2-less region, corresponding to the center quarter circle in the diagram that is topologically equivalent to the quarter circle near the organ edge. BADJ, bronchoalveolar duct junction. Only arterial vessels are drawn for clarity. It is predicted and subsequently tested that AT2-less regions are invaded by SOX2 cells, while AT2 cell proliferation and differentiation occur further distally.

This conclusion was substantiated by lineage-tracing experiments showing that AT2 cell clusters including adjacent AT1 cells arose from preexisting AT2 cells that were labeled prior to infection, whereas no expansion of labeled AT2 cells nor their differentiation into AT1 cells was observed in AT2-less regions (Fig. 4). Taken together, at least in the SeV model, AT2 cell-mediated tissue growth occurred in topologically distal regions and away from obviously damaged regions, possibly to avoid generating excessive AT1 cells given the persistence of AT1 cells (Fig. 2). Our analysis also highlighted the importance of spatial information in distinguishing de novo growth versus in situ repair.

**Figure 4:**
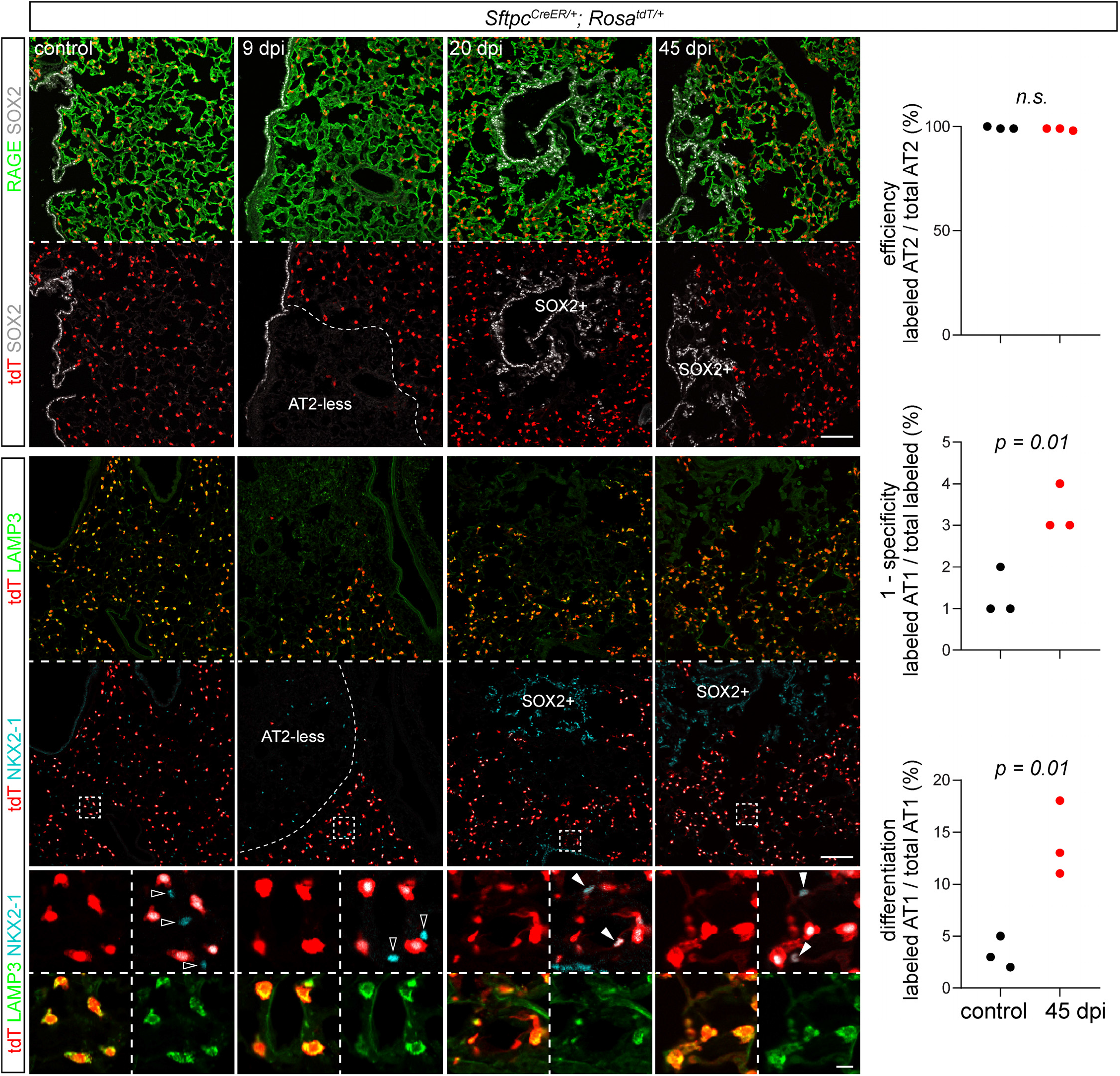
AT2 cells differentiate into AT1 cells in topologically distal regions but not AT2-less regions. Section immunostaining images from control and SeV infected lungs of *Sftpc*^*CreER/+*^; *Rosa*^*tdT/+*^ mice that received 2 mg tamoxifen 3 weeks before infection. Lineage-labeled AT2 cells are excluded from AT2-less regions (RAGE+ LAMP3-; 9 dpi) and later SOX2 clusters (20 and 45 dpi), but are present and differentiate into AT1 cells (open arrowheads; NKX2-1+ LAMP3-) in topologically distal regions (boxed) as explained in Fig. 3. Scale: 100 um (10 um for enlarged boxed regions). Quantification shows near-complete labeling of AT2 cells and their increased differentiation into AT1 cells upon infection, constituting 10-20% AT1 cells in topologically distal regions. Labeling of AT1 cells in the control is due to cumulative baseline cell turnover that is additionally amplified by tamoxifen-independent, leaky recombination of *Sftpc*^*CreER*^.

### Invading airway cells displace AT1 cells via basal and luminal spreading

As AT2-less regions, without being repaired by AT2 cells, disappeared with the same kinetics as the expansion of SOX2+ clusters (Fig. 3A), we posited that the persistent and then disappearing AT1 cells were displaced by the invading SOX2+ cells. Indeed, at 12 dpi when SOX2+ clusters were first detected consistently in the alveolar region, SOX2+ cells were in three conformations relative to RAGE+ AT1 cell extensions: (1) basal, in which SOX2+ cells were underneath an intact RAGE membrane and embedded in a collagen matrix; (2) luminal I, in which 2-3 SOX2+ cells gained access to the airspace, accompanied by weakened or gapped RAGE staining; and (3) luminal II, in which tens of SOX2+ cells accumulated in the airspace and spilled over AT1 cell surfaces with normal RAGE staining, possibly causing further AT1 cell displacement (Fig. 5A). Although live imaging of this displacement was not available, SOX2+ cells of the two luminal conformations were connected by adherens junctions and continuous with those embedded in the tissue, suggesting that the three conformations represented snapshots of a continuous, if not sequential, process (Fig. 5B).

**Figure 5:**
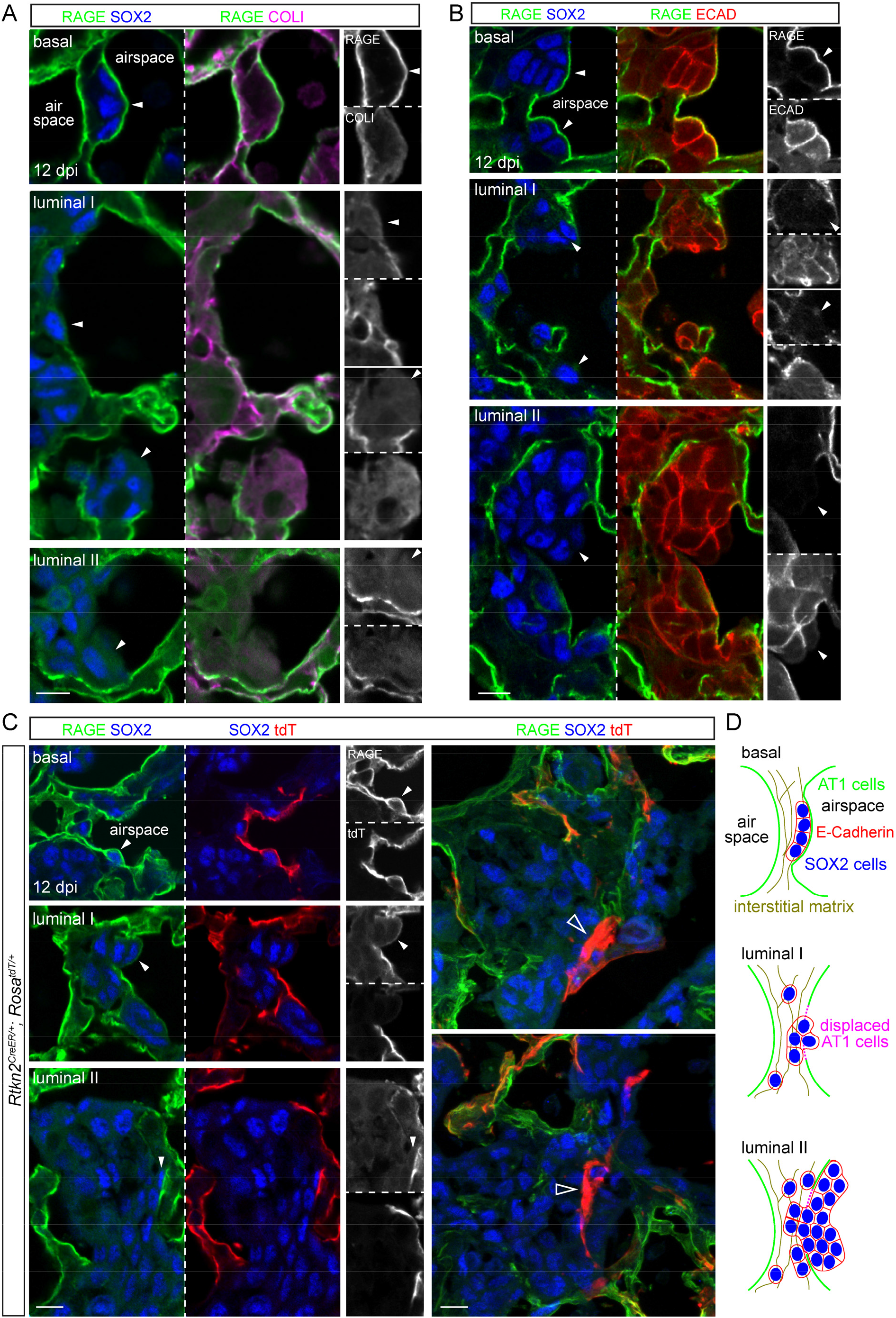
Invading airway cells displace AT1 cells via basal and luminal spreading. Section immunostaining images of lungs at 12 dpi when SOX2 clusters are reliably detected. (**A**) Invading SOX2+ cells (arrowheads) are found embedded in the interstitial matrix (COLI) and basal to the AT1 cell membrane (RAGE), gaining access to the airspace and covered by reduced RAGE (luminal I), or accumulated in the airspace and spreading over neighboring RAGE membrane (luminal II). (**B**) Invading SOX2+ cells of the 3 conformations form continuous cell junctions (ECAD). (**C**) Left, RAGE staining is from AT1 cells lineage-labeled at P5 with 500 ug tamoxifen. Right, AT1 cells surrounded by SOX2+ cells accumulate brighter tdT (open arrowheads), possibly amid cell displacement. (**D**) Schematics showing the relationship of SOX2+ cells and AT1 cells. Scale: 10 um.

Lineage labeling using the AT1 cell driver *Rtkn2*^*CreER*^ (Little et al., 2021) confirmed that the displaced RAGE staining was from AT1 cells, and that SOX2+ cells existed basal or luminal to AT1 cells (Fig. 5C). Strikingly, at their interface where AT1 cell displacement was hypothesized to occur, shrunken fragments of AT1 cells accumulated an excessive amount of the lineage reporter, perhaps as a result of cell condensation before its shedding (Fig. 5C). Our observations were consistent with a model where AT1 cells in AT2-less regions temporarily sealed the alveolar surface before being displaced by airway-originated SOX2+ clusters spreading both basally and luminally (Fig. 5D).

### AT1 cells, as well as AT2 cells, mount an acute interferon response, as revealed by time-course single-cell RNA-seq

To examine cellular and molecular dynamics upon SeV infection in an unbiased manner, we profiled all major lung cell types using single-cell RNA-seq (scRNA-seq) at 7, 14, and 49 dpi, representing the injury, repair, and chronic phases, respectively (Fig. 6A, Fig. S1). Several infection-induced cell clusters were readily identified and included (1) *Trp63*+ basal-like cells at 14 and 49 dpi (cluster 4), corresponding to the invading SOX2+ clusters (Fig. 1); (2) *Ifi27l2a*+ cells mainly at 7 dpi for both AT1 (clusters 14 and 15) and AT2 (clusters 6 and 11) cells, corresponding to an acute interferon response to SeV; and (3) *Mki67*+ proliferative cells that were present as a result of homeostatic cell turnover but increased upon infection at 14 dpi, consistent with de novo clustering and hyperplasia of AT2 cells (Fig. 3, 4). Intriguingly, at 14 dpi, AT1 cells included a cell cluster 12 that shared genes with normal AT2 cells, such as *Napsa* and *H2-Aa*; such residual expression of AT2 genes raised the possibility that they arose from AT2 cells, as observed in regions of de novo growth (Fig. 4). Direct comparison of AT1 cells at each time point revealed induction of additional interferon genes predominantly at 7 dpi, including those for antigen processing and presentation (*Psmb8/9/10, B2m, Cd74, H2-K1, H2-D1*, and *H2-Q6*), antiviral effectors (*Ifit1, Oasl2, Rsad2*, and *Rtp4*), and interferon pathway transcription factors (*Irf1* and *Irf7*) (Fig. 6B and Table S1-3). In contrast, AT1 cell-enriched genes including *Vegfa* and *Abca1* were reduced, suggesting reduction of angiogenic and cholesterol transport functions upon infection (Bates et al., 2005; Vila Ellis et al., 2020).

**Figure 6:**
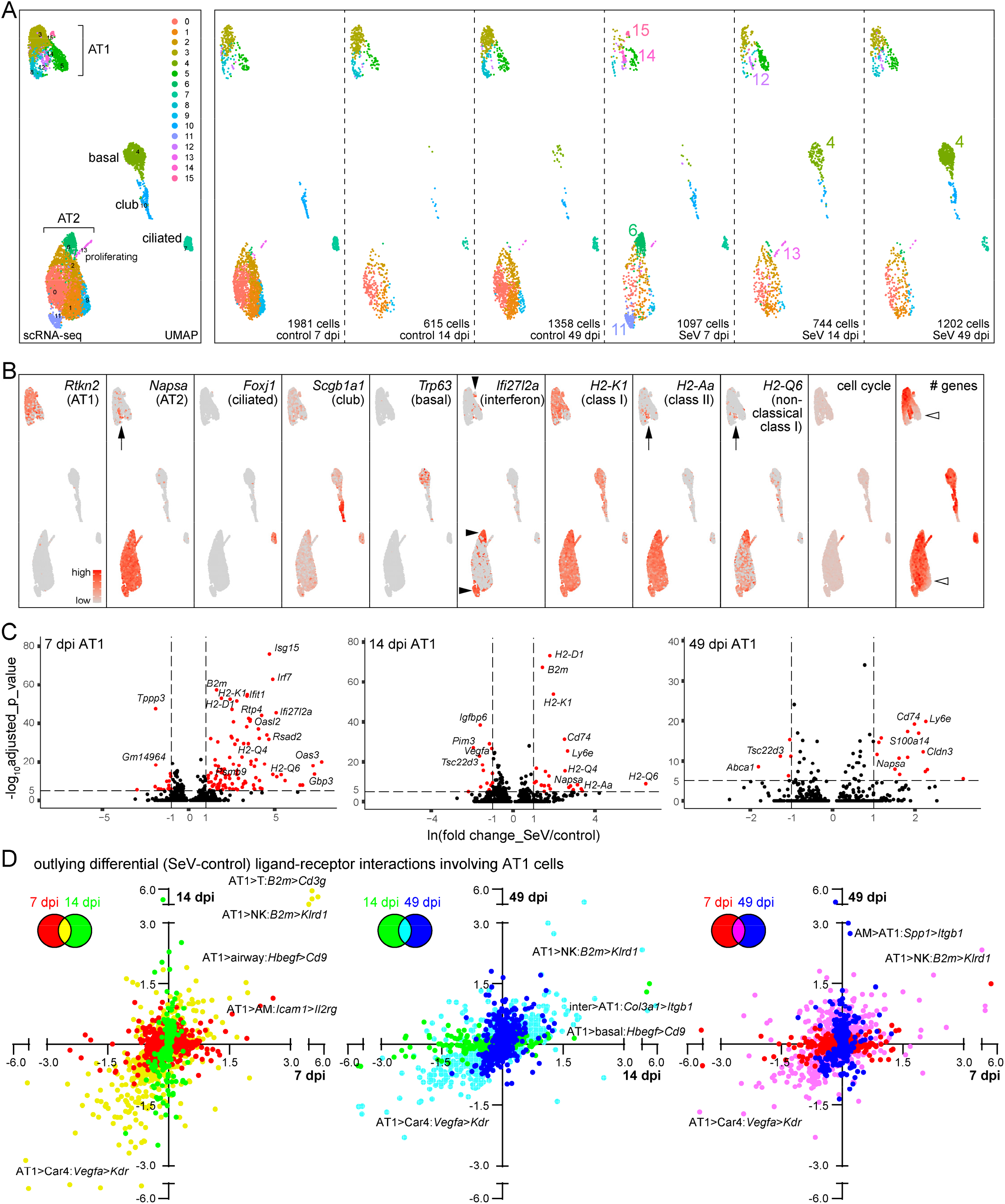
Time-course single-cell RNA-seq reveals interferon and signaling responses in AT1 cells. **(A)** Merged and split UMAP (uniform manifold approximation and projection) graphs of epithelial cells from paired control and SeV-infected lungs at 7, 14, and 49 dpi, color-coded for computer-identified clusters with differential clusters numbered in the split UMAP graphs. Cluster 4 corresponds to persistent basal-like cells. Clusters 6, 11, 14, and 15 are prominent for 7 dpi SeV lungs, clusters 13 and 12 are predicted as proliferating and differentiating AT2 cells, respectively. **(B)** Feature plots of cell-type-specific markers, supporting cell type names in (**A**). Cluster 14 (arrows) expresses markers of AT2 cells including *Napsa*, an MHC class II gene *H2-Aa*, and an MHC non-classical I gene *H2-Q6*, the last of which has highest expression in clusters 6 and 11. An interferon signature gene *Ifi27I2a* is highest (filled arrowheads) in clusters 6, 11, and 15 (lower in 14) at 7 dpi. Cell cycle gene score marks the proliferative cluster 13. Additional heterogeneity arises from the total number of genes detected (open arrowheads). **(C)** Volcano plots comparing AT1 cells from control and infected lungs at 7, 14, and 49 dpi. See Table S1, S2, S3 for complete data. **(D)** Pairwise interactome analyses of outlying changes in ligand-receptor interactions (directionality indicated with “>” for labeled examples) involving AT1 cells upon infection, color-coded for unique and shared changes as in the inset diagram. See Table S4, S5 for complete data.

To place these differentially expressed genes in the context of all lung cell types and focus on intercellular signaling, we applied our interactome algorithm (Cain et al., 2020) to identify outlying changes in ligand-receptor interactions with AT1 cells receiving or sending signals. Pairwise comparison of data from 7, 14, and 49 dpi highlighted altered interactions unique and common to two or all time points (Fig. 6C and Table S4, S5), supporting an increased recognition of AT1 cells by NK and T cells and a decreased AT1 cell-derived angiogenic signal toward the endothelium. Intriguingly, AT1 cells at 14 dpi upregulated *Hbegf*, a possible mitotic signal for AT2 cells or invading airway cells (Leslie et al., 1997). Increased *Icam1* expression in AT1 cells might facilitate immune cell extravasation (Gahmberg, 1997). With respect to AT1 cells receiving signals, a number of matrix genes including *Tnc, Fn1, Fbn1, Col1a1*, and *Col5a2* were upregulated in various mesenchymal cells, potentially affecting tissue stiffness and epithelial cell adhesion; whereas *Il18* from immune cells could stimulate inflammation via *Il18r1* in epithelial cells (Krasna et al., 2005) (Table S5). Taken together, our time-course scRNA-seq profiling confirmed our imaging-based molecular and cellular characterization and implicated active signaling roles of AT1 cells in countering SeV infection.

## DISCUSSION

In this study, we use 3D imaging, lineage-tracing, and single-cell transcriptomics to delineate the spatiotemporal and cellular heterogeneity in the lung’s response to natural viral infection. As diagramed in Fig. 7, the lung epithelium, when viewed as a 2D surface, is predominantly made of AT1 cells with minor contributions from SOX2+ airway cells and AT2 cells. SeV infection causes wide-spread loss of airway and AT2 cells, but most AT1 cells persist – forming AT2-less regions. SOX2+/P63+ basal-like cells expand to heal the airways and aberrantly extend to the AT2-less regions, displacing AT1 cells without differentiating into alveolar cells. AT2 cells form hyperplastic clusters and undergo AT1 cell differentiation in topologically distal regions, generating de novo alveolar surface with limited contribution to healing the AT2-less regions. AT1 cells have intermediary structural and signaling roles by sealing the alveolar surface temporarily and mounting an antiviral interferon response and possibly modulating growth signals, such as *Vegfa* and *Hbegf*. Such molecular and cellular understanding is necessary to dissect associated mechanisms and test therapies using the SeV or other models of lung injury-repair.

**Figure 7:**
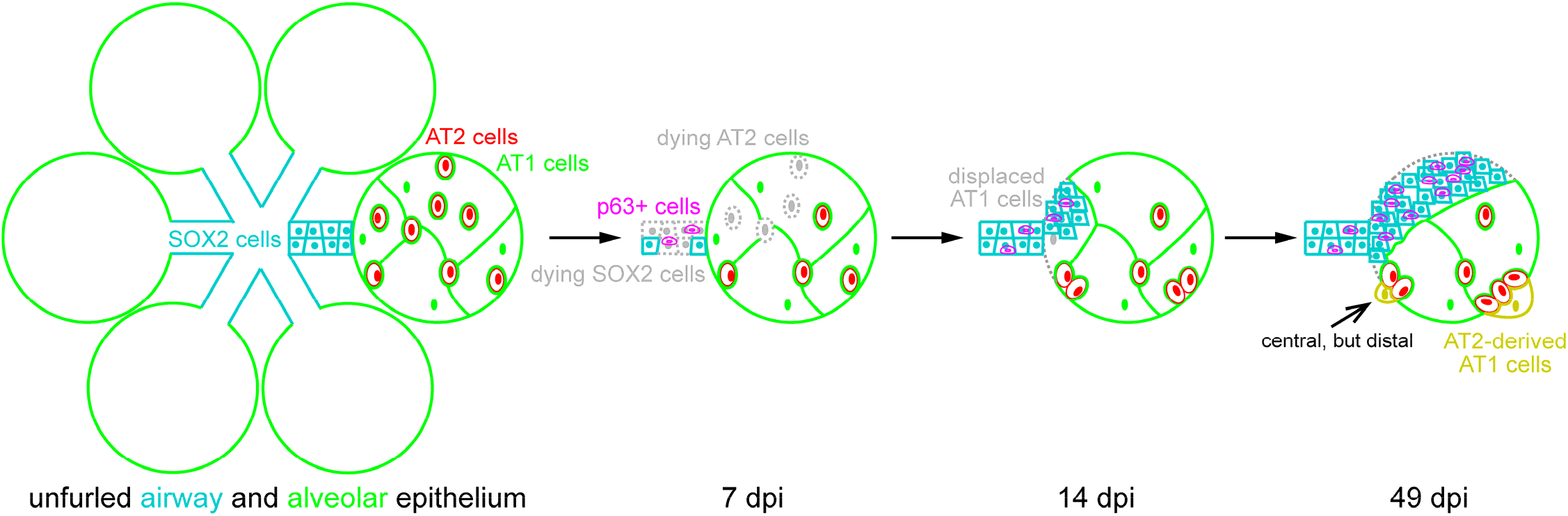
Diagram illustrating cellular changes in the lung epithelium upon Sendai virus infection. Unfurled lung epithelium consists of proximal SOX2+ columnar airway cells and distal intermingled flat AT1 and cuboidal AT2 cells where a central region can be topologically distal. Upon SeV infection, loss of airway cells amplifies P63+ basal-like cells, while loss of AT2 cells, but not AT1 cells, forms AT2-less regions. Subsequently, SOX2+ airway cells including P63+ basal-like cells invade alveoli and displace AT1 cells; AT2 cells proliferate in topologically distal regions and differentiate into AT1 cells to form de novo alveolar surface.

A notable feature in the SeV model is the formation of AT2-less regions where AT2 cells are lost while AT1 cells are spared. The initial gaps left by disappearing AT2 cells can be readily sealed by AT1 cells given their ability to expand their surface by over 10-fold during development (Yang et al., 2016). The resulting AT2-less surface must be sensed and then induce the invasion of SOX2+/P63+ cells. Loss of AT2 cells is unlikely the only trigger as genetic ablation of AT2 cells stimulates clonal expansion of remaining AT2 cells but not airway cell invasion (Barkauskas et al., 2013). Hypoxia and Fgf signaling pathways have been implicated in influenza and bleomycin models (Xi et al., 2017; Yuan et al., 2019), although it will be necessary to map their activation spatiotemporally and determine their sufficiency in providing a directional cue. The correlation between AT2-less regions and SOX2+ clusters in their size, location, and kinetics, as well as their physical interactions (Fig. 1, 2, 5), supports the replacement of AT1 cells by SOX2+ cells. However, we cannot rule out that areas bordering the AT2-less regions could be populated by spared AT2 cells nearby, although this scenario predicts that AT2 cells migrate across AT1 cells for dispersion as they do not differentiate into AT1 cells in large numbers (Fig. 4). Future live imaging experiments will provide definitive evidence for the cellular sources and interactions during the repair of AT2-less regions. It also remains to be determined if similar AT2-less regions form in other microbial and chemical injury models.

The existence of AT2-less regions also highlights the uncoupling of AT2 cell activation and in situ healing of preexisting-and-then-damaged alveolar surface. We notice AT2 cell proliferation and differentiation in topologically distal regions – close to tissue edges, airways, or vessels (Fig. 3, 4), the last of which have been suggested as a niche for AT2 cells partly due to their spatial proximity (Lee et al., 2014; Nabhan et al., 2018). This de novo formation of alveolar surface is intuitive during compensatory growth induced by pneumonectomy, but is less appreciated in models of chemical and microbial injury. Nevertheless, to counter life-threatening damage, the lung is as likely to mobilize multiple stem and progenitor cell populations as to evoke both in situ healing and de novo growth. These two repair processes may involve distinct mechanisms where in situ healing is similar to wound healing, while de novo growth forms nascent alveolar lumen, reminiscent of developmental sacculation. Distinguishing these two processes will allow further understanding of recently identified KRT8+ intermediate AT2 cells (Jiang et al., 2020; Kobayashi et al., 2020; Strunz et al., 2020).

The persistence of SOX2+ clusters with minimal differentiation into alveolar cells (Fig. 1C) supports the idea that P63+ basal-like cells repair the airways properly but are resorted to for dysplastic alveolar repair only upon severe injury, as seen in the influenza model as well as COVID-19 patients (Delorey et al., 2021; Kanegai et al., 2016; Kumar et al., 2011; Xi et al., 2017). The resulting ectopic airway differentiation in the alveolar region could form excessive mucus-producing cells or tuft cells (Goldblatt et al., 2020; Rane et al., 2019), contributing to the asthma phenotype independent of a deviated immune environment (Walter et al., 2002). The SOX2+ clusters could also associate with mesenchymal cells of the airways, such as those in the bronchovascular bundles, which might produce more matrix proteins than those of the alveoli, offering a potential link between pulmonary fibrosis and alveolar bronchiolization in fibrotic patient lungs (Cassandras et al., 2020; Dahlgren et al., 2019; Moiseenko et al., 2020; Tsukui et al., 2020). The seemingly irreversible nature of the SOX2+ clusters and associated P63+ cells (Fig. 1, 6) prompts future studies of their responses to repeated injuries and the cumulative consequences of seasonal flu in humans.

## MATERIALS AND METHODS

### Mouse strains and Sendai virus infection

The following mouse strains were used: *Sox2*^*GFP*^ (Arnold et al., 2011), *Sox2*^*CreER*^ (Arnold et al., 2011), *Rosa*^*tdT*^ (Madisen et al., 2010), *Rosa*^*mTmG*^ (Muzumdar et al., 2007), *Sftpc*^*CreER*^ (Rock et al., 2011), and *Rtkn2*^*CreER*^ (Little et al., 2021). Tamoxifen was administered intraperitoneally at least 3 weeks prior to infection at time points and doses were specified in figure legends. Animal infections were carried out as previously described (Cain et al., 2020; Goldblatt et al., 2020). Briefly, mice of at least 8 weeks of age were anaesthetized with isofluorane and suspended by the maxillary incisors on a board with a 60° incline. Mice were infected through oropharyngeal aspiration of a sub-lethal dose of 2.1 × 10^7^ plaque forming units (pfu) of Sendai virus (parainfluenza type 1) strain 52 (ATCC #VR-105, RRID:SCR_001672) in 40 µl PBS. Mice of both genders and mixed genetic background were used. Investigators were not blind to the genotypes. All animal experiments were approved by the Institutional Animal Care and Use at Texas A&M Health Science Center Institute of Biosciences and Technology and MD Anderson Cancer Center.

### Section and wholemount immunostaining

The following antibodies and dilutions were used: rabbit anti-transformation related protein 63 (P63, 1:250, 619002, Biolegend), rabbit anti-red fluorescent protein (tdT, 1:1000, 600-401-379, Rockland), goat anti-sex determining region Y-box 2 (SOX2, 1:250, Santa Cruz Biotechnology, SC-17320), rat anti-antigen identified by monoclonal antibody Ki 67 (Ki67, 1:1000, eBioscience, 14-5698-80), rat anti-cadherin 1 (ECAD, 1:1000, Invitrogen, 131900), chicken anti-green fluorescent protein (GFP, 1:2500, Abcam, ab13970), guinea pig anti-lysosomal-associated membrane protein 3 (LAMP3, 1:500, Synaptic Systems, 391005), rat anti-advanced glycosylation end product-specific receptor (RAGE, 1:1000, R&D Systems, MAB1179), rabbit anti-surfactant associated protein C (SFTPC, 1:1000, Millipore, AB3786), goat anti-podoplanin (PDPN, 1:500, R&D Systems, AF3244), rat anti-protein tyrosine phosphatase, receptor type, C (CD45, 1:1000, eBioscience, 14-0451-81), rabbit anti-NK2 homeobox 1 (NKX2-1, 1:1000, Santa Cruz Biotechnology, sc-13040), rabbit anti-collagen I (COLI, 1:250, Rockland, 600-401-103), and chicken anti-Sendai virus antigen (SeV, 1:5000, Charles River, 10100648).

Lung harvest and immunostaining was carried out as published (Chang et al., 2013; Little et al., 2021). Briefly, Avertin (T48402, Sigma) was used to anesthetize the mice after which the chest cavity was opened for lung perfusion with phosphate buffered saline (PBS). Lungs were gravity inflated with 0.5% paraformaldehyde (PFA; P6148, Sigma) in PBS at 25 cm H_2_O pressure then tied off and fixed in 0.5% PFA over 4-6 hours at room temperature. After washing overnight in PBS at 4°C, lobes were dissected and cryoprotected in a mixture of 20% sucrose and 10% optimal cutting temperature compound (OCT; 4583, Tissue-Tek) in PBS. The lobes were frozen into blocks with OCT compound and stored at 80°C until sectioned. The tissue blocks were sectioned at 10 um thickness and allowed to dry at room temperature for 2 hours. Tissue was circled with a pap pen and after washing three times with PBS was blocked in in PBS with 0.3% Triton X-100 and 5% normal donkey serum (017-000-121, Jackson ImmunoResearch). After blocking, the sections were incubated with primary antibodies overnight in a humidified chamber at 4°C. The slides were washed by submersion in PBS for 30 minutes and incubated with donkey secondary antibodies (Jackson ImmunoResearch) diluted in PBS with 0.3% Triton X-100 plus 4’,6-diamidino-2-phenylindole (DAPI) for 1 hour at room temperature. The slides were washed for another 30 minutes in PBS before being mounted in Aqua-Poly/Mount (18606, Polysciences) and imaged on a Nikon A1plus confocal microscope or an Olympus FV1000 confocal microscope. Cells were quantified manually with Image J.

For wholemount immunostaining, lobes were blocked and incubated with primary and secondary antibodies as above, except for being washed with PBS with 1% Triton X-100 and 1% Tween-20. Fresh or immunostained lobes were imaged with a Leica M205C fluorescence stereomicroscope or an optical projection tomography scanner (OPT, Bioptonics, 3001M) as published (Alanis et al., 2014; Chang et al., 2013).

### Single-cell RNA-seq and data analysis

#### Cell dissociation and labeling

Mice were anaesthetized with Avertin and the lungs perfused as described above. The harvested lungs were minced in PBS and digested in Leibovitz’s L-15 media (Gibco, 21083-027) with 2 mg/mL Collagenase Type I (Worthington, CLS-1, LS004197), 2 mg/mL Elastase (Worthington, ESL, LS002294), and 0.5 mg/mL DNase I (Worthington, D, LS002007) for 30 minutes at 37°C. After the first 15 minutes, the sample was mechanically dissociated through pipetting and then replaced in the heat block to finish the incubation. Fetal bovine serum (FBS; Invitrogen, 10082-139) to a final concentration of 20% and removal of the samples to ice stopped the digestion. The samples were transferred to the cold room and pipetted until no large chunks of tissue were visible then filtered through a 70 μm cell strainer (Falcon, 352350). The cells were centrifuged at 5,000 rpm for 1 minute and resuspended in red blood cell lysis buffer (15 mM NH_4_Cl, 12 mM NaHCO_3_, 0.1 mM EDTA, pH 8.0). After a 3 min incubation, the cells were centrifuged again and washed with media plus 10% FBS. The samples were filtered through a cell strainer cap into a 5 ml tube (Falcon, 352235) and transferred to 2 ml tubes for incubation with the following antibodies diluted in media containing FBS: CD45-PE/Cy7 (BioLegend, 103114), ICAM2-A647 (Life tech, A15452), E-cadherin-A488 (Invitrogen, 53-3249-82) at a concentration of 1:250 for 30 min. The cells were once again pelleted, washed in media with FBS for 5 minutes, resuspended in the fresh media with FBS and filtered through a strainer cap into the final 5 ml tube for FACS.

#### FACS sorting and normalization

A BD FACSAria Fusion Cell Sorter was used for sorting, and SYTOX Blue (Invitrogen, S34857) was added for the exclusion of dead cells. The remaining viable cells were gated using a serial gating strategy as follows: all CD45 positive cells were collected as the immune cell population; CD45 negative cells positive for ICAM2 were collected as endothelial cells; CD45 negative, ICAM2 negative, E-cadherin positive cells were collected as epithelial cells; cells negative for all markers were collected as the mesenchymal population. Each population was collected in a volume of 250 ul PBS with 10% FBS. The 4 cell lineages were normalized as previously described (Cain et al., 2020). Briefly, to determine cell lineage population-specific viability post-sorting, an equal number of cells from each lineage were sorted but collected into the same tube. After adding SYTOX Blue, the mixed population sample was reanalyzed to determine the percentage of viable cells for each lineage. These percentages were used to calculate how much of each of the 4 lineages populations to mix together to achieve as close to equal proportions as possible for scRNA-seq.

#### Data processing

The sorted and normalized lung samples were processed as previously described (Vila Ellis et al., 2020), using the Chromium Single Cell 3’ Library and Gel Bead Kit (v2, rev D). An Illumina NextSeq500 was used for sequencing. The reads were aligned against the mm10 mouse reference genome (provided by 10x Genomics), counted, and aggregated using the Cell Ranger pipeline (version 3.0, 10x Genomics). Single-cell RNA-seq data was processed using Seurat (version 3) following the standard pipeline, as we published for the 14 dpi control and SeV-infected lungs (Cain et al., 2020). The specific gene and cell filters, as well as markers used in naming cell lineages and cell types, were detailed in the R script provided in Supplemental File 1 and shown in Fig. S1. Raw data for the control and SeV-infected lungs were deposited at GEO under accession number GSE144678 and GSE180950.

#### Interactome analysis

The interactome method was published (Cain et al., 2020) and the associated R script was provided in Supplemental File 2. The normalized count data for ligand and receptor genes (gene list from (Ramilowski et al., 2015)) was averaged over all the cells of each cell type. To obtain an interaction strength value for each ligand-receptor pair for each cell type pair, average ligand and receptor gene expression was multiplied. The interquartile ranges (IQR) for the interaction strength values were calculated across all cell type pairs for each ligand-receptor interaction. To be considered outlying, a cell type pair’s interaction strength value had to be above 3 times the IQR. We generated a matrix mapping the locations of the outlying interactions, whether outlying as an enhanced interaction upon SeV infection (1), as a repressed interaction after infection (−1), or not outlying (0).

#### Comparison across time points

The outlier matrices from each of the 3 time points was collapsed into a list and only the information for the cell type-cell type pairs that included AT1 cells as either the signaling or the receiving cell was extracted. We extracted corresponding information from the fold-change matrix to obtain fold-change values and separated time point pairwise comparisons based on whether interaction was outlying for one or both time points, as well as whether outlying between all 3 time points. Colors correspond to time point comparisons.

## ACKNOWLEDGEMENT

The University of Texas MD Anderson Cancer Center DNA Analysis Facility and Flow Cytometry and Cellular Imaging Core Facility are supported by the Cancer Center Support Grant (CA #16672). This work was supported by the University of Texas MD Anderson Cancer Center Start-up and Retention Fund, and National Institutes of Health R01HL130129 and R01HL153511 (JC).

## AUTHOR CONTRIBUTIONS

BJH, MPC, and JC designed and performed research, and wrote the paper; JRF, MJT, and BFD performed research; all authors read and approved the paper.

## REFERENCES

Alanis, D.M., Chang, D.R., Akiyama, H., Krasnow, M.A., and Chen, J. (2014). Two nested developmental waves demarcate a compartment boundary in the mouse lung. Nature communications 5, 3923.

Arnold, K., Sarkar, A., Yram, M.A., Polo, J.M., Bronson, R., Sengupta, S., Seandel, M., Geijsen, N., and Hochedlinger, K. (2011). Sox2(+) adult stem and progenitor cells are important for tissue regeneration and survival of mice. Cell Stem Cell 9, 317–329.

Barkauskas, C.E., Cronce, M.J., Rackley, C.R., Bowie, E.J., Keene, D.R., Stripp, B.R., Randell, S.H., Noble, P.W., and Hogan, B.L. (2013). Type 2 alveolar cells are stem cells in adult lung. J Clin Invest 123, 3025–3036.

Bates, S.R., Tao, J.Q., Collins, H.L., Francone, O.L., and Rothblat, G.H. (2005). Pulmonary abnormalities due to ABCA1 deficiency in mice. Am J Physiol Lung Cell Mol Physiol 289, L980–989.

Cain, M.P., Hernandez, B.J., and Chen, J. (2020). Quantitative single-cell interactomes in normal and virus-infected mouse lungs. Dis Model Mech 13.

Cassandras, M., Wang, C., Kathiriya, J., Tsukui, T., Matatia, P., Matthay, M., Wolters, P., Molofsky, A., Sheppard, D., Chapman, H., et al. (2020). Gli1(+) mesenchymal stromal cells form a pathological niche to promote airway progenitor metaplasia in the fibrotic lung. Nat Cell Biol 22, 1295–1306.

Chang, D.R., Martinez Alanis, D., Miller, R.K., Ji, H., Akiyama, H., McCrea, P.D., and Chen, J. (2013). Lung epithelial branching program antagonizes alveolar differentiation. Proc Natl Acad Sci U S A 110, 18042–18051.

Chen, J. (2017). Origin and regulation of a lung repair kit. Nat Cell Biol 19, 885–886.

Dahlgren, M.W., Jones, S.W., Cautivo, K.M., Dubinin, A., Ortiz-Carpena, J.F., Farhat, S., Yu, K.S., Lee, K., Wang, C., Molofsky, A.V., et al. (2019). Adventitial Stromal Cells Define Group 2 Innate Lymphoid Cell Tissue Niches. Immunity 50, 707–722 e706.

Delorey, T.M., Ziegler, C.G.K., Heimberg, G., Normand, R., Yang, Y., Segerstolpe, A., Abbondanza, D., Fleming, S.J., Subramanian, A., Montoro, D.T., et al. (2021). COVID-19 tissue atlases reveal SARS-CoV-2 pathology and cellular targets. Nature 595, 107–113.

Desai, T.J., Brownfield, D.G., and Krasnow, M.A. (2014). Alveolar progenitor and stem cells in lung development, renewal and cancer. Nature.

Faisca, P., and Desmecht, D. (2007). Sendai virus, the mouse parainfluenza type 1: a longstanding pathogen that remains up-to-date. Res Vet Sci 82, 115–125.

Filipovic, N., Gibney, B.C., Kojic, M., Nikolic, D., Isailovic, V., Ysasi, A., Konerding, M.A., Mentzer, S.J., and Tsuda, A. (2013). Mapping cyclic stretch in the postpneumonectomy murine lung. Journal of applied physiology 115, 1370–1378.

Gahmberg, C.G. (1997). Leukocyte adhesion: CD11/CD18 integrins and intercellular adhesion molecules. Curr Opin Cell Biol 9, 643–650.

Goldblatt, D.L., Flores, J.R., Valverde Ha, G., Jaramillo, A.M., Tkachman, S., Kirkpatrick, C.T., Wali, S., Hernandez, B., Ost, D.E., Scott, B.L., et al. (2020). Inducible epithelial resistance against acute Sendai virus infection prevents chronic asthma-like lung disease in mice. Br J Pharmacol 177, 2256–2273.

Hogan, B.L., Barkauskas, C.E., Chapman, H.A., Epstein, J.A., Jain, R., Hsia, C.C., Niklason, L., Calle, E., Le, A., Randell, S.H., et al. (2014). Repair and regeneration of the respiratory system: complexity, plasticity, and mechanisms of lung stem cell function. Cell Stem Cell 15, 123–138.

Holtzman, M.J., Tyner, J.W., Kim, E.Y., Lo, M.S., Patel, A.C., Shornick, L.P., Agapov, E., and Zhang, Y. (2005). Acute and chronic airway responses to viral infection: implications for asthma and chronic obstructive pulmonary disease. Proc Am Thorac Soc 2, 132–140.

Jiang, P., Gil de Rubio, R., Hrycaj, S.M., Gurczynski, S.J., Riemondy, K.A., Moore, B.B., Omary, M.B., Ridge, K.M., and Zemans, R.L. (2020). Ineffectual Type 2-to-Type 1 Alveolar Epithelial Cell Differentiation in Idiopathic Pulmonary Fibrosis: Persistence of the KRT8(hi) Transitional State. Am J Respir Crit Care Med 201, 1443–1447.

Kanegai, C.M., Xi, Y., Donne, M.L., Gotts, J.E., Driver, I.H., Amidzic, G., Lechner, A.J., Jones, K.D., Vaughan, A.E., Chapman, H.A., et al. (2016). Persistent Pathology in Influenza-Infected Mouse Lungs. Am J Respir Cell Mol Biol 55, 613–615.

Kanner, W.A., Galgano, M.T., and Atkins, K.A. (2010). Podoplanin expression in basal and myoepithelial cells: utility and potential pitfalls. Appl Immunohistochem Mol Morphol 18, 226–230.

Kobayashi, Y., Tata, A., Konkimalla, A., Katsura, H., Lee, R.F., Ou, J., Banovich, N.E., Kropski, J.A., and Tata, P.R. (2020). Persistence of a regeneration-associated, transitional alveolar epithelial cell state in pulmonary fibrosis. Nat Cell Biol 22, 934–946.

Krasna, E., Kolesar, L., Slavcev, A., Valhova, S., Kronosova, B., Jaresova, M., and Striz, I. (2005). IL-18 receptor expression on epithelial cells is upregulated by TNF alpha. Inflammation 29, 33–37.

Kumar, P.A., Hu, Y., Yamamoto, Y., Hoe, N.B., Wei, T.S., Mu, D., Sun, Y., Joo, L.S., Dagher, R., Zielonka, E.M., et al. (2011). Distal airway stem cells yield alveoli in vitro and during lung regeneration following H1N1 influenza infection. Cell 147, 525–538.

Lee, J.H., Bhang, D.H., Beede, A., Huang, T.L., Stripp, B.R., Bloch, K.D., Wagers, A.J., Tseng, Y.H., Ryeom, S., and Kim, C.F. (2014). Lung stem cell differentiation in mice directed by endothelial cells via a BMP4-NFATc1-thrombospondin-1 axis. Cell 156, 440–455.

Lee, J.H., and Rawlins, E.L. (2018). Developmental mechanisms and adult stem cells for therapeutic lung regeneration. Dev Biol 433, 166–176.

Leslie, C.C., McCormick-Shannon, K., Shannon, J.M., Garrick, B., Damm, D., Abraham, J.A., and Mason, R.J. (1997). Heparin-binding EGF-like growth factor is a mitogen for rat alveolar type II cells. Am J Respir Cell Mol Biol 16, 379–387.

Little, D.R., Gerner-Mauro, K.N., Flodby, P., Crandall, E.D., Borok, Z., Akiyama, H., Kimura, S., Ostrin, E.J., and Chen, J. (2019). Transcriptional control of lung alveolar type 1 cell development and maintenance by NK homeobox 2-1. Proc Natl Acad Sci U S A 116, 20545–20555.

Little, D.R., Lynch, A.M., Yan, Y., Akiyama, H., Kimura, S., and Chen, J. (2021). Differential chromatin binding of the lung lineage transcription factor NKX2-1 resolves opposing murine alveolar cell fates in vivo. Nature communications 12, 2509.

Liu, Z., Wu, H., Jiang, K., Wang, Y., Zhang, W., Chu, Q., Li, J., Huang, H., Cai, T., Ji, H., et al. (2016). MAPK-Mediated YAP Activation Controls Mechanical-Tension-Induced Pulmonary Alveolar Regeneration. Cell Rep 16, 1810–1819.

Madisen, L., Zwingman, T.A., Sunkin, S.M., Oh, S.W., Zariwala, H.A., Gu, H., Ng, L.L., Palmiter, R.D., Hawrylycz, M.J., Jones, A.R., et al. (2010). A robust and high-throughput Cre reporting and characterization system for the whole mouse brain. Nat Neurosci 13, 133–140.

Markwell, M.A., and Paulson, J.C. (1980). Sendai virus utilizes specific sialyloligosaccharides as host cell receptor determinants. Proc Natl Acad Sci U S A 77, 5693–5697.

Moiseenko, A., Vazquez-Armendariz, A.I., Kheirollahi, V., Chu, X., Tata, A., Rivetti, S., Gunther, S., Lebrigand, K., Herold, S., Braun, T., et al. (2020). Identification of a Repair-Supportive Mesenchymal Cell Population during Airway Epithelial Regeneration. Cell Rep 33, 108549.

Muzumdar, M.D., Tasic, B., Miyamichi, K., Li, L., and Luo, L. (2007). A global double-fluorescent Cre reporter mouse. Genesis 45, 593–605.

Nabhan, A.N., Brownfield, D.G., Harbury, P.B., Krasnow, M.A., and Desai, T.J. (2018). Single-cell Wnt signaling niches maintain stemness of alveolar type 2 cells. Science 359, 1118–1123.

Ramilowski, J.A., Goldberg, T., Harshbarger, J., Kloppmann, E., Lizio, M., Satagopam, V.P., Itoh, M., Kawaji, H., Carninci, P., Rost, B., et al. (2015). A draft network of ligand-receptor-mediated multicellular signalling in human. Nature communications 6, 7866.

Rane, C.K., Jackson, S.R., Pastore, C.F., Zhao, G., Weiner, A.I., Patel, N.N., Herbert, D.R., Cohen, N.A., and Vaughan, A.E. (2019). Development of solitary chemosensory cells in the distal lung after severe influenza injury. Am J Physiol Lung Cell Mol Physiol 316, L1141–L1149.

Rock, J.R., Barkauskas, C.E., Cronce, M.J., Xue, Y., Harris, J.R., Liang, J., Noble, P.W., and Hogan, B.L. (2011). Multiple stromal populations contribute to pulmonary fibrosis without evidence for epithelial to mesenchymal transition. Proc Natl Acad Sci U S A 108, E1475–1483.

Strunz, M., Simon, L.M., Ansari, M., Kathiriya, J.J., Angelidis, I., Mayr, C.H., Tsidiridis, G., Lange, M., Mattner, L.F., Yee, M., et al. (2020). Alveolar regeneration through a Krt8+ transitional stem cell state that persists in human lung fibrosis. Nature communications 11, 3559.

Tsukui, T., Sun, K.H., Wetter, J.B., Wilson-Kanamori, J.R., Hazelwood, L.A., Henderson, N.C., Adams, T.S., Schupp, J.C., Poli, S.D., Rosas, I.O., et al. (2020). Collagen-producing lung cell atlas identifies multiple subsets with distinct localization and relevance to fibrosis. Nature communications 11, 1920.

Uriarte, J.J., Uhl, F.E., Rolandsson Enes, S.E., Pouliot, R.A., and Weiss, D.J. (2018). Lung bioengineering: advances and challenges in lung decellularization and recellularization. Curr Opin Organ Transplant 23, 673–678.

Vila Ellis, L., Cain, M.P., Hutchison, V., Flodby, P., Crandall, E.D., Borok, Z., Zhou, B., Ostrin, E.J., Wythe, J.D., and Chen, J. (2020). Epithelial Vegfa Specifies a Distinct Endothelial Population in the Mouse Lung. Dev Cell.

Vila Ellis, L., and Chen, J. (2020). A cell-centric view of lung alveologenesis. Dev Dyn.

Walter, M.J., Morton, J.D., Kajiwara, N., Agapov, E., and Holtzman, M.J. (2002). Viral induction of a chronic asthma phenotype and genetic segregation from the acute response. J Clin Invest 110, 165–175.

Weibel, E.R. (2009). What makes a good lung? Swiss Med Wkly 139, 375–386.

Xi, Y., Kim, T., Brumwell, A.N., Driver, I.H., Wei, Y., Tan, V., Jackson, J.R., Xu, J., Lee, D.K., Gotts, J.E., et al. (2017). Local lung hypoxia determines epithelial fate decisions during alveolar regeneration. Nat Cell Biol 19, 904–914.

Yang, J., Hernandez, B.J., Martinez Alanis, D., Narvaez del Pilar, O., Vila-Ellis, L., Akiyama, H., Evans, S.E., Ostrin, E.J., and Chen, J. (2016). The development and plasticity of alveolar type 1 cells. Development 143, 54–65.

Yuan, T., Volckaert, T., Redente, E.F., Hopkins, S., Klinkhammer, K., Wasnick, R., Chao, C.M., Yuan, J., Zhang, J.S., Yao, C., et al. (2019). FGF10-FGFR2B Signaling Generates Basal Cells and Drives Alveolar Epithelial Regeneration by Bronchial Epithelial Stem Cells after Lung Injury. Stem cell reports 12, 1041–1055.

