## Supplemental Figure for "Intermediary role of lung alveolar type 1 cells in epithelial repair upon Sendai virus infection"

**1 Supplemental Figure**

**5 Supplemental Tables**

**2 Supplemental Files**

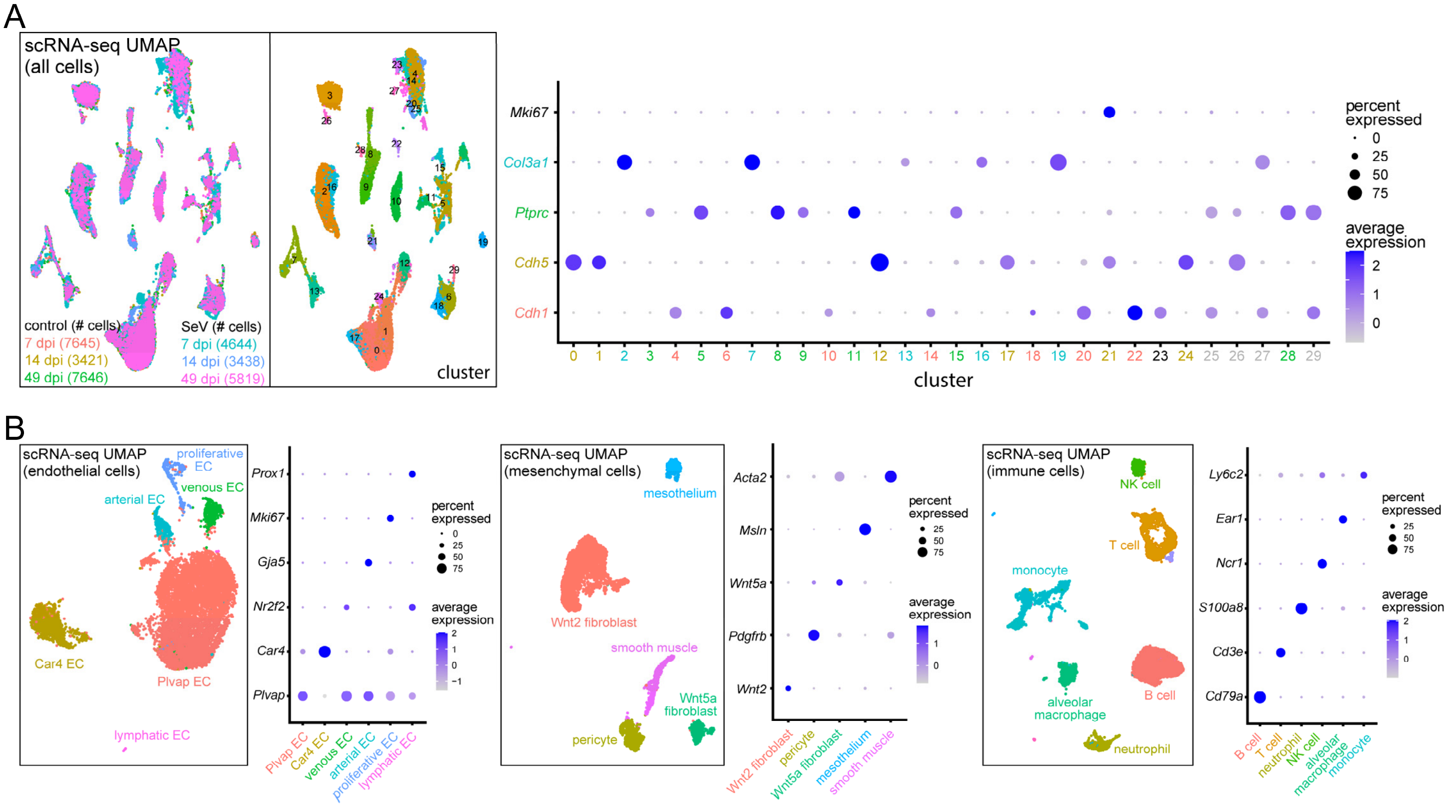


**Figure S1: Identification of cell lineages and cell types in scRNA-seq.**

(**A**) UMAP of all cells from paired control and SeV-infected lungs at 7, 14, and 49 dpi. Dot plot assigns Seurat clusters to color-coded epithelial (*Cdh1*), endothelial (*Cdh5*), immune (*Ptprc*), and mesenchymal (*Col3a1*) lineages. Clusters with multiple lineage markers are considered doublets (grey).

(**B**) UMAPs and dot plots of non-epithelial lineages identify color-coded cell types. The monocyte cluster includes monocytes, dendritic cells, and interstitial macrophages. Immune cell clusters of low abundance are not named.

**Table S1: Differentially expressed genes in AT1 cells at 7 dpi.**

**Table S2: Differentially expressed genes in AT1 cells at 14 dpi.**

**Table S3: Differentially expressed genes in AT1 cells at 49 dpi.**

**Table S4: Ligand-receptor interaction scores in each cell type pair for 7 dpi, 14 dpi, and 49 dpi in control lungs (worksheets 1, 3, 5, respectively) and SeV-infected lungs (worksheets 2, 4, 6, respectively).**

**Table S5: Differences in ligand-receptor interaction in each outlying cell type pair between control and infected lungs and their binary indicators (1, increased; −1, decreased) for each pairwise time point comparison for which the third time point was not outlying (worksheet 1, 7 dpi vs 14 dpi; worksheet 2, 14 dpi vs 49 dpi; worksheet 3, 7 dpi vs 49 dpi).**

**Supplemental File 1: R script for scRNA-seq data processing.**

**Supplemental File 2: R script for interactome analysis.**
