## Supplemental File 1 for "Intermediary role of lung alveolar type 1 cells in epithelial repair upon Sendai virus infection": SeVcellrangerMergeHarmonyAT_virus_DMM.R

```
library(Seurat)
library(cowplot)
library(dplyr)
library(ggplot2)
library(monocle)
library(VGAM)
library(grid)
library(EnhancedVolcano)
library(harmony)


data <- Read10X(data.dir = "Y:/Latitude05262019active/SeV/Sendai1wk2wk7wk-SeVtranscripts/outs/")

integrate <- CreateSeuratObject(data, min.cells = 3, min.features = 200) %>% PercentageFeatureSet(pattern = "^mt-", col.name = "percent.mt") %>% NormalizeData() %>% FindVariableFeatures() %>% ScaleData()
VlnPlot(integrate, features = c("nFeature_RNA","nCount_RNA","percent.mt"))
integrate <- subset(integrate, subset = nFeature_RNA > 200 & nFeature_RNA < 3500 & percent.mt < 15)
integrate$geno <- substr(colnames(integrate@assays$RNA@counts), 18, 18) #cell barcode
integrate$geno <- plyr::mapvalues(x = integrate$geno,
                                            from = 1:6,
                                            to = c("ctrl1wk","ctrl2wk","ctrl7wk","SeV1wk","SeV2wk","SeV7wk"))


integrate <- integrate %>% NormalizeData() %>% FindVariableFeatures() %>% ScaleData() %>% RunPCA(npcs=30) %>% RunHarmony("geno", max.iter.harmony=20) %>% RunUMAP (reduction="harmony", dims=1:30) %>% FindNeighbors(reduction="harmony") %>% FindClusters()
p1 <- DimPlot(integrate, group.by = "geno")
p2 <- DimPlot(integrate, label=T)
plot_grid(p1,p2)
DotPlot(integrate, features = c("Cdh1","Cdh5","Ptprc","Col3a1","Mki67")) + coord_flip()

table(integrate$geno)

rm(data)
saveRDS(integrate, file = "Y:/Latitude05262019active/SeV/Sendai1wk2wk7wk-SeVtranscripts/integrate")
integrate <- readRDS(file = "Y:/Latitude05262019active/SeV/Sendai1wk2wk7wk-SeVtranscripts/integrate")

####include doublets in the initial analysis in case of losing good, proliferative epithelial cells.
integrate_epi <- subset(integrate, idents=c(4,6,10,14,18,20,22,23,25,27,29))
integrate_epi <- integrate_epi %>% NormalizeData() %>% FindVariableFeatures() %>% ScaleData() %>% RunPCA(npcs=30) %>% RunHarmony("geno") %>% RunUMAP (reduction="harmony", dims=1:30) %>% FindNeighbors(reduction="harmony") %>% FindClusters()
p1 <- DimPlot(integrate_epi, group.by = "geno")
p2 <- DimPlot(integrate_epi, label=T)
plot_grid(p1,p2)
DotPlot(integrate_epi, features = c("Cdh1","Cdh5","Ptprc","Col3a1","Mki67")) + coord_flip()

#remove clusters that are all doublets
integrate_epi <- subset(integrate_epi, idents=c(0:6,8:11,15,16))
integrate_epi <- integrate_epi %>% NormalizeData() %>% FindVariableFeatures() %>% ScaleData() %>% RunPCA(npcs=30) %>% RunHarmony("geno") %>% RunUMAP (reduction="harmony", dims=1:30) %>% FindNeighbors(reduction="harmony") %>% FindClusters()
p1 <- DimPlot(integrate_epi, group.by = "geno")
p2 <- DimPlot(integrate_epi, label=T)
plot_grid(p1,p2)

# for figures
DimPlot(integrate_epi, group.by = "seurat_clusters", label=T, repel = T) + scale_y_reverse()
DimPlot(integrate_epi, group.by = "seurat_clusters", split.by = "geno") + NoLegend()
table(integrate_epi$geno)
FeaturePlot(integrate_epi, features=c("Rtkn2","Napsa","Foxj1","Trp63","Scgb1a1","Ifi27l2a","CC.score1","nFeature_RNA"), cols = c('lightgrey','red'), ncol=8)
FeaturePlot(integrate_epi, features=c("Krt8","Lgals3","Sfn","Cdkn1a","Ly6a","H2-K1","H2-Aa","H2-Q7"), cols = c('lightgrey','red'), ncol=8)

integrate_epi$geno_cluster <- paste(integrate_epi$geno, integrate_epi$seurat_clusters, sep="_")
Idents(integrate_epi) <- "geno_cluster"
integrate_epi$cell_type <- integrate_epi$seurat_clusters
integrate_epi$cell_type <- plyr::mapvalues(x = integrate_epi$cell_type,
                                                     from = c(2,5,10,12,13,15,0,1,3,6,7,11,14,4,9,8),
                                                     to = c(rep('AT1',6),rep('AT2',6),'prolifAT2','basal','club','ciliated'))
integrate_epi$geno_cell_type <- paste(integrate_epi$geno, integrate_epi$cell_type, sep="_")
Idents(integrate_epi) <- "geno_cell_type"

diffvol <- FindMarkers(integrate_epi, ident.1 = "SeV7wk_AT1", ident.2 = "ctrl7wk_AT1", min.pct = 0.1, logfc.threshold = 0.1, pseudocount.use = 0.001)
EnhancedVolcano(diffvol, lab = rownames(diffvol), x='avg_logFC', y='p_val_adj', FCcutoff = 1, pCutoff = 10e-6, 
                xlim=c(-max(abs(diffvol$avg_logFC))-.1,max(abs(diffvol$avg_logFC))+.1),
                gridlines.major = F, gridlines.minor = F, xlab = "ln(fold change)", colAlpha = 1,
                title=NULL, subtitle=NULL, col=c('black','black','black','red'), borderWidth = 0.2, vlineWidth = 0.2, hlineWidth = 0.2)
write.csv(diffvol, file = "Y:/Latitude05262019active/SeV/Sendai1wk2wk7wk-SeVtranscripts/AT/diffvol.csv")

EnhancedVolcano(diffvol, lab = rownames(diffvol), x='avg_logFC', y='p_val_adj', FCcutoff = 1, pCutoff = 10e-6, 
                xlim=c(-max(abs(diffvol$avg_logFC))-.1,max(abs(diffvol$avg_logFC))+.1), selectLab=c("Rtkn2","Napsa","Ifi27l2a","H2-K1","H2-Aa","H2-Q6","Cd74","B2m"),
                gridlines.major = F, gridlines.minor = F, xlab = "ln(fold change)", colAlpha = 1,
                title=NULL, subtitle=NULL, col=c('black','black','black','red'), borderWidth = 0.2, vlineWidth = 0.2, hlineWidth = 0.2)


#cell cycle module score. Note that not all human genes are present in mouse genome
firstup <- function(x) {
  substr(x, 1, 1) <- toupper(substr(x, 1, 1))
  x
}
s.genes <- firstup(tolower(cc.genes$s.genes))
g2m.genes <- firstup(tolower(cc.genes$g2m.genes))
tmp <- c(s.genes, g2m.genes)
integrate_epi <- AddModuleScore(integrate_epi, features = list (tmp[tmp %in% integrate_epi@assays[["RNA"]]@counts@Dimnames[[1]]]), name = "CC.score")

saveRDS(integrate_epi, file = "Y:/Latitude05262019active/SeV/Sendai1wk2wk7wk-SeVtranscripts/AT/integrate_epi")
integrate_epi <- readRDS(file = "Y:/Latitude05262019active/SeV/Sendai1wk2wk7wk-SeVtranscripts/AT/integrate_epi")

#### other cell lineages with doublets
integrate_endo <- subset(integrate, idents=c(0,1,12,17,21,24,26))#21 is proliferative
integrate_imm <- subset(integrate, idents=c(3,5,8,9,11,15,21,25,26,28,29))#21 is proliferative
integrate_mes <- subset(integrate, idents=c(2,7,13,16,19,27))

# endothelial cell type
integrate_endo <- integrate_endo %>% NormalizeData() %>% FindVariableFeatures() %>% ScaleData() %>% RunPCA(npcs=30) %>% RunHarmony("geno") %>% RunUMAP (reduction="harmony", dims=1:30) %>% FindNeighbors(reduction="harmony") %>% FindClusters()
integrate_endo <- integrate_endo %>% FindClusters(resolution=2)

integrate_endo <- subset (integrate_endo, idents=c(0:21,24:26))
integrate_endo <- integrate_endo %>% NormalizeData() %>% FindVariableFeatures() %>% ScaleData() %>% RunPCA(npcs=30) %>% RunHarmony("geno") %>% RunUMAP (reduction="harmony", dims=1:30) %>% FindNeighbors(reduction="harmony") %>% FindClusters()

integrate_endo$cell_type <- integrate_endo$seurat_clusters
integrate_endo$cell_type <- plyr::mapvalues(x = integrate_endo$cell_type,
                                           from = c(0:2,4:7,3,11,10,9,8,12),
                                           to = c(rep('Plvap',7),rep('Car4',2),'prolifPlvap','arterial','venous','lymphatics'))
saveRDS(integrate_endo, file = "Y:/Latitude05262019active/SeV/Sendai1wk2wk7wk-SeVtranscripts/AT/integrate_endo")
integrate_endo <- readRDS(file = "Y:/Latitude05262019active/SeV/Sendai1wk2wk7wk-SeVtranscripts/AT/integrate_endo")

Idents(integrate_endo) <- "cell_type"
DimPlot(integrate_endo, group.by = "cell_type", label = F)
DotPlot(integrate_endo, features = c("Plvap","Car4","Nr2f2","Gja5","Mki67","Prox1")) + coord_flip()


# mesenchymal cell type
integrate_mes <- integrate_mes %>% NormalizeData() %>% FindVariableFeatures() %>% ScaleData() %>% RunPCA(npcs=30) %>% RunHarmony("geno") %>% RunUMAP (reduction="harmony", dims=1:30) %>% FindNeighbors(reduction="harmony") %>% FindClusters()

integrate_mes <- subset(integrate_mes, idents=c(0:8,10:12,15,16))
integrate_mes <- integrate_mes %>% NormalizeData() %>% FindVariableFeatures() %>% ScaleData() %>% RunPCA(npcs=30) %>% RunHarmony("geno") %>% RunUMAP (reduction="harmony", dims=1:30) %>% FindNeighbors(reduction="harmony") %>% FindClusters()

integrate_mes$cell_type <- integrate_mes$seurat_clusters
integrate_mes$cell_type <- plyr::mapvalues(x = integrate_mes$cell_type,
                                            from = c(0:2,4,9,10,12,5,13,3,11,7,8,6),
                                            to = c(rep('Interstitial',7),rep('Wnt5',2),rep('pericyte',2),rep('AVSM',2),'mesothelial'))
saveRDS(integrate_mes, file = "Y:/Latitude05262019active/SeV/Sendai1wk2wk7wk-SeVtranscripts/AT/integrate_mes")
integrate_mes <- readRDS(file = "Y:/Latitude05262019active/SeV/Sendai1wk2wk7wk-SeVtranscripts/AT/integrate_mes")

Idents(integrate_mes) <- "cell_type"
DimPlot(integrate_mes, group.by = "cell_type", label = F)
DotPlot(integrate_mes, features = c("Wnt2","Pdgfrb","Wnt5a","Msln","Acta2")) + coord_flip()


# immune cell type
integrate_imm <- integrate_imm %>% NormalizeData() %>% FindVariableFeatures() %>% ScaleData() %>% RunPCA(npcs=30) %>% RunHarmony("geno") %>% RunUMAP (reduction="harmony", dims=1:30) %>% FindNeighbors(reduction="harmony") %>% FindClusters()

integrate_imm <- subset(integrate_imm, idents=c(0:5,7:9,11,15:19))
integrate_imm <- integrate_imm %>% NormalizeData() %>% FindVariableFeatures() %>% ScaleData() %>% RunPCA(npcs=30) %>% RunHarmony("geno") %>% RunUMAP (reduction="harmony", dims=1:30) %>% FindNeighbors(reduction="harmony") %>% FindClusters()
integrate_imm <- integrate_imm %>% FindClusters(resolution=2)
integrate_imm <- subset(integrate_imm, idents=c(0:21,23:25))


integrate_imm$cell_type <- integrate_imm$seurat_clusters
integrate_imm$cell_type <- plyr::mapvalues(x = integrate_imm$cell_type,
                                           from = c(0:2,14,3,4,6,13,7,8,15,20,5,21,9:12,16,17,25),
                                           to = c(rep('B',4),rep('T',4),'NK',rep('AM',3),rep('Neu',2),rep('mono_IM_DC',7)))

saveRDS(integrate_imm, file = "Y:/Latitude05262019active/SeV/Sendai1wk2wk7wk-SeVtranscripts/AT/integrate_imm")
integrate_imm <- readRDS(file = "Y:/Latitude05262019active/SeV/Sendai1wk2wk7wk-SeVtranscripts/AT/integrate_imm")

#integrate_imm <- subset(integrate_imm, idents=c(0:17,20:21,25))
Idents(integrate_imm) <- "cell_type"
DimPlot(integrate_imm, group.by = "cell_type", label = F)
DotPlot(integrate_imm, features = c("Cd79a","Cd3e","S100a8","Ncr1","Ear1","Ly6c2"), idents=c("B","T","Neu","NK","AM","mono_IM_DC")) + coord_flip()
```
