## Supplemental File 2 for "Intermediary role of lung alveolar type 1 cells in epithelial repair upon Sendai virus infection": Step1endo.expression.matrices.R

```
# R code
# Margo P. Cain
# This code will pull expression values for each endothelial cell type that will pass through interactome script in Hernandez et al. 2021

# Load R libs ####
library(Seurat)

# Load Seurat objects ####
integrate.endo <- readRDS("/Users/mpcain/Desktop/Harmony/integrate_endo")

Idents(object = integrate.endo) <- "geno"
ctrl1wk.endo.subset <- subset(integrate.endo, idents = c("ctrl1wk"))
ctrl2wk.endo.subset <- subset(integrate.endo, idents = c("ctrl2wk"))
ctrl7wk.endo.subset <- subset(integrate.endo, idents = c("ctrl7wk"))

SeV1wk.endo.subset <- subset(integrate.endo, idents = c("SeV1wk"))
SeV2wk.endo.subset <- subset(integrate.endo, idents = c("SeV2wk"))
SeV7wk.endo.subset <- subset(integrate.endo, idents = c("SeV7wk"))
#p4.BPD.subset <- readRDS("../p4.BPD.subset.rds")

ctrl.subset <- ctrl2wk.endo.subset
SeV.subset <- SeV2wk.endo.subset

# Load ligand and receptor gene lists from Ramilowski et al.
value_matchReceptors <- as.data.frame(read.csv(file="/Users/mpcain/Documents/scRNAseqData/Visualization/value_matchReceptorsSC40.0810.csv", header=TRUE, sep=","))
value_matchLigands <- as.data.frame(read.csv(file="/Users/mpcain/Documents/scRNAseqData/Visualization/value_matchLigandsSC40.0810.csv", header=TRUE, sep=","))

# Names of ligand and receptor genes from
receptors <- as.vector(value_matchReceptors$uniqueReceptors.uniqueReceptors)
ligands <- as.vector(value_matchLigands$uniqueLigands.uniqueLigands)

# Plvap ####
# control ####

# Extract expression matrix from Seurat object for Plvap cells
Idents(object = ctrl.subset) <- "cell_type"
Plvap.data.ctrl.subset <- subset(ctrl.subset, idents = c("Plvap","prolifPlvap")) #
Plvap.data <- as.matrix(GetAssayData(ctrl.subset, slot = "data")[, WhichCells(ctrl.subset, idents = c("Plvap","prolifPlvap"))])

# Create vector of gene lists from Seurat object
Plvap.data.transpose <- t(Plvap.data)
Plvap.data.df <- as.data.frame(Plvap.data.transpose)
#write.csv(Plvap.data.df, file = "Plvap.data.df.csv")
Plvap_ident <- as.vector(unlist(dimnames(Plvap.data.ctrl.subset)[2]))
barcodes <- Plvap_ident
df_Plvap_ident <- as.character(Plvap.data.ctrl.subset$cell_type)
genelist <- colnames(Plvap.data.df)

# Matches Ramilowski gene names to gene locations in Seurat gene expression matrix
matchedReceptors <- match(receptors, toupper(genelist))
matchedLigands <- match(ligands, toupper(genelist))

# Bind match locations and gene names ####
# Receptors ####
matchRnum <- length(matchedReceptors) # number of receptors that matched to gene list from Seurat matrix
numReceptors <- as.vector(paste(1:matchRnum))
numMatchReceptors <- cbind(as.data.frame(numReceptors), matchedReceptors, receptors)
#null_matchReceptors <- subset(numMatchReceptors, is.na(matchedReceptors)) #
value_matchReceptors_Seurat <- subset(numMatchReceptors, !is.na(matchedReceptors)) #there are 646 receptors

# Ligands ####
matchLnum <- length(matchedLigands)
numLigands <- as.vector(paste(1:matchLnum))
numMatchLigands <- cbind(as.data.frame(numLigands), matchedLigands, ligands)
null_matchLigands <- subset(numMatchLigands, is.na(matchedLigands)) #
value_matchLigands_Seurat <- subset(numMatchLigands, !is.na(matchedLigands)) #there are 647 ligands


# Extract ligand and receptor gene data only for one cell type
Seurat.locations.receptors <- subset(matchedReceptors, !is.na(matchedReceptors))
Seurat.locations.ligands <- subset(matchedLigands, !is.na(matchedLigands))
ReceptGeneExprs <- Plvap.data.df[,Seurat.locations.receptors]
LigGeneExprs <-  Plvap.data.df[,Seurat.locations.ligands]

# Combine Cluster information, Receptor Gene Expression, and Ligand Gene Expression data frames to one data frame
Expression.Table.ctrl.Plvap <- cbind(df_Plvap_ident, LigGeneExprs, ReceptGeneExprs)
write.csv(Expression.Table.ctrl.Plvap, file = "/Users/mpcain/Desktop/Harmony/Expression.Table.ctrl.Plvap.csv")

# SeV ####

# Extract expression matrix from Seurat object for Plvap cells
Idents(object = SeV.subset) <- "cell_type"
Plvap.data.SeV.subset <- subset(SeV.subset, idents = c("Plvap","prolifPlvap")) # 894 cells
Plvap.data <- as.matrix(GetAssayData(SeV.subset, slot = "data")[, WhichCells(SeV.subset, idents = c("Plvap","prolifPlvap"))])

# Create vector of gene lists from Seurat object
Plvap.data.transpose <- t(Plvap.data)
Plvap.data.df <- as.data.frame(Plvap.data.transpose)
#write.csv(Plvap.data.df, file = "Plvap.data.df.csv")
Plvap_ident <- as.vector(unlist(dimnames(Plvap.data.SeV.subset)[2]))
barcodes <- Plvap_ident
df_Plvap_ident <- as.character(Plvap.data.SeV.subset$cell_type)
genelist <- colnames(Plvap.data.df)

# Extract ligand and receptor gene data only for one cell type
Seurat.locations.receptors <- subset(matchedReceptors, !is.na(matchedReceptors))
Seurat.locations.ligands <- subset(matchedLigands, !is.na(matchedLigands))
ReceptGeneExprs <- Plvap.data.df[,Seurat.locations.receptors]
LigGeneExprs <-  Plvap.data.df[,Seurat.locations.ligands]

# Combine Cluster information, Receptor Gene Expression, and Ligand Gene Expression data frames to one data frame
Expression.Table.SeV.Plvap <- cbind(df_Plvap_ident, LigGeneExprs, ReceptGeneExprs)
write.csv(Expression.Table.SeV.Plvap, file = "/Users/mpcain/Desktop/Harmony/Expression.Table.SeV.Plvap.csv")


# Car4 ####
# control ####

# Extract expression matrix from Seurat object for Car4 cells
Idents(object = ctrl.subset) <- "cell_type"
Car4.data.ctrl.subset <- subset(ctrl.subset, idents = c("Car4")) # 338 cells
Car4.data <- as.matrix(GetAssayData(ctrl.subset, slot = "data")[, WhichCells(ctrl.subset, idents = c("Car4"))])

# Create vector of gene lists from Seurat object
Car4.data.transpose <- t(Car4.data)
Car4.data.df <- as.data.frame(Car4.data.transpose)
#write.csv(Car4.data.df, file = "Car4.data.df.csv")
Car4_ident <- as.vector(unlist(dimnames(Car4.data.ctrl.subset)[2]))
barcodes <- Car4_ident
df_Car4_ident <- as.character(Car4.data.ctrl.subset$cell_type)
genelist <- colnames(Car4.data.df)

# Extract ligand and receptor gene data only for one cell type
Seurat.locations.receptors <- subset(matchedReceptors, !is.na(matchedReceptors))
Seurat.locations.ligands <- subset(matchedLigands, !is.na(matchedLigands))
ReceptGeneExprs <- Car4.data.df[,Seurat.locations.receptors]
LigGeneExprs <-  Car4.data.df[,Seurat.locations.ligands]

# Combine Cluster information, Receptor Gene Expression, and Ligand Gene Expression data frames to one data frame
Expression.Table.ctrl.Car4 <- cbind(df_Car4_ident, LigGeneExprs, ReceptGeneExprs)
write.csv(Expression.Table.ctrl.Car4, file = "/Users/mpcain/Desktop/Harmony/Expression.Table.ctrl.Car4.csv")

# SeV ####

# Extract expression matrix from Seurat object for Car4 cells
Idents(object = SeV.subset) <- "cell_type"
Car4.data.SeV.subset <- subset(SeV.subset, idents = c("Car4")) # 456 cells
Car4.data <- as.matrix(GetAssayData(SeV.subset, slot = "data")[, WhichCells(SeV.subset, idents = c("Car4"))])

# Create vector of gene lists from Seurat object
Car4.data.transpose <- t(Car4.data)
Car4.data.df <- as.data.frame(Car4.data.transpose)
#write.csv(Car4.data.df, file = "Car4.data.df.csv")
Car4_ident <- as.vector(unlist(dimnames(Car4.data.SeV.subset)[2]))
barcodes <- Car4_ident
df_Car4_ident <- as.character(Car4.data.SeV.subset$cell_type)
genelist <- colnames(Car4.data.df)

# Extract ligand and receptor gene data only for one cell type
Seurat.locations.receptors <- subset(matchedReceptors, !is.na(matchedReceptors))
Seurat.locations.ligands <- subset(matchedLigands, !is.na(matchedLigands))
ReceptGeneExprs <- Car4.data.df[,Seurat.locations.receptors]
LigGeneExprs <-  Car4.data.df[,Seurat.locations.ligands]

# Combine Cluster information, Receptor Gene Expression, and Ligand Gene Expression data frames to one data frame
Expression.Table.SeV.Car4 <- cbind(df_Car4_ident, LigGeneExprs, ReceptGeneExprs)
write.csv(Expression.Table.SeV.Car4, file = "/Users/mpcain/Desktop/Harmony/Expression.Table.SeV.Car4.csv")


# Vwf ####
# control ####

# Extract expression matrix from Seurat object for Vwf cells
Idents(object = ctrl.subset) <- "cell_type"
Vwf.data.ctrl.subset <- subset(ctrl.subset, idents = c("venous","arterial")) # 292 cells
Vwf.data <- as.matrix(GetAssayData(ctrl.subset, slot = "data")[, WhichCells(ctrl.subset, idents = c("venous","arterial"))])

# Create vector of gene lists from Seurat object
Vwf.data.transpose <- t(Vwf.data)
Vwf.data.df <- as.data.frame(Vwf.data.transpose)
#write.csv(Vwf.data.df, file = "Vwf.data.df.csv")
Vwf_ident <- as.vector(unlist(dimnames(Vwf.data.ctrl.subset)[2]))
barcodes <- Vwf_ident
df_Vwf_ident <- as.character(Vwf.data.ctrl.subset$cell_type)
genelist <- colnames(Vwf.data.df)

# Extract ligand and receptor gene data only for one cell type
Seurat.locations.receptors <- subset(matchedReceptors, !is.na(matchedReceptors))
Seurat.locations.ligands <- subset(matchedLigands, !is.na(matchedLigands))
ReceptGeneExprs <- Vwf.data.df[,Seurat.locations.receptors]
LigGeneExprs <-  Vwf.data.df[,Seurat.locations.ligands]

# Combine Cluster information, Receptor Gene Expression, and Ligand Gene Expression data frames to one data frame
Expression.Table.ctrl.Vwf <- cbind(df_Vwf_ident, LigGeneExprs, ReceptGeneExprs)
write.csv(Expression.Table.ctrl.Vwf, file = "/Users/mpcain/Desktop/Harmony/Expression.Table.ctrl.Vwf.csv")

# SeV ####

# Extract expression matrix from Seurat object for Vwf cells
Idents(object = SeV.subset) <- "cell_type"
Vwf.data.SeV.subset <- subset(SeV.subset, idents = c("venous","arterial")) # 223 cells
Vwf.data <- as.matrix(GetAssayData(SeV.subset, slot = "data")[, WhichCells(SeV.subset, idents = c("venous","arterial"))])

# Create vector of gene lists from Seurat object
Vwf.data.transpose <- t(Vwf.data)
Vwf.data.df <- as.data.frame(Vwf.data.transpose)
#write.csv(Vwf.data.df, file = "Vwf.data.df.csv")
Vwf_ident <- as.vector(unlist(dimnames(Vwf.data.SeV.subset)[2]))
barcodes <- Vwf_ident
df_Vwf_ident <- as.character(Vwf.data.SeV.subset$cell_type)
genelist <- colnames(Vwf.data.df)

# Extract ligand and receptor gene data only for one cell type
Seurat.locations.receptors <- subset(matchedReceptors, !is.na(matchedReceptors))
Seurat.locations.ligands <- subset(matchedLigands, !is.na(matchedLigands))
ReceptGeneExprs <- Vwf.data.df[,Seurat.locations.receptors]
LigGeneExprs <-  Vwf.data.df[,Seurat.locations.ligands]

# Combine Cluster information, Receptor Gene Expression, and Ligand Gene Expression data frames to one data frame
Expression.Table.SeV.Vwf <- cbind(df_Vwf_ident, LigGeneExprs, ReceptGeneExprs)
write.csv(Expression.Table.SeV.Vwf, file = "/Users/mpcain/Desktop/Harmony/Expression.Table.SeV.Vwf.csv")
```
