## Supplemental File 2 for "Intermediary role of lung alveolar type 1 cells in epithelial repair upon Sendai virus infection": Step1epi.expression.matrices.R

```
# R code
# Margo P. Cain
# This code will pull expression values for each epithelial cell type that will pass through interactome script in Hernandez et al. 2021

# Load R libs ####
library(Seurat)

# Load Seurat objects ####

integrate.epi <- readRDS("/Users/mpcain/Desktop/Harmony/integrate_epi")

Idents(object = integrate.epi) <- "geno"
ctrl1wk.subset <- subset(integrate.epi, idents = c("ctrl1wk"))
ctrl2wk.subset <- subset(integrate.epi, idents = c("ctrl2wk"))
ctrl7wk.subset <- subset(integrate.epi, idents = c("ctrl7wk"))

SeV1wk.subset <- subset(integrate.epi, idents = c("SeV1wk"))
SeV2wk.subset <- subset(integrate.epi, idents = c("SeV2wk"))
SeV7wk.subset <- subset(integrate.epi, idents = c("SeV7wk"))

ctrl.subset <- ctrl2wk.subset
SeV.subset <- SeV2wk.subset

# Load ligand and receptor gene lists from Ramilowski et al.
value_matchReceptors <- as.data.frame(read.csv(file="/Users/mpcain/Documents/scRNAseqData/Visualization/value_matchReceptorsSC40.0810.csv", header=TRUE, sep=","))
value_matchLigands <- as.data.frame(read.csv(file="/Users/mpcain/Documents/scRNAseqData/Visualization/value_matchLigandsSC40.0810.csv", header=TRUE, sep=","))

# Names of ligand and receptor genes from
receptors <- as.vector(value_matchReceptors$uniqueReceptors.uniqueReceptors)
ligands <- as.vector(value_matchLigands$uniqueLigands.uniqueLigands)

# Airway ####
# control ####

# Extract expression matrix from Seurat object for airway cells
Idents(object = ctrl.subset) <- "cell_type"
airway.data.ctrl.subset <- subset(ctrl.subset, idents = c("club","ciliated")) #
airway.data <- as.matrix(GetAssayData(ctrl.subset, slot = "data")[, WhichCells(ctrl.subset, idents = c("club","ciliated"))])

# Create vector of gene lists from Seurat object
airway.data.transpose <- t(airway.data)
airway.data.df <- as.data.frame(airway.data.transpose)
#write.csv(airway.data.df, file = "airway.data.df.csv")
airway_ident <- as.vector(unlist(dimnames(airway.data.ctrl.subset)[2]))
barcodes <- airway_ident
df_airway_ident <- as.character(airway.data.ctrl.subset$cell_type)
genelist <- colnames(airway.data.df)

# Matches Ramilowski gene names to gene locations in Seurat gene expression matrix ####
matchedReceptors <- match(receptors, toupper(genelist))
matchedLigands <- match(ligands, toupper(genelist))

# Bind match locations and gene names ####
# Receptors ####
matchRnum <- length(matchedReceptors) # number of receptors that matched to gene list from Seurat matrix
numReceptors <- as.vector(paste(1:matchRnum))
numMatchReceptors <- cbind(as.data.frame(numReceptors), matchedReceptors, receptors)
#null_matchReceptors <- subset(numMatchReceptors, is.na(matchedReceptors)) #
value_matchReceptors_Seurat <- subset(numMatchReceptors, !is.na(matchedReceptors)) #there are 646 receptors

# Ligands ####
matchLnum <- length(matchedLigands)
numLigands <- as.vector(paste(1:matchLnum))
numMatchLigands <- cbind(as.data.frame(numLigands), matchedLigands, ligands)
null_matchLigands <- subset(numMatchLigands, is.na(matchedLigands)) #
value_matchLigands_Seurat <- subset(numMatchLigands, !is.na(matchedLigands)) #there are 647 ligands

# Extract ligand and receptor gene data only for one cell type
Seurat.locations.receptors <- subset(matchedReceptors, !is.na(matchedReceptors))
Seurat.locations.ligands <- subset(matchedLigands, !is.na(matchedLigands))
ReceptGeneExprs <- airway.data.df[,Seurat.locations.receptors]
LigGeneExprs <-  airway.data.df[,Seurat.locations.ligands]

# Combine Cluster information, Receptor Gene Expression, and Ligand Gene Expression data frames to one data frame
Expression.Table.ctrl.airway <- cbind(df_airway_ident, LigGeneExprs, ReceptGeneExprs)
write.csv(Expression.Table.ctrl.airway, file = "/Users/mpcain/Desktop/Harmony/Expression.Table.ctrl.airway.csv")

# SeV ####

# Extract expression matrix from Seurat object for airway cells
Idents(object = SeV.subset) <- "cell_type"
#airway.data.SeV.subset <- subset(SeV.subset, idents = c("club","ciliated")) # no ciliated
#airway.data <- as.matrix(GetAssayData(SeV.subset, slot = "data")[, WhichCells(SeV.subset, idents = c("club","ciliated"))]) # no ciliated
airway.data.SeV.subset <- subset(SeV.subset, idents = c("club")) # no ciliated
airway.data <- as.matrix(GetAssayData(SeV.subset, slot = "data")[, WhichCells(SeV.subset, idents = c("club"))]) # no ciliated


# Create vector of gene lists from Seurat object
airway.data.transpose <- t(airway.data)
airway.data.df <- as.data.frame(airway.data.transpose)
#write.csv(airway.data.df, file = "airway.data.df.csv")
airway_ident <- as.vector(unlist(dimnames(airway.data.SeV.subset)[2]))
barcodes <- airway_ident
df_airway_ident <- as.character(airway.data.SeV.subset$cell_type)
genelist <- colnames(airway.data.df)

# Extract ligand and receptor gene data only for one cell type
Seurat.locations.receptors <- subset(matchedReceptors, !is.na(matchedReceptors))
Seurat.locations.ligands <- subset(matchedLigands, !is.na(matchedLigands))
ReceptGeneExprs <- airway.data.df[,Seurat.locations.receptors]
LigGeneExprs <-  airway.data.df[,Seurat.locations.ligands]

# Combine Cluster information, Receptor Gene Expression, and Ligand Gene Expression data frames to one data frame
Expression.Table.SeV.airway <- cbind(df_airway_ident, LigGeneExprs, ReceptGeneExprs)
write.csv(Expression.Table.SeV.airway, file = "/Users/mpcain/Desktop/Harmony/Expression.Table.SeV.airway.csv")

# AT1 ####
# control ####

# Extract expression matrix from Seurat object for AT1 cells
Idents(object = ctrl.subset) <- "cell_type"
AT1.data.ctrl.subset <- subset(ctrl.subset, idents = c("AT1")) # 1038 cells
AT1.data <- as.matrix(GetAssayData(ctrl.subset, slot = "data")[, WhichCells(ctrl.subset, idents = c("AT1"))])

# Create vector of gene lists from Seurat object
AT1.data.transpose <- t(AT1.data)
AT1.data.df <- as.data.frame(AT1.data.transpose)
#write.csv(AT1.data.df, file = "AT1.data.df.csv")
AT1_ident <- as.vector(unlist(dimnames(AT1.data.ctrl.subset)[2]))
barcodes <- AT1_ident
df_AT1_ident <- as.character(AT1.data.ctrl.subset$cell_type)
genelist <- colnames(AT1.data.df)

# Extract ligand and receptor gene data only for one cell type
Seurat.locations.receptors <- subset(matchedReceptors, !is.na(matchedReceptors))
Seurat.locations.ligands <- subset(matchedLigands, !is.na(matchedLigands))
ReceptGeneExprs <- AT1.data.df[,Seurat.locations.receptors]
LigGeneExprs <-  AT1.data.df[,Seurat.locations.ligands]

# Combine Cluster information, Receptor Gene Expression, and Ligand Gene Expression data frames to one data frame
Expression.Table.ctrl.AT1 <- cbind(df_AT1_ident, LigGeneExprs, ReceptGeneExprs)
write.csv(Expression.Table.ctrl.AT1, file = "/Users/mpcain/Desktop/Harmony/Expression.Table.ctrl.AT1.csv")

# SeV ####

# Extract expression matrix from Seurat object for AT1 cells
Idents(object = SeV.subset) <- "cell_type"
AT1.data.SeV.subset <- subset(SeV.subset, idents = c("AT1")) # 827 cells
AT1.data <- as.matrix(GetAssayData(SeV.subset, slot = "data")[, WhichCells(SeV.subset, idents = c("AT1"))])

# Create vector of gene lists from Seurat object
AT1.data.transpose <- t(AT1.data)
AT1.data.df <- as.data.frame(AT1.data.transpose)
#write.csv(AT1.data.df, file = "AT1.data.df.csv")
AT1_ident <- as.vector(unlist(dimnames(AT1.data.SeV.subset)[2]))
barcodes <- AT1_ident
df_AT1_ident <- as.character(AT1.data.SeV.subset$cell_type)
genelist <- colnames(AT1.data.df)

# Extract ligand and receptor gene data only for one cell type
Seurat.locations.receptors <- subset(matchedReceptors, !is.na(matchedReceptors))
Seurat.locations.ligands <- subset(matchedLigands, !is.na(matchedLigands))
ReceptGeneExprs <- AT1.data.df[,Seurat.locations.receptors]
LigGeneExprs <-  AT1.data.df[,Seurat.locations.ligands]

# Combine Cluster information, Receptor Gene Expression, and Ligand Gene Expression data frames to one data frame
Expression.Table.SeV.AT1 <- cbind(df_AT1_ident, LigGeneExprs, ReceptGeneExprs)
write.csv(Expression.Table.SeV.AT1, file = "/Users/mpcain/Desktop/Harmony/Expression.Table.SeV.AT1.csv")

# AT2 ####
# control ####

# Extract expression matrix from Seurat object for AT2 cells
Idents(object = ctrl.subset) <- "cell_type"
#AT2.data.ctrl.subset <- subset(ctrl.subset, idents = c("AT2","prolifAT2")) # no prolifAT2
#AT2.data <- as.matrix(GetAssayData(ctrl.subset, slot = "data")[, WhichCells(ctrl.subset, idents = c("AT2","prolifAT2"))])
AT2.data.ctrl.subset <- subset(ctrl.subset, idents = c("AT2")) # no prolifAT2
AT2.data <- as.matrix(GetAssayData(ctrl.subset, slot = "data")[, WhichCells(ctrl.subset, idents = c("AT2"))])

# Create vector of gene lists from Seurat object
AT2.data.transpose <- t(AT2.data)
AT2.data.df <- as.data.frame(AT2.data.transpose)
#write.csv(AT2.data.df, file = "AT2.data.df.csv")
AT2_ident <- as.vector(unlist(dimnames(AT2.data.ctrl.subset)[2]))
barcodes <- AT2_ident
df_AT2_ident <- as.character(AT2.data.ctrl.subset$cell_type)
genelist <- colnames(AT2.data.df)

# Extract ligand and receptor gene data only for one cell type
Seurat.locations.receptors <- subset(matchedReceptors, !is.na(matchedReceptors))
Seurat.locations.ligands <- subset(matchedLigands, !is.na(matchedLigands))
ReceptGeneExprs <- AT2.data.df[,Seurat.locations.receptors]
LigGeneExprs <-  AT2.data.df[,Seurat.locations.ligands]

# Combine Cluster information, Receptor Gene Expression, and Ligand Gene Expression data frames to one data frame
Expression.Table.ctrl.AT2 <- cbind(df_AT2_ident, LigGeneExprs, ReceptGeneExprs)
write.csv(Expression.Table.ctrl.AT2, file = "/Users/mpcain/Desktop/Harmony/Expression.Table.ctrl.AT2.csv")

# SeV ####

# Extract expression matrix from Seurat object for AT2 cells
Idents(object = SeV.subset) <- "cell_type"
AT2.data.SeV.subset <- subset(SeV.subset, idents = c("AT2","prolifAT2")) # 984 cells
AT2.data <- as.matrix(GetAssayData(SeV.subset, slot = "data")[, WhichCells(SeV.subset, idents = c("AT2","prolifAT2"))])

# Create vector of gene lists from Seurat object
AT2.data.transpose <- t(AT2.data)
AT2.data.df <- as.data.frame(AT2.data.transpose)
#write.csv(AT2.data.df, file = "AT2.data.df.csv")
AT2_ident <- as.vector(unlist(dimnames(AT2.data.SeV.subset)[2]))
barcodes <- AT2_ident
df_AT2_ident <- as.character(AT2.data.SeV.subset$cell_type)
genelist <- colnames(AT2.data.df)

# Extract ligand and receptor gene data only for one cell type
Seurat.locations.receptors <- subset(matchedReceptors, !is.na(matchedReceptors))
Seurat.locations.ligands <- subset(matchedLigands, !is.na(matchedLigands))
ReceptGeneExprs <- AT2.data.df[,Seurat.locations.receptors]
LigGeneExprs <-  AT2.data.df[,Seurat.locations.ligands]

# Combine Cluster information, Receptor Gene Expression, and Ligand Gene Expression data frames to one data frame
Expression.Table.SeV.AT2 <- cbind(df_AT2_ident, LigGeneExprs, ReceptGeneExprs)
write.csv(Expression.Table.SeV.AT2, file = "/Users/mpcain/Desktop/Harmony/Expression.Table.SeV.AT2.csv")

# Basal ####
# control ####

# Extract expression matrix from Seurat object for basal cells
Idents(object = ctrl.subset) <- "cell_type"
basal.data.ctrl.subset <- subset(ctrl.subset, idents = c("basal")) # 823 cells
basal.data <- as.matrix(GetAssayData(ctrl.subset, slot = "data")[, WhichCells(ctrl.subset, idents = c("basal"))])

# Create vector of gene lists from Seurat object
basal.data.transpose <- t(basal.data)
basal.data.df <- as.data.frame(basal.data.transpose)
#write.csv(basal.data.df, file = "basal.data.df.csv")
basal_ident <- as.vector(unlist(dimnames(basal.data.ctrl.subset)[2]))
barcodes <- basal_ident
df_basal_ident <- as.character(basal.data.ctrl.subset$cell_type)
genelist <- colnames(basal.data.df)

# Extract ligand and receptor gene data only for one cell type
Seurat.locations.receptors <- subset(matchedReceptors, !is.na(matchedReceptors))
Seurat.locations.ligands <- subset(matchedLigands, !is.na(matchedLigands))
ReceptGeneExprs <- basal.data.df[,Seurat.locations.receptors]
LigGeneExprs <-  basal.data.df[,Seurat.locations.ligands]

# Combine Cluster information, Receptor Gene Expression, and Ligand Gene Expression data frames to one data frame
Expression.Table.ctrl.basal <- cbind(df_basal_ident, LigGeneExprs, ReceptGeneExprs)
write.csv(Expression.Table.ctrl.basal, file = "/Users/mpcain/Desktop/Harmony/Expression.Table.ctrl.basal.csv")

# SeV ####

# Extract expression matrix from Seurat object for basal cells
Idents(object = SeV.subset) <- "cell_type"
basal.data.SeV.subset <- subset(SeV.subset, idents = c("basal")) # 984 cells
basal.data <- as.matrix(GetAssayData(SeV.subset, slot = "data")[, WhichCells(SeV.subset, idents = c("basal"))])

# Create vector of gene lists from Seurat object
basal.data.transpose <- t(basal.data)
basal.data.df <- as.data.frame(basal.data.transpose)
#write.csv(basal.data.df, file = "basal.data.df.csv")
basal_ident <- as.vector(unlist(dimnames(basal.data.SeV.subset)[2]))
barcodes <- basal_ident
df_basal_ident <- as.character(basal.data.SeV.subset$cell_type)
genelist <- colnames(basal.data.df)

# Extract ligand and receptor gene data only for one cell type
Seurat.locations.receptors <- subset(matchedReceptors, !is.na(matchedReceptors))
Seurat.locations.ligands <- subset(matchedLigands, !is.na(matchedLigands))
ReceptGeneExprs <- basal.data.df[,Seurat.locations.receptors]
LigGeneExprs <-  basal.data.df[,Seurat.locations.ligands]

# Combine Cluster information, Receptor Gene Expression, and Ligand Gene Expression data frames to one data frame
Expression.Table.SeV.basal <- cbind(df_basal_ident, LigGeneExprs, ReceptGeneExprs)
write.csv(Expression.Table.SeV.basal, file = "/Users/mpcain/Desktop/Harmony/Expression.Table.SeV.basal.csv")
```
