## Supplemental File 2 for "Intermediary role of lung alveolar type 1 cells in epithelial repair upon Sendai virus infection": Step1immune.expression.matrices.R

```
# R code
# Margo P. Cain
# This code will pull expression values for each immune cell type that will pass through interactome script in Hernandez et al. 2021

# Load R libs ####
library(Seurat)

# Load Seurat objects ####
integrate.imm <- readRDS("/Users/mpcain/Desktop/Harmony/integrate_imm")

Idents(object = integrate.imm) <- "geno"
ctrl1wk.imm.subset <- subset(integrate.imm, idents = c("ctrl1wk"))
ctrl2wk.imm.subset <- subset(integrate.imm, idents = c("ctrl2wk"))
ctrl7wk.imm.subset <- subset(integrate.imm, idents = c("ctrl7wk"))

SeV1wk.imm.subset <- subset(integrate.imm, idents = c("SeV1wk"))
SeV2wk.imm.subset <- subset(integrate.imm, idents = c("SeV2wk"))
SeV7wk.imm.subset <- subset(integrate.imm, idents = c("SeV7wk"))
#p4.BPD.subset <- readRDS("../p4.BPD.subset.rds")

ctrl.subset <- ctrl2wk.imm.subset
SeV.subset <- SeV2wk.imm.subset

# Load ligand and receptor gene lists from Ramilowski et al.
value_matchReceptors <- as.data.frame(read.csv(file="/Users/mpcain/Documents/scRNAseqData/Visualization/value_matchReceptorsSC40.0810.csv", header=TRUE, sep=","))
value_matchLigands <- as.data.frame(read.csv(file="/Users/mpcain/Documents/scRNAseqData/Visualization/value_matchLigandsSC40.0810.csv", header=TRUE, sep=","))

# Names of ligand and receptor genes from
receptors <- as.vector(value_matchReceptors$uniqueReceptors.uniqueReceptors)
ligands <- as.vector(value_matchLigands$uniqueLigands.uniqueLigands)

# NK cells ####
# control ####

# Extract expression matrix from Seurat object for NK cells
Idents(object = ctrl.subset) <- "cell_type"
NK.data.ctrl.subset <- subset(ctrl.subset, idents = c("NK")) # 24 cells
NK.data <- as.matrix(GetAssayData(ctrl.subset, slot = "data")[, WhichCells(ctrl.subset, idents = c("NK"))])

# Create vector of gene lists from Seurat object
NK.data.transpose <- t(NK.data)
NK.data.df <- as.data.frame(NK.data.transpose)
#write.csv(NK.data.df, file = "NK.data.df.csv")
NK_ident <- as.vector(unlist(dimnames(NK.data.ctrl.subset)[2]))
barcodes <- NK_ident
df_NK_ident <- as.character(NK.data.ctrl.subset$cell_type)
genelist <- colnames(NK.data.df)

# Matches Ramilowski gene names to gene locations in Seurat gene expression matrix
matchedReceptors <- match(receptors, toupper(genelist))
matchedLigands <- match(ligands, toupper(genelist))

# Bind match locations and gene names ####
# Receptors ####
matchRnum <- length(matchedReceptors) # number of receptors that matched to gene list from Seurat matrix
numReceptors <- as.vector(paste(1:matchRnum))
numMatchReceptors <- cbind(as.data.frame(numReceptors), matchedReceptors, receptors)
#null_matchReceptors <- subset(numMatchReceptors, is.na(matchedReceptors)) #
value_matchReceptors_Seurat <- subset(numMatchReceptors, !is.na(matchedReceptors)) #there are 646 receptors

# Ligands ####
matchLnum <- length(matchedLigands)
numLigands <- as.vector(paste(1:matchLnum))
numMatchLigands <- cbind(as.data.frame(numLigands), matchedLigands, ligands)
null_matchLigands <- subset(numMatchLigands, is.na(matchedLigands)) #
value_matchLigands_Seurat <- subset(numMatchLigands, !is.na(matchedLigands)) #there are 647 ligands

# Extract ligand and receptor gene data only for one cell type
Seurat.locations.receptors <- subset(matchedReceptors, !is.na(matchedReceptors))
Seurat.locations.ligands <- subset(matchedLigands, !is.na(matchedLigands))
ReceptGeneExprs <- NK.data.df[,Seurat.locations.receptors]
LigGeneExprs <-  NK.data.df[,Seurat.locations.ligands]

# Combine Cluster information, Receptor Gene Expression, and Ligand Gene Expression data frames to one data frame
Expression.Table.ctrl.NK <- cbind(df_NK_ident, LigGeneExprs, ReceptGeneExprs)
write.csv(Expression.Table.ctrl.NK, file = "/Users/mpcain/Desktop/Harmony/Expression.Table.ctrl.NK.csv")

# SeV ####

# Extract expression matrix from Seurat object for NK cells
Idents(object = SeV.subset) <- "cell_type"
NK.data.SeV.subset <- subset(SeV.subset, idents = c("NK")) # 25 cells
NK.data <- as.matrix(GetAssayData(SeV.subset, slot = "data")[, WhichCells(SeV.subset, idents = c("NK"))])

# Create vector of gene lists from Seurat object
NK.data.transpose <- t(NK.data)
NK.data.df <- as.data.frame(NK.data.transpose)
#write.csv(NK.data.df, file = "NK.data.df.csv")
NK_ident <- as.vector(unlist(dimnames(NK.data.SeV.subset)[2]))
barcodes <- NK_ident
df_NK_ident <- as.character(NK.data.SeV.subset$cell_type)
genelist <- colnames(NK.data.df)

# Extract ligand and receptor gene data only for one cell type
Seurat.locations.receptors <- subset(matchedReceptors, !is.na(matchedReceptors))
Seurat.locations.ligands <- subset(matchedLigands, !is.na(matchedLigands))
ReceptGeneExprs <- NK.data.df[,Seurat.locations.receptors]
LigGeneExprs <-  NK.data.df[,Seurat.locations.ligands]

# Combine Cluster information, Receptor Gene Expression, and Ligand Gene Expression data frames to one data frame
Expression.Table.SeV.NK <- cbind(df_NK_ident, LigGeneExprs, ReceptGeneExprs)
write.csv(Expression.Table.SeV.NK, file = "/Users/mpcain/Desktop/Harmony/Expression.Table.SeV.NK.csv")


# neutrophils ####
# control ####

# Extract expression matrix from Seurat object for neutrophils
Idents(object = ctrl.subset) <- "cell_type"
neutrophils.data.ctrl.subset <- subset(ctrl.subset, idents = c("Neu")) # 489 cells
neutrophils.data <- as.matrix(GetAssayData(ctrl.subset, slot = "data")[, WhichCells(ctrl.subset, idents = c("Neu"))])

# Create vector of gene lists from Seurat object
neutrophils.data.transpose <- t(neutrophils.data)
neutrophils.data.df <- as.data.frame(neutrophils.data.transpose)
#write.csv(neutrophils.data.df, file = "neutrophils.data.df.csv")
neutrophils_ident <- as.vector(unlist(dimnames(neutrophils.data.ctrl.subset)[2]))
barcodes <- neutrophils_ident
df_neutrophils_ident <- as.character(neutrophils.data.ctrl.subset$cell_type)
genelist <- colnames(neutrophils.data.df)

# Extract ligand and receptor gene data only for one cell type
Seurat.locations.receptors <- subset(matchedReceptors, !is.na(matchedReceptors))
Seurat.locations.ligands <- subset(matchedLigands, !is.na(matchedLigands))
ReceptGeneExprs <- neutrophils.data.df[,Seurat.locations.receptors]
LigGeneExprs <-  neutrophils.data.df[,Seurat.locations.ligands]

# Combine Cluster information, Receptor Gene Expression, and Ligand Gene Expression data frames to one data frame
Expression.Table.ctrl.neutrophils <- cbind(df_neutrophils_ident, LigGeneExprs, ReceptGeneExprs)
write.csv(Expression.Table.ctrl.neutrophils, file = "/Users/mpcain/Desktop/Harmony/Expression.Table.ctrl.neutrophils.csv")

# SeV ####

# Extract expression matrix from Seurat object for neutrophils cells
Idents(object = SeV.subset) <- "cell_type"
neutrophils.data.SeV.subset <- subset(SeV.subset, idents = c("Neu")) # 356 cells
neutrophils.data <- as.matrix(GetAssayData(SeV.subset, slot = "data")[, WhichCells(SeV.subset, idents = c("Neu"))])

# Create vector of gene lists from Seurat object
neutrophils.data.transpose <- t(neutrophils.data)
neutrophils.data.df <- as.data.frame(neutrophils.data.transpose)
#write.csv(neutrophils.data.df, file = "neutrophils.data.df.csv")
neutrophils_ident <- as.vector(unlist(dimnames(neutrophils.data.SeV.subset)[2]))
barcodes <- neutrophils_ident
df_neutrophils_ident <- as.character(neutrophils.data.SeV.subset$cell_type)
genelist <- colnames(neutrophils.data.df)

# Extract ligand and receptor gene data only for one cell type
Seurat.locations.receptors <- subset(matchedReceptors, !is.na(matchedReceptors))
Seurat.locations.ligands <- subset(matchedLigands, !is.na(matchedLigands))
ReceptGeneExprs <- neutrophils.data.df[,Seurat.locations.receptors]
LigGeneExprs <-  neutrophils.data.df[,Seurat.locations.ligands]

# Combine Cluster information, Receptor Gene Expression, and Ligand Gene Expression data frames to one data frame
Expression.Table.SeV.neutrophils <- cbind(df_neutrophils_ident, LigGeneExprs, ReceptGeneExprs)
write.csv(Expression.Table.SeV.neutrophils, file = "/Users/mpcain/Desktop/Harmony/Expression.Table.SeV.neutrophils.csv")

# AM ####
# control ####

# Extract expression matrix from Seurat object for AM
Idents(object = ctrl.subset) <- "cell_type"
AM.data.ctrl.subset <- subset(ctrl.subset, idents = c("AM")) # 177 cells
AM.data <- as.matrix(GetAssayData(ctrl.subset, slot = "data")[, WhichCells(ctrl.subset, idents = c("AM"))])

# Create vector of gene lists from Seurat object
AM.data.transpose <- t(AM.data)
AM.data.df <- as.data.frame(AM.data.transpose)
#write.csv(AM.data.df, file = "AM.data.df.csv")
AM_ident <- as.vector(unlist(dimnames(AM.data.ctrl.subset)[2]))
barcodes <- AM_ident
df_AM_ident <- as.character(AM.data.ctrl.subset$cell_type)
genelist <- colnames(AM.data.df)

# Extract ligand and receptor gene data only for one cell type
Seurat.locations.receptors <- subset(matchedReceptors, !is.na(matchedReceptors))
Seurat.locations.ligands <- subset(matchedLigands, !is.na(matchedLigands))
ReceptGeneExprs <- AM.data.df[,Seurat.locations.receptors]
LigGeneExprs <-  AM.data.df[,Seurat.locations.ligands]

# Combine Cluster information, Receptor Gene Expression, and Ligand Gene Expression data frames to one data frame
Expression.Table.ctrl.AM <- cbind(df_AM_ident, LigGeneExprs, ReceptGeneExprs)
write.csv(Expression.Table.ctrl.AM, file = "/Users/mpcain/Desktop/Harmony/Expression.Table.ctrl.AM.csv")

# SeV ####

# Extract expression matrix from Seurat object for AM cells
Idents(object = SeV.subset) <- "cell_type"
AM.data.SeV.subset <- subset(SeV.subset, idents = c("AM")) # 88 cells
AM.data <- as.matrix(GetAssayData(SeV.subset, slot = "data")[, WhichCells(SeV.subset, idents = c("AM"))])

# Create vector of gene lists from Seurat object
AM.data.transpose <- t(AM.data)
AM.data.df <- as.data.frame(AM.data.transpose)
#write.csv(AM.data.df, file = "AM.data.df.csv")
AM_ident <- as.vector(unlist(dimnames(AM.data.SeV.subset)[2]))
barcodes <- AM_ident
df_AM_ident <- as.character(AM.data.SeV.subset$cell_type)
genelist <- colnames(AM.data.df)

# Extract ligand and receptor gene data only for one cell type
Seurat.locations.receptors <- subset(matchedReceptors, !is.na(matchedReceptors))
Seurat.locations.ligands <- subset(matchedLigands, !is.na(matchedLigands))
ReceptGeneExprs <- AM.data.df[,Seurat.locations.receptors]
LigGeneExprs <-  AM.data.df[,Seurat.locations.ligands]

# Combine Cluster information, Receptor Gene Expression, and Ligand Gene Expression data frames to one data frame
Expression.Table.SeV.AM <- cbind(df_AM_ident, LigGeneExprs, ReceptGeneExprs)
write.csv(Expression.Table.SeV.AM, file = "/Users/mpcain/Desktop/Harmony/Expression.Table.SeV.AM.csv")


# B cells ####
# control ####

# Extract expression matrix from Seurat object for B cells
Idents(object = ctrl.subset) <- "cell_type"
B.data.ctrl.subset <- subset(ctrl.subset, idents = c("B")) # 285 cells
B.data <- as.matrix(GetAssayData(ctrl.subset, slot = "data")[, WhichCells(ctrl.subset, idents = c("B"))])

# Create vector of gene lists from Seurat object
B.data.transpose <- t(B.data)
B.data.df <- as.data.frame(B.data.transpose)
#write.csv(B.data.df, file = "B.data.df.csv")
B_ident <- as.vector(unlist(dimnames(B.data.ctrl.subset)[2]))
barcodes <- B_ident
df_B_ident <- as.character(B.data.ctrl.subset$cell_type)
genelist <- colnames(B.data.df)

# Extract ligand and receptor gene data only for one cell type
Seurat.locations.receptors <- subset(matchedReceptors, !is.na(matchedReceptors))
Seurat.locations.ligands <- subset(matchedLigands, !is.na(matchedLigands))
ReceptGeneExprs <- B.data.df[,Seurat.locations.receptors]
LigGeneExprs <-  B.data.df[,Seurat.locations.ligands]

# Combine Cluster information, Receptor Gene Expression, and Ligand Gene Expression data frames to one data frame
Expression.Table.ctrl.B <- cbind(df_B_ident, LigGeneExprs, ReceptGeneExprs)
write.csv(Expression.Table.ctrl.B, file = "/Users/mpcain/Desktop/Harmony/Expression.Table.ctrl.B.csv")

# SeV ####

# Extract expression matrix from Seurat object for B cells
Idents(object = SeV.subset) <- "cell_type"
B.data.SeV.subset <- subset(SeV.subset, idents = c("B")) # 330 cells
B.data <- as.matrix(GetAssayData(SeV.subset, slot = "data")[, WhichCells(SeV.subset, idents = c("B"))])

# Create vector of gene lists from Seurat object
B.data.transpose <- t(B.data)
B.data.df <- as.data.frame(B.data.transpose)
#write.csv(B.data.df, file = "B.data.df.csv")
B_ident <- as.vector(unlist(dimnames(B.data.SeV.subset)[2]))
barcodes <- B_ident
df_B_ident <- as.character(B.data.SeV.subset$cell_type)
genelist <- colnames(B.data.df)

# Extract ligand and receptor gene data only for one cell type
Seurat.locations.receptors <- subset(matchedReceptors, !is.na(matchedReceptors))
Seurat.locations.ligands <- subset(matchedLigands, !is.na(matchedLigands))
ReceptGeneExprs <- B.data.df[,Seurat.locations.receptors]
LigGeneExprs <-  B.data.df[,Seurat.locations.ligands]

# Combine Cluster information, Receptor Gene Expression, and Ligand Gene Expression data frames to one data frame
Expression.Table.SeV.B <- cbind(df_B_ident, LigGeneExprs, ReceptGeneExprs)
write.csv(Expression.Table.SeV.B, file = "/Users/mpcain/Desktop/Harmony/Expression.Table.SeV.B.csv")


# T cells ####
# control ####

# Extract expression matrix from Seurat object for T cells
Idents(object = ctrl.subset) <- "cell_type"
T.data.ctrl.subset <- subset(ctrl.subset, idents = c("T")) # 79 cells
T.data <- as.matrix(GetAssayData(ctrl.subset, slot = "data")[, WhichCells(ctrl.subset, idents = c("T"))])

# Create vector of gene lists from Seurat object
T.data.transpose <- t(T.data)
T.data.df <- as.data.frame(T.data.transpose)
#write.csv(T.data.df, file = "T.data.df.csv")
T_ident <- as.vector(unlist(dimnames(T.data.ctrl.subset)[2]))
barcodes <- T_ident
df_T_ident <- as.character(T.data.ctrl.subset$cell_type)
genelist <- colnames(T.data.df)

# Extract ligand and receptor gene data only for one cell type
Seurat.locations.receptors <- subset(matchedReceptors, !is.na(matchedReceptors))
Seurat.locations.ligands <- subset(matchedLigands, !is.na(matchedLigands))
ReceptGeneExprs <- T.data.df[,Seurat.locations.receptors]
LigGeneExprs <-  T.data.df[,Seurat.locations.ligands]

# Combine Cluster information, Receptor Gene Expression, and Ligand Gene Expression data frames to one data frame
Expression.Table.ctrl.T <- cbind(df_T_ident, LigGeneExprs, ReceptGeneExprs)
write.csv(Expression.Table.ctrl.T, file = "/Users/mpcain/Desktop/Harmony/Expression.Table.ctrl.T.csv")

# SeV ####

# Extract expression matrix from Seurat object for T cells
Idents(object = SeV.subset) <- "cell_type"
T.data.SeV.subset <- subset(SeV.subset, idents = c("T")) # 61 cells
T.data <- as.matrix(GetAssayData(SeV.subset, slot = "data")[, WhichCells(SeV.subset, idents = c("T"))])

# Create vector of gene lists from Seurat object
T.data.transpose <- t(T.data)
T.data.df <- as.data.frame(T.data.transpose)
#write.csv(T.data.df, file = "T.data.df.csv")
T_ident <- as.vector(unlist(dimnames(T.data.SeV.subset)[2]))
barcodes <- T_ident
df_T_ident <- as.character(T.data.SeV.subset$cell_type)
genelist <- colnames(T.data.df)

# Extract ligand and receptor gene data only for one cell type
Seurat.locations.receptors <- subset(matchedReceptors, !is.na(matchedReceptors))
Seurat.locations.ligands <- subset(matchedLigands, !is.na(matchedLigands))
ReceptGeneExprs <- T.data.df[,Seurat.locations.receptors]
LigGeneExprs <-  T.data.df[,Seurat.locations.ligands]

# Combine Cluster information, Receptor Gene Expression, and Ligand Gene Expression data frames to one data frame
Expression.Table.SeV.T <- cbind(df_T_ident, LigGeneExprs, ReceptGeneExprs)
write.csv(Expression.Table.SeV.T, file = "/Users/mpcain/Desktop/Harmony/Expression.Table.SeV.T.csv")

# mono.IM.DC ####
# control ####

# Extract expression matrix from Seurat object for IM and DC
Idents(object = ctrl.subset) <- "cell_type"
IM.data.ctrl.subset <- subset(ctrl.subset, idents = c("mono_IM_DC")) # 181 cells
IM.data <- as.matrix(GetAssayData(ctrl.subset, slot = "data")[, WhichCells(ctrl.subset, idents = c("mono_IM_DC"))])

# Create vector of gene lists from Seurat object
IM.data.transpose <- t(IM.data)
IM.data.df <- as.data.frame(IM.data.transpose)
#write.csv(IM.data.df, file = "IM.data.df.csv")
IM_ident <- as.vector(unlist(dimnames(IM.data.ctrl.subset)[2]))
barcodes <- IM_ident
df_IM_ident <- as.character(IM.data.ctrl.subset$cell_type)
genelist <- colnames(IM.data.df)

# Extract ligand and receptor gene data only for one cell type
Seurat.locations.receptors <- subset(matchedReceptors, !is.na(matchedReceptors))
Seurat.locations.ligands <- subset(matchedLigands, !is.na(matchedLigands))
ReceptGeneExprs <- IM.data.df[,Seurat.locations.receptors]
LigGeneExprs <-  IM.data.df[,Seurat.locations.ligands]

# Combine Cluster information, Receptor Gene Expression, and Ligand Gene Expression data frames to one data frame
Expression.Table.ctrl.IM.DC <- cbind(df_IM_ident, LigGeneExprs, ReceptGeneExprs)
write.csv(Expression.Table.ctrl.IM.DC, file = "/Users/mpcain/Desktop/Harmony/Expression.Table.ctrl.mono.IM.DC.csv")

# SeV ####

# Extract expression matrix from Seurat object for IM and DC
Idents(object = SeV.subset) <- "cell_type"
IM.data.SeV.subset <- subset(SeV.subset, idents = c("mono_IM_DC")) # 238 cells
IM.data <- as.matrix(GetAssayData(SeV.subset, slot = "data")[, WhichCells(SeV.subset, idents = c("mono_IM_DC"))])

# Create vector of gene lists from Seurat object
IM.data.transpose <- t(IM.data)
IM.data.df <- as.data.frame(IM.data.transpose)
#write.csv(IM.data.df, file = "IM.data.df.csv")
IM_ident <- as.vector(unlist(dimnames(IM.data.SeV.subset)[2]))
barcodes <- IM_ident
df_IM_ident <- as.character(IM.data.SeV.subset$cell_type)
genelist <- colnames(IM.data.df)

# Extract ligand and receptor gene data only for one cell type
Seurat.locations.receptors <- subset(matchedReceptors, !is.na(matchedReceptors))
Seurat.locations.ligands <- subset(matchedLigands, !is.na(matchedLigands))
ReceptGeneExprs <- IM.data.df[,Seurat.locations.receptors]
LigGeneExprs <-  IM.data.df[,Seurat.locations.ligands]

# Combine Cluster information, Receptor Gene Expression, and Ligand Gene Expression data frames to one data frame
Expression.Table.SeV.IM.DC <- cbind(df_IM_ident, LigGeneExprs, ReceptGeneExprs)
write.csv(Expression.Table.SeV.IM.DC, file = "/Users/mpcain/Desktop/Harmony/Expression.Table.SeV.mono.IM.DC.csv")
```
