## Supplemental File 2 for "Intermediary role of lung alveolar type 1 cells in epithelial repair upon Sendai virus infection": Step1mes.expression.matrices.R

```
# R code
# Margo P. Cain
# This code will pull expression values for each mesenchymal cell type that will pass through interactome script in Hernandez et al. 2021

# Load R libs ####
library(Seurat)

# Load Seurat objects ####
integrate.mes <- readRDS("/Users/mpcain/Desktop/Harmony/integrate_mes")

Idents(object = integrate.mes) <- "geno"
ctrl1wk.mes.subset <- subset(integrate.mes, idents = c("ctrl1wk"))
ctrl2wk.mes.subset <- subset(integrate.mes, idents = c("ctrl2wk"))
ctrl7wk.mes.subset <- subset(integrate.mes, idents = c("ctrl7wk"))

SeV1wk.mes.subset <- subset(integrate.mes, idents = c("SeV1wk"))
SeV2wk.mes.subset <- subset(integrate.mes, idents = c("SeV2wk"))
SeV7wk.mes.subset <- subset(integrate.mes, idents = c("SeV7wk"))

ctrl.subset <- ctrl1wk.mes.subset
SeV.subset <- SeV1wk.mes.subset

# Load ligand and receptor gene lists from Ramilowski et al.
value_matchReceptors <- as.data.frame(read.csv(file="/Users/mpcain/Documents/scRNAseqData/Visualization/value_matchReceptorsSC40.0810.csv", header=TRUE, sep=","))
value_matchLigands <- as.data.frame(read.csv(file="/Users/mpcain/Documents/scRNAseqData/Visualization/value_matchLigandsSC40.0810.csv", header=TRUE, sep=","))

# Names of ligand and receptor genes from
receptors <- as.vector(value_matchReceptors$uniqueReceptors.uniqueReceptors)
ligands <- as.vector(value_matchLigands$uniqueLigands.uniqueLigands)

# interstitial fibroblasts ####
# control ####

# Extract expression matrix from Seurat object for interstitial cells
Idents(object = ctrl.subset) <- "cell_type"
interstitial.data.ctrl.subset <- subset(ctrl.subset, idents = c("Interstitial")) # 1549 cells
interstitial.data <- as.matrix(GetAssayData(ctrl.subset, slot = "data")[, WhichCells(ctrl.subset, idents = c("Interstitial"))])

# Create vector of gene lists from Seurat object
interstitial.data.transpose <- t(interstitial.data)
interstitial.data.df <- as.data.frame(interstitial.data.transpose)
#write.csv(interstitial.data.df, file = "interstitial.data.df.csv")
interstitial_ident <- as.vector(unlist(dimnames(interstitial.data.ctrl.subset)[2]))
barcodes <- interstitial_ident
df_interstitial_ident <- as.character(interstitial.data.ctrl.subset$cell_type)
genelist <- colnames(interstitial.data.df)

# Matches Ramilowski gene names to gene locations in Seurat gene expression matrix
matchedReceptors <- match(receptors, toupper(genelist))
matchedLigands <- match(ligands, toupper(genelist))

# Bind match locations and gene names ####
# Receptors ####
matchRnum <- length(matchedReceptors) # number of receptors that matched to gene list from Seurat matrix
numReceptors <- as.vector(paste(1:matchRnum))
numMatchReceptors <- cbind(as.data.frame(numReceptors), matchedReceptors, receptors)
#null_matchReceptors <- subset(numMatchReceptors, is.na(matchedReceptors)) #
value_matchReceptors_Seurat <- subset(numMatchReceptors, !is.na(matchedReceptors)) #there are 646 receptors

# Ligands ####
matchLnum <- length(matchedLigands)
numLigands <- as.vector(paste(1:matchLnum))
numMatchLigands <- cbind(as.data.frame(numLigands), matchedLigands, ligands)
null_matchLigands <- subset(numMatchLigands, is.na(matchedLigands)) #
value_matchLigands_Seurat <- subset(numMatchLigands, !is.na(matchedLigands)) #there are 647 ligands

# Extract ligand and receptor gene data only for one cell type
Seurat.locations.receptors <- subset(matchedReceptors, !is.na(matchedReceptors))
Seurat.locations.ligands <- subset(matchedLigands, !is.na(matchedLigands))
ReceptGeneExprs <- interstitial.data.df[,Seurat.locations.receptors]
LigGeneExprs <-  interstitial.data.df[,Seurat.locations.ligands]

# Combine Cluster information, Receptor Gene Expression, and Ligand Gene Expression data frames to one data frame
Expression.Table.ctrl.interstitial <- cbind(df_interstitial_ident, LigGeneExprs, ReceptGeneExprs)
write.csv(Expression.Table.ctrl.interstitial, file = "/Users/mpcain/Desktop/Harmony.1wk/Expression.Table.ctrl.interstitial.csv")

# SeV ####

# Extract expression matrix from Seurat object for interstitial cells
Idents(object = SeV.subset) <- "cell_type"
interstitial.data.SeV.subset <- subset(SeV.subset, idents = c("Interstitial")) # 1443 cells
interstitial.data <- as.matrix(GetAssayData(SeV.subset, slot = "data")[, WhichCells(SeV.subset, idents = c("Interstitial"))])

# Create vector of gene lists from Seurat object
interstitial.data.transpose <- t(interstitial.data)
interstitial.data.df <- as.data.frame(interstitial.data.transpose)
#write.csv(interstitial.data.df, file = "interstitial.data.df.csv")
interstitial_ident <- as.vector(unlist(dimnames(interstitial.data.SeV.subset)[2]))
barcodes <- interstitial_ident
df_interstitial_ident <- as.character(interstitial.data.SeV.subset$cell_type)
genelist <- colnames(interstitial.data.df)

# Extract ligand and receptor gene data only for one cell type
Seurat.locations.receptors <- subset(matchedReceptors, !is.na(matchedReceptors))
Seurat.locations.ligands <- subset(matchedLigands, !is.na(matchedLigands))
ReceptGeneExprs <- interstitial.data.df[,Seurat.locations.receptors]
LigGeneExprs <-  interstitial.data.df[,Seurat.locations.ligands]

# Combine Cluster information, Receptor Gene Expression, and Ligand Gene Expression data frames to one data frame
Expression.Table.SeV.interstitial <- cbind(df_interstitial_ident, LigGeneExprs, ReceptGeneExprs)
write.csv(Expression.Table.SeV.interstitial, file = "/Users/mpcain/Desktop/Harmony.1wk/Expression.Table.SeV.interstitial.csv")


# Wnt5 ####
# control ####

# Extract expression matrix from Seurat object for Wnt5 cells
Idents(object = ctrl.subset) <- "cell_type"
Wnt5.data.ctrl.subset <- subset(ctrl.subset, idents = c("Wnt5")) # 470 cells
Wnt5.data <- as.matrix(GetAssayData(ctrl.subset, slot = "data")[, WhichCells(ctrl.subset, idents = c("Wnt5"))])

# Create vector of gene lists from Seurat object
Wnt5.data.transpose <- t(Wnt5.data)
Wnt5.data.df <- as.data.frame(Wnt5.data.transpose)
#write.csv(Wnt5.data.df, file = "Wnt5.data.df.csv")
Wnt5_ident <- as.vector(unlist(dimnames(Wnt5.data.ctrl.subset)[2]))
barcodes <- Wnt5_ident
df_Wnt5_ident <- as.character(Wnt5.data.ctrl.subset$cell_type)
genelist <- colnames(Wnt5.data.df)

# Extract ligand and receptor gene data only for one cell type
Seurat.locations.receptors <- subset(matchedReceptors, !is.na(matchedReceptors))
Seurat.locations.ligands <- subset(matchedLigands, !is.na(matchedLigands))
ReceptGeneExprs <- Wnt5.data.df[,Seurat.locations.receptors]
LigGeneExprs <-  Wnt5.data.df[,Seurat.locations.ligands]

# Combine Cluster information, Receptor Gene Expression, and Ligand Gene Expression data frames to one data frame
Expression.Table.ctrl.Wnt5 <- cbind(df_Wnt5_ident, LigGeneExprs, ReceptGeneExprs)
write.csv(Expression.Table.ctrl.Wnt5, file = "/Users/mpcain/Desktop/Harmony.1wk/Expression.Table.ctrl.Wnt5.csv")

# SeV ####

# Extract expression matrix from Seurat object for Wnt5 cells
Idents(object = SeV.subset) <- "cell_type"
Wnt5.data.SeV.subset <- subset(SeV.subset, idents = c("Wnt5")) # 542 cells
Wnt5.data <- as.matrix(GetAssayData(SeV.subset, slot = "data")[, WhichCells(SeV.subset, idents = c("Wnt5"))])

# Create vector of gene lists from Seurat object
Wnt5.data.transpose <- t(Wnt5.data)
Wnt5.data.df <- as.data.frame(Wnt5.data.transpose)
#write.csv(Wnt5.data.df, file = "Wnt5.data.df.csv")
Wnt5_ident <- as.vector(unlist(dimnames(Wnt5.data.SeV.subset)[2]))
barcodes <- Wnt5_ident
df_Wnt5_ident <- as.character(Wnt5.data.SeV.subset$cell_type)
genelist <- colnames(Wnt5.data.df)

# Extract ligand and receptor gene data only for one cell type
Seurat.locations.receptors <- subset(matchedReceptors, !is.na(matchedReceptors))
Seurat.locations.ligands <- subset(matchedLigands, !is.na(matchedLigands))
ReceptGeneExprs <- Wnt5.data.df[,Seurat.locations.receptors]
LigGeneExprs <-  Wnt5.data.df[,Seurat.locations.ligands]

# Combine Cluster information, Receptor Gene Expression, and Ligand Gene Expression data frames to one data frame
Expression.Table.SeV.Wnt5 <- cbind(df_Wnt5_ident, LigGeneExprs, ReceptGeneExprs)
write.csv(Expression.Table.SeV.Wnt5, file = "/Users/mpcain/Desktop/Harmony.1wk/Expression.Table.SeV.Wnt5.csv")

# pericyte ####
# control ####

# Extract expression matrix from Seurat object for pericyte
Idents(object = ctrl.subset) <- "cell_type"
pericyte.data.ctrl.subset <- subset(ctrl.subset, idents = c("pericyte")) # 296 cells
pericyte.data <- as.matrix(GetAssayData(ctrl.subset, slot = "data")[, WhichCells(ctrl.subset, idents = c("pericyte"))])

# Create vector of gene lists from Seurat object
pericyte.data.transpose <- t(pericyte.data)
pericyte.data.df <- as.data.frame(pericyte.data.transpose)
#write.csv(pericyte.data.df, file = "pericyte.data.df.csv")
pericyte_ident <- as.vector(unlist(dimnames(pericyte.data.ctrl.subset)[2]))
barcodes <- pericyte_ident
df_pericyte_ident <- as.character(pericyte.data.ctrl.subset$cell_type)
genelist <- colnames(pericyte.data.df)

# Extract ligand and receptor gene data only for one cell type
Seurat.locations.receptors <- subset(matchedReceptors, !is.na(matchedReceptors))
Seurat.locations.ligands <- subset(matchedLigands, !is.na(matchedLigands))
ReceptGeneExprs <- pericyte.data.df[,Seurat.locations.receptors]
LigGeneExprs <-  pericyte.data.df[,Seurat.locations.ligands]

# Combine Cluster information, Receptor Gene Expression, and Ligand Gene Expression data frames to one data frame
Expression.Table.ctrl.pericyte <- cbind(df_pericyte_ident, LigGeneExprs, ReceptGeneExprs)
write.csv(Expression.Table.ctrl.pericyte, file = "/Users/mpcain/Desktop/Harmony.1wk/Expression.Table.ctrl.pericyte.csv")

# SeV ####

# Extract expression matrix from Seurat object for pericyte cells
Idents(object = SeV.subset) <- "cell_type"
pericyte.data.SeV.subset <- subset(SeV.subset, idents = c("pericyte")) # 140 cells
pericyte.data <- as.matrix(GetAssayData(SeV.subset, slot = "data")[, WhichCells(SeV.subset, idents = c("pericyte"))])

# Create vector of gene lists from Seurat object
pericyte.data.transpose <- t(pericyte.data)
pericyte.data.df <- as.data.frame(pericyte.data.transpose)
#write.csv(pericyte.data.df, file = "pericyte.data.df.csv")
pericyte_ident <- as.vector(unlist(dimnames(pericyte.data.SeV.subset)[2]))
barcodes <- pericyte_ident
df_pericyte_ident <- as.character(pericyte.data.SeV.subset$cell_type)
genelist <- colnames(pericyte.data.df)

# Extract ligand and receptor gene data only for one cell type
Seurat.locations.receptors <- subset(matchedReceptors, !is.na(matchedReceptors))
Seurat.locations.ligands <- subset(matchedLigands, !is.na(matchedLigands))
ReceptGeneExprs <- pericyte.data.df[,Seurat.locations.receptors]
LigGeneExprs <-  pericyte.data.df[,Seurat.locations.ligands]

# Combine Cluster information, Receptor Gene Expression, and Ligand Gene Expression data frames to one data frame
Expression.Table.SeV.pericyte <- cbind(df_pericyte_ident, LigGeneExprs, ReceptGeneExprs)
write.csv(Expression.Table.SeV.pericyte, file = "/Users/mpcain/Desktop/Harmony.1wk/Expression.Table.SeV.pericyte.csv")


# AVSM ####
# control ####

# Extract expression matrix from Seurat object for AVSM
Idents(object = ctrl.subset) <- "cell_type"
AVSM.data.ctrl.subset <- subset(ctrl.subset, idents = c("AVSM")) # 142 cells
AVSM.data <- as.matrix(GetAssayData(ctrl.subset, slot = "data")[, WhichCells(ctrl.subset, idents = c("AVSM"))])

# Create vector of gene lists from Seurat object
AVSM.data.transpose <- t(AVSM.data)
AVSM.data.df <- as.data.frame(AVSM.data.transpose)
#write.csv(AVSM.data.df, file = "AVSM.data.df.csv")
AVSM_ident <- as.vector(unlist(dimnames(AVSM.data.ctrl.subset)[2]))
barcodes <- AVSM_ident
df_AVSM_ident <- as.character(AVSM.data.ctrl.subset$cell_type)
genelist <- colnames(AVSM.data.df)

# Extract ligand and receptor gene data only for one cell type
Seurat.locations.receptors <- subset(matchedReceptors, !is.na(matchedReceptors))
Seurat.locations.ligands <- subset(matchedLigands, !is.na(matchedLigands))
ReceptGeneExprs <- AVSM.data.df[,Seurat.locations.receptors]
LigGeneExprs <-  AVSM.data.df[,Seurat.locations.ligands]

# Combine Cluster information, Receptor Gene Expression, and Ligand Gene Expression data frames to one data frame
Expression.Table.ctrl.AVSM <- cbind(df_AVSM_ident, LigGeneExprs, ReceptGeneExprs)
write.csv(Expression.Table.ctrl.AVSM, file = "/Users/mpcain/Desktop/Harmony.1wk/Expression.Table.ctrl.AVSM.csv")

# SeV ####

# Extract expression matrix from Seurat object for AVSM cells
Idents(object = SeV.subset) <- "cell_type"
AVSM.data.SeV.subset <- subset(SeV.subset, idents = c("AVSM")) # 163 cells
AVSM.data <- as.matrix(GetAssayData(SeV.subset, slot = "data")[, WhichCells(SeV.subset, idents = c("AVSM"))])

# Create vector of gene lists from Seurat object
AVSM.data.transpose <- t(AVSM.data)
AVSM.data.df <- as.data.frame(AVSM.data.transpose)
#write.csv(AVSM.data.df, file = "AVSM.data.df.csv")
AVSM_ident <- as.vector(unlist(dimnames(AVSM.data.SeV.subset)[2]))
barcodes <- AVSM_ident
df_AVSM_ident <- as.character(AVSM.data.SeV.subset$cell_type)
genelist <- colnames(AVSM.data.df)

# Extract ligand and receptor gene data only for one cell type
Seurat.locations.receptors <- subset(matchedReceptors, !is.na(matchedReceptors))
Seurat.locations.ligands <- subset(matchedLigands, !is.na(matchedLigands))
ReceptGeneExprs <- AVSM.data.df[,Seurat.locations.receptors]
LigGeneExprs <-  AVSM.data.df[,Seurat.locations.ligands]

# Combine Cluster information, Receptor Gene Expression, and Ligand Gene Expression data frames to one data frame
Expression.Table.SeV.AVSM <- cbind(df_AVSM_ident, LigGeneExprs, ReceptGeneExprs)
write.csv(Expression.Table.SeV.AVSM, file = "/Users/mpcain/Desktop/Harmony.1wk/Expression.Table.SeV.AVSM.csv")


# mesothelial ####
# control ####

# Extract expression matrix from Seurat object for mesothelial
Idents(object = ctrl.subset) <- "cell_type"
mesothelial.data.ctrl.subset <- subset(ctrl.subset, idents = c("mesothelial")) # 108 cells
mesothelial.data <- as.matrix(GetAssayData(ctrl.subset, slot = "data")[, WhichCells(ctrl.subset, idents = c("mesothelial"))])

# Create vector of gene lists from Seurat object
mesothelial.data.transpose <- t(mesothelial.data)
mesothelial.data.df <- as.data.frame(mesothelial.data.transpose)
#write.csv(mesothelial.data.df, file = "mesothelial.data.df.csv")
mesothelial_ident <- as.vector(unlist(dimnames(mesothelial.data.ctrl.subset)[2]))
barcodes <- mesothelial_ident
df_mesothelial_ident <- as.character(mesothelial.data.ctrl.subset$cell_type)
genelist <- colnames(mesothelial.data.df)

# Extract ligand and receptor gene data only for one cell type
Seurat.locations.receptors <- subset(matchedReceptors, !is.na(matchedReceptors))
Seurat.locations.ligands <- subset(matchedLigands, !is.na(matchedLigands))
ReceptGeneExprs <- mesothelial.data.df[,Seurat.locations.receptors]
LigGeneExprs <-  mesothelial.data.df[,Seurat.locations.ligands]

# Combine Cluster information, Receptor Gene Expression, and Ligand Gene Expression data frames to one data frame
Expression.Table.ctrl.mesothelial <- cbind(df_mesothelial_ident, LigGeneExprs, ReceptGeneExprs)
write.csv(Expression.Table.ctrl.mesothelial, file = "/Users/mpcain/Desktop/Harmony.1wk/Expression.Table.ctrl.mesothelial.csv")

# SeV ####

# Extract expression matrix from Seurat object for mesothelial cells
Idents(object = SeV.subset) <- "cell_type"
mesothelial.data.SeV.subset <- subset(SeV.subset, idents = c("mesothelial")) # 147 cells
mesothelial.data <- as.matrix(GetAssayData(SeV.subset, slot = "data")[, WhichCells(SeV.subset, idents = c("mesothelial"))])

# Create vector of gene lists from Seurat object
mesothelial.data.transpose <- t(mesothelial.data)
mesothelial.data.df <- as.data.frame(mesothelial.data.transpose)
#write.csv(mesothelial.data.df, file = "mesothelial.data.df.csv")
mesothelial_ident <- as.vector(unlist(dimnames(mesothelial.data.SeV.subset)[2]))
barcodes <- mesothelial_ident
df_mesothelial_ident <- as.character(mesothelial.data.SeV.subset$cell_type)
genelist <- colnames(mesothelial.data.df)

# Extract ligand and receptor gene data only for one cell type
Seurat.locations.receptors <- subset(matchedReceptors, !is.na(matchedReceptors))
Seurat.locations.ligands <- subset(matchedLigands, !is.na(matchedLigands))
ReceptGeneExprs <- mesothelial.data.df[,Seurat.locations.receptors]
LigGeneExprs <-  mesothelial.data.df[,Seurat.locations.ligands]

# Combine Cluster information, Receptor Gene Expression, and Ligand Gene Expression data frames to one data frame
Expression.Table.SeV.mesothelial <- cbind(df_mesothelial_ident, LigGeneExprs, ReceptGeneExprs)
write.csv(Expression.Table.SeV.mesothelial, file = "/Users/mpcain/Desktop/Harmony.1wk/Expression.Table.SeV.mesothelial.csv")
```
