## Supplemental File 2 for "Intermediary role of lung alveolar type 1 cells in epithelial repair upon Sendai virus infection": Step2.interactome.R

```
# R code
# Margo P. Cain
# This code will generate the interactome used in Hernandez et al. Figure 6
sessionInfo

# Load R libs ####
library(reshape2)
library(ggplot2)

# Load expression tables for each cell of each cell types, control and Sendai-infected (indicated by 'mut')  ####
Expression.Table.CtrlApln <- as.data.frame(read.csv(file="/Users/mpcain/Desktop/Harmony/Expression.Table.ctrl.Car4.csv", header=TRUE, sep=","))
Expression.Table.CtrlBulkEndo <- as.data.frame(read.csv(file="/Users/mpcain/Desktop/Harmony/Expression.Table.ctrl.Plvap.csv", header=TRUE, sep=","))
Expression.Table.CtrlMatureVessels <- as.data.frame(read.csv(file="/Users/mpcain/Desktop/Harmony/Expression.Table.ctrl.Vwf.csv", header=TRUE, sep=","))

Expression.Table.MutApln <- as.data.frame(read.csv(file="/Users/mpcain/Desktop/Harmony/Expression.Table.SeV.Car4.csv", header=TRUE, sep=","))
Expression.Table.MutBulkEndo <- as.data.frame(read.csv(file="/Users/mpcain/Desktop/Harmony/Expression.Table.SeV.Plvap.csv", header=TRUE, sep=","))
Expression.Table.MutMatureVessels <- as.data.frame(read.csv(file="/Users/mpcain/Desktop/Harmony/Expression.Table.SeV.Vwf.csv", header=TRUE, sep=","))

Expression.Table.CtrlInterstitial <- as.data.frame(read.csv(file="/Users/mpcain/Desktop/Harmony/Expression.Table.ctrl.interstitial.csv", header=TRUE, sep=","))
Expression.Table.CtrlMeso <- as.data.frame(read.csv(file="/Users/mpcain/Desktop/Harmony/Expression.Table.ctrl.mesothelial.csv", header=TRUE, sep=","))
Expression.Table.CtrlPeri <- as.data.frame(read.csv(file="/Users/mpcain/Desktop/Harmony/Expression.Table.ctrl.pericyte.csv", header=TRUE, sep=","))
Expression.Table.CtrlWnt5 <- as.data.frame(read.csv(file="/Users/mpcain/Desktop/Harmony/Expression.Table.ctrl.Wnt5.csv", header=TRUE, sep=","))
Expression.Table.CtrlAVSM <- as.data.frame(read.csv(file="/Users/mpcain/Desktop/Harmony/Expression.Table.ctrl.AVSM.csv", header=TRUE, sep=","))

Expression.Table.MutInterstitial <- as.data.frame(read.csv(file="/Users/mpcain/Desktop/Harmony/Expression.Table.SeV.interstitial.csv", header=TRUE, sep=","))
Expression.Table.MutMeso <- as.data.frame(read.csv(file="/Users/mpcain/Desktop/Harmony/Expression.Table.SeV.mesothelial.csv", header=TRUE, sep=","))
Expression.Table.MutPeri <- as.data.frame(read.csv(file="/Users/mpcain/Desktop/Harmony/Expression.Table.SeV.pericyte.csv", header=TRUE, sep=","))
Expression.Table.MutWnt5 <- as.data.frame(read.csv(file="/Users/mpcain/Desktop/Harmony/Expression.Table.SeV.Wnt5.csv", header=TRUE, sep=","))
Expression.Table.MutAVSM <- as.data.frame(read.csv(file="/Users/mpcain/Desktop/Harmony/Expression.Table.SeV.AVSM.csv", header=TRUE, sep=","))

Expression.Table.CtrlAT1 <- as.data.frame(read.csv(file="/Users/mpcain/Desktop/Harmony/Expression.Table.ctrl.AT1.csv", header=TRUE, sep=","))
Expression.Table.CtrlAT2 <- as.data.frame(read.csv(file="/Users/mpcain/Desktop/Harmony/Expression.Table.ctrl.AT2.csv", header=TRUE, sep=","))
Expression.Table.CtrlAirway <- as.data.frame(read.csv(file="/Users/mpcain/Desktop/Harmony/Expression.Table.ctrl.airway.csv", header=TRUE, sep=","))
Expression.Table.CtrlBasal <- as.data.frame(read.csv(file="/Users/mpcain/Desktop/Harmony/Expression.Table.ctrl.basal.csv", header=TRUE, sep=","))

Expression.Table.MutAT1 <- as.data.frame(read.csv(file="/Users/mpcain/Desktop/Harmony/Expression.Table.SeV.AT1.csv", header=TRUE, sep=","))
Expression.Table.MutAT2 <- as.data.frame(read.csv(file="/Users/mpcain/Desktop/Harmony/Expression.Table.SeV.AT2.csv", header=TRUE, sep=","))
Expression.Table.MutAirway <- as.data.frame(read.csv(file="/Users/mpcain/Desktop/Harmony/Expression.Table.SeV.airway.csv", header=TRUE, sep=","))
Expression.Table.MutBasal <- as.data.frame(read.csv(file="/Users/mpcain/Desktop/Harmony/Expression.Table.SeV.basal.csv", header=TRUE, sep=","))

Expression.Table.CtrlT <- as.data.frame(read.csv(file="/Users/mpcain/Desktop/Harmony/Expression.Table.ctrl.T.csv", header=TRUE, sep=","))
Expression.Table.CtrlAM <- as.data.frame(read.csv(file="/Users/mpcain/Desktop/Harmony/Expression.Table.ctrl.AM.csv", header=TRUE, sep=","))
Expression.Table.CtrlNeu <- as.data.frame(read.csv(file="/Users/mpcain/Desktop/Harmony/Expression.Table.ctrl.neutrophils.csv", header=TRUE, sep=","))
Expression.Table.CtrlB <- as.data.frame(read.csv(file="/Users/mpcain/Desktop/Harmony/Expression.Table.ctrl.B.csv", header=TRUE, sep=","))
Expression.Table.CtrlNK <- as.data.frame(read.csv(file="/Users/mpcain/Desktop/Harmony/Expression.Table.ctrl.NK.csv", header=TRUE, sep=","))
Expression.Table.CtrlMono.IM.DC <- as.data.frame(read.csv(file="/Users/mpcain/Desktop/Harmony/Expression.Table.ctrl.mono.IM.DC.csv", header=TRUE, sep=","))

Expression.Table.MutT <- as.data.frame(read.csv(file="/Users/mpcain/Desktop/Harmony/Expression.Table.SeV.T.csv", header=TRUE, sep=","))
Expression.Table.MutAM <- as.data.frame(read.csv(file="/Users/mpcain/Desktop/Harmony/Expression.Table.SeV.AM.csv", header=TRUE, sep=","))
Expression.Table.MutNeu <- as.data.frame(read.csv(file="/Users/mpcain/Desktop/Harmony/Expression.Table.SeV.neutrophils.csv", header=TRUE, sep=","))
Expression.Table.MutB <- as.data.frame(read.csv(file="/Users/mpcain/Desktop/Harmony/Expression.Table.SeV.B.csv", header=TRUE, sep=","))
Expression.Table.MutNK <- as.data.frame(read.csv(file="/Users/mpcain/Desktop/Harmony/Expression.Table.SeV.NK.csv", header=TRUE, sep=","))
Expression.Table.MutMono.IM.DC <- as.data.frame(read.csv(file="/Users/mpcain/Desktop/Harmony/Expression.Table.SeV.mono.IM.DC.csv", header=TRUE, sep=","))


# Populate a list with the data for each cell type to make it easier to average the gene expression data for each cell type ####
Expr.Table.List <- list()
             
Expr.Table.List[[1]] <- Expression.Table.CtrlApln
Expr.Table.List[[2]] <- Expression.Table.CtrlBulkEndo
Expr.Table.List[[3]] <- Expression.Table.CtrlMatureVessels

Expr.Table.List[[4]] <- Expression.Table.MutApln
Expr.Table.List[[5]] <- Expression.Table.MutBulkEndo
Expr.Table.List[[6]] <- Expression.Table.MutMatureVessels

Expr.Table.List[[7]] <- Expression.Table.CtrlInterstitial
Expr.Table.List[[8]] <- Expression.Table.CtrlMeso
Expr.Table.List[[9]] <- Expression.Table.CtrlPeri
Expr.Table.List[[10]] <- Expression.Table.CtrlWnt5
Expr.Table.List[[11]] <- Expression.Table.CtrlAVSM

Expr.Table.List[[12]] <- Expression.Table.MutInterstitial
Expr.Table.List[[13]] <- Expression.Table.MutMeso
Expr.Table.List[[14]] <- Expression.Table.MutPeri
Expr.Table.List[[15]] <- Expression.Table.MutWnt5
Expr.Table.List[[16]] <- Expression.Table.MutAVSM

Expr.Table.List[[17]] <- Expression.Table.CtrlAT1
Expr.Table.List[[18]] <- Expression.Table.CtrlAT2
Expr.Table.List[[19]] <- Expression.Table.CtrlAirway
Expr.Table.List[[20]] <- Expression.Table.CtrlBasal

Expr.Table.List[[21]] <- Expression.Table.MutAT1
Expr.Table.List[[22]] <- Expression.Table.MutAT2
Expr.Table.List[[23]] <- Expression.Table.MutAirway
Expr.Table.List[[24]] <- Expression.Table.MutBasal

Expr.Table.List[[25]] <- Expression.Table.CtrlT
Expr.Table.List[[26]] <- Expression.Table.CtrlAM
Expr.Table.List[[27]] <- Expression.Table.CtrlNeu
Expr.Table.List[[28]] <- Expression.Table.CtrlB
Expr.Table.List[[29]] <- Expression.Table.CtrlNK
Expr.Table.List[[30]] <- Expression.Table.CtrlMono.IM.DC

Expr.Table.List[[31]] <- Expression.Table.MutT
Expr.Table.List[[32]] <- Expression.Table.MutAM
Expr.Table.List[[33]] <- Expression.Table.MutNeu
Expr.Table.List[[34]] <- Expression.Table.MutB
Expr.Table.List[[35]] <- Expression.Table.MutNK
Expr.Table.List[[36]] <- Expression.Table.MutMono.IM.DC

# Subset for each cell type the expression data only ####
Expr.Tbl.length <- length(Expression.Table.CtrlApln)
subset_genes <- function(Expr.Table.List, x){
  v <- as.data.frame(Expr.Table.List[x])
  list_df2_genes_only <- rbind(as.vector(v[,3:Expr.Tbl.length]))
}
list_df2_genes_only <- lapply(Expr.Table.List, subset_genes) # runs the above function on the list

# Average expression for each across all cell per cell type in the list ####
cluster_means <- function(list_df2_genes_only, x){
  v <- as.data.frame(list_df2_genes_only[x])
  Cmeans <- rbind(as.vector(colMeans(v)))
}
Cmeans <- lapply(list_df2_genes_only, cluster_means) # runs the above function on the list

#export Cmeans as excel file
Cmeans2 <- do.call(rbind.data.frame, Cmeans) # combines column means for each cell type into one data frame of all the cell types
LRnames <- names(Expression.Table.CtrlApln[3:Expr.Tbl.length])
colnames(Cmeans2) <- LRnames
rownames(Cmeans2) <- c("CtrlApln", "CtrlBulkEndo","CtrlMatureVessels","MutApln","MutBulkEndo","MutMatureVessels","CtrlInterstitial", "CtrlMeso", "CtrlPeri", "CtrlWnt5", "CtrlAVSM", "MutInterstitial", "MutMeso", "MutPeri", "MutWnt5", "MutAVSM", "CtrlAT1", "CtrlAT2", "CtrlAirway","CtrlBasal", "MutAT1", "MutAT2", "MutAirway","MutBasal", "CtrlT", "CtrlAM", "CtrlNeu", "CtrlB","CtrlNK","CtrlMono.IM.DC", "MutT", "MutAM", "MutNeu","MutB","MutNK","MutMono.IM.DC")
write.csv(Cmeans2, file = "/Users/mpcain/Desktop/Harmony/SeV.harmony.LRavgExpressionMeansPerCellType2020.csv")

###
SeV.2wk.Cmeans <- as.data.frame(read.csv(file="/Users/mpcain/Desktop/Harmony/SeV.harmony.LRavgExpressionMeansPerCellType2020.csv", header=TRUE, sep=","))
LRnames <- as.character(colnames(SeV.2wk.Cmeans)) #gene names of all ligand and receptor genes from column means csv
cellTypesNum <- as.numeric(nrow(SeV.2wk.Cmeans)) #number of cell types from column means csv

#ligand-receptor pair key list from Ramilowski et al.
LRpairs <- read.csv(file="/Users/mpcain/Desktop/Interactome.Manuscript/RamilowskiAllCondensed02082018mm.expressmatch_edit08112019.csv", header=TRUE, sep=",")

# Separate control and infected cell types into two different objects ####
List.Ctrl <- list()
ctrlnums <- c(1:3,7:11,17:20,25:30)
List.Ctrl <- SeV.2wk.Cmeans[ctrlnums,]
write.csv(List.Ctrl, file="/Users/mpcain/Desktop/Harmony/List.SeV.2wk.Ctrl2020.csv")

List.Mut <- list()
mutnums <- c(4:6,12:16,21:24,31:36)
List.Mut <- SeV.2wk.Cmeans[mutnums,]
write.csv(List.Mut, file="/Users/mpcain/Desktop/Harmony/List.SeV.2wk.Mut2020.csv")

# Need to separate ligand and receptor genes into different lists now
genesnum <- as.numeric(length(colnames(List.Ctrl))) #number of genes/columns in column means

DimPlot(integrate.imm, reduction = "umap",group.by = 'cell_type', label = TRUE)

colnames(List.Ctrl)[537] #end of ligands
colnames(List.Ctrl)[538] #beginning of receptors

List.Ctrl.ligands <- List.Ctrl[2:537] #skips first column, which contains cell type names, then takes all ligand gene columns
List.Mut.ligands <- List.Mut[2:537] #same for infected as done above with control

List.Ctrl.receptors <- List.Ctrl[538:genesnum] #takes all receptors
List.Mut.receptors <- List.Mut[538:genesnum] #same as above

# ligands are already in alphabetical order but receptors now need to be put in the order that matches Ligand-receptor pair list
# This will be the correct L-R pair order needed to reporder receptor genes in Control and Infected receptor data lists
#Extracts receptor names from L-R list
LRpairs.receptors <- LRpairs$Receptor.ApprovedSymbol

# Gets rid of receptor duplicate names
uniqueLRpairsReceptors <- unique(LRpairs.receptors)

# match function to find locations where Seurat order of (lower case) receptor genes (from Cmeans) match unique receptor names from Ramilowski L-R list (in upper case)
Cmeans.receptor.genenames <- colnames(List.Ctrl)[538:genesnum]
Cmeans.receptor.genenames
LRpair.receptor.order.locations <- match(uniqueLRpairsReceptors, toupper(Cmeans.receptor.genenames)) #

# Reordering cmeans receptor gene/column names to same order as Ramilowski using match locations
LRnames.recept.order <- Cmeans.receptor.genenames[LRpair.receptor.order.locations]

# you manually can look at this data frame below to see if both columns match - they should
receptor.match.df <- as.data.frame(uniqueLRpairsReceptors,LRnames.recept.order) # match but several are missing

# need to get rid of NAs
LRpair.receptor.order.locations <- LRpair.receptor.order.locations[which(!is.na(LRpair.receptor.order.locations))]
LRnames.recept.order <- Cmeans.receptor.genenames[LRpair.receptor.order.locations]

#uniqueLRpairsReceptors2 <- uniqueLRpairsReceptors[which(!is.na(LRpair.receptor.order.locations))] # maybe doesn't matter

#receptor.match.df2 <- as.data.frame(uniqueLRpairsReceptors2,LRnames.recept.order) # maybe doesn't matter

# Reorder control and mutant receptors list to correct order for making LR pairs
List.Ctrl.receptors.reorder <- List.Ctrl.receptors[,LRpair.receptor.order.locations]
List.Mut.receptors.reorder <- List.Mut.receptors[,LRpair.receptor.order.locations]

# Vectors in which to store the locations for each receptor and ligand to use when matching up L-R values
ligandsSeuratordered <- toupper(colnames(List.Ctrl.ligands))
receptorsSeuratordered <- toupper(colnames(List.Ctrl.receptors.reorder))

# must still have NA genes...will need to just get rid of them??? bc LRpair.ligand.order.locations.final

LRpairs.ligands <- as.character(LRpairs$Ligand.ApprovedSymbol)
LRpairs.receptors <- as.character(LRpairs$Receptor.ApprovedSymbol)

LRpair.ligand.order.locations.final <- match(LRpairs.ligands,ligandsSeuratordered) #correct
LRpair.receptor.order.locations.final <- match(LRpairs.receptors,receptorsSeuratordered) #correct
LRpair.ligand.order.locations.final
LRpair.receptor.order.locations.final

#need to create variables with the number of cell types and number of L-R pairs for loops below
cellTypeslength <- nrow(List.Ctrl.ligands)
LRpair.num <- nrow(LRpairs)

### Makes correct order of x and y to make plots in 18 x 18 matrix
# For X, want to repeat cell types 1 through 18
plotOrderX <- rep.int(1:cellTypeslength, cellTypeslength)
plotOrderX

# For y, want to repeat cell type 1 eighteen times, then cell type 2 eighteen times, etc so that we get each cell-cell combination
cellTypesnumlist <- 1:cellTypeslength
plotOrderY <- rep(1:cellTypeslength, each = cellTypeslength)
plotOrderY
Plotsnum <- length(plotOrderY) # this is how many cell-cell combinations there are

# creating empty matrices to fill using loop below
# X and Y alone for control and mutant form Figure 3 plots, while CtrlXnYn and MutXnYn will form Figure 4 plots
# CONTROL ####
CtrlXn <- data.frame(matrix(nrow = LRpair.num, ncol = cellTypeslength))
CtrlYn <- data.frame(matrix(nrow = LRpair.num, ncol = cellTypeslength))
CtrlXnYn <- data.frame(matrix(nrow = LRpair.num, ncol = cellTypeslength))

# Makes all x cooordinates for every plot in plot matrix. Each column is the ligands' expressions, in order of LRpairs list, for one cell type
colnames(LRpairs)[1] <- paste("Pair.Name")
for (i in 1:Plotsnum){
  m <- plotOrderX[i]
  v <- as.data.frame(unlist(List.Ctrl.ligands[m,]))
  CtrlXn[i] <- v[LRpair.ligand.order.locations.final,]
  rownames(CtrlXn) <- LRpairs$Pair.Name
}  

# Makes all y cooordinates for every plot in plot matrix. Each column is the receptors' expressions, in order of LRpairs list, for one cell type
for (j in 1:Plotsnum){
  l <- plotOrderY[j]
  t <- as.data.frame(unlist(List.Ctrl.receptors.reorder[l,]))
  
  CtrlYn[j] <- t[LRpair.receptor.order.locations.final,]
  rownames(CtrlYn) <- LRpairs$Pair.Name
}

# Makes all cooordinates for every plot in plot matrix for visualizationplot3. each column is the product of the ligands' and receptors' expressions, in order of LRpairs list, for one cell type
for (i in 1:Plotsnum){
  m <- plotOrderX[i]
  l <- plotOrderY[i]
  CtrlXnYn[i] <- CtrlXn[,i]*CtrlYn[,i]
  rownames(CtrlXnYn) <- LRpairs$Pair.Name
}

### Makes correct order of x and y to make plots in 18 x 18 matrix - this time using cell type labels
# For X, want to repeat cell types 1 through 18
plotOrderLigandCells <- rep.int(List.Ctrl[,1], cellTypeslength)
plotOrderLigandCells

# For y, want to repeat cell type 1 eighteen times, then cell type 2 eighteen times, etc so that we get each cell-cell combination
cellTypesnumlist <- 1:cellTypeslength
plotOrderReceptorCells <- rep(List.Ctrl[,1], each = cellTypeslength)
plotOrderReceptorCells
cell.cell.interxns.ctrl <- paste(plotOrderLigandCells, plotOrderReceptorCells, sep = "-")

#now can use these as column labels for Xn, Yn, and XnYn
names(CtrlXn) <- cell.cell.interxns.ctrl
names(CtrlYn) <- cell.cell.interxns.ctrl
names(CtrlXnYn) <- cell.cell.interxns.ctrl

write.csv(CtrlXn, file = "/Users/mpcain/Desktop/Harmony/SeV.2wk.CtrlXn.allcellcombos.csv")
write.csv(CtrlYn, file = "/Users/mpcain/Desktop/Harmony/SeV.2wk.CtrlYn.allcellcombos.csv")
write.csv(CtrlXnYn, file = "/Users/mpcain/Desktop/Harmony/SeV.2wk.CtrlXnYn.allcellcombos.csv")

# Transpose so that cell type pairs are the columns and ligand-receptor pairs are the rows
Table.CtrlXn <- t(CtrlXn)
Table.CtrlYn <- t(CtrlYn)
Table.CtrlXnYn <- t(CtrlXnYn)

write.csv(Table.CtrlXn, file = "SeV.2wk.CtrlXn.allcellcombos-table.csv")
write.csv(Table.CtrlYn, file = "SeV.2wk.CtrlYn.allcellcombos-table.csv")
write.csv(Table.CtrlXnYn, file = "SeV.2wk.CtrlXnYn.allcellcombos-table.csv")

# creating empty matrices to fill using loop below
# X and Y alone for control and mutant form Figure 3 plots, while CtrlXnYn and MutXnYn will form Figure 4 plots
# INFECTED ####
MutXn <- data.frame(matrix(nrow = LRpair.num, ncol = cellTypeslength))
MutYn <- data.frame(matrix(nrow = LRpair.num, ncol = cellTypeslength))
MutXnYn <- data.frame(matrix(nrow = LRpair.num, ncol = cellTypeslength))

# Makes all x cooordinates for every plot in plot matrix.  each column is the ligands' expressions, in order of LRpairs list, for one cell type
for (i in 1:Plotsnum){
  m <- plotOrderX[i]
  v <- as.data.frame(unlist(List.Mut.ligands[m,]))
  
  MutXn[i] <- v[LRpair.ligand.order.locations.final,]
  rownames(MutXn) <- LRpairs$Pair.Name
}  

# Makes all y cooordinates for every plot in plot matrix.  each column is the receptors' expressions, in order of LRpairs list, for one cell type
for (j in 1:Plotsnum){
  l <- plotOrderY[j]
  t <- as.data.frame(unlist(List.Mut.receptors.reorder[l,]))
  
  MutYn[j] <- t[LRpair.receptor.order.locations.final,]
  rownames(MutYn) <- LRpairs$Pair.Name
}

# Makes all cooordinates for every plot in plot matrix for visualizationplot3.  each column is the product of the ligands' and receptors' expressions, in order of LRpairs list, for one cell type
for (i in 1:Plotsnum){
  m <- plotOrderX[i]
  l <- plotOrderY[i]
  MutXnYn[i] <- MutXn[,i]*MutYn[,i]
  rownames(MutXnYn) <- LRpairs$Pair.Name
}

### Makes correct order of x and y to make plots in 18 x 18 matrix - this time using cell type labels
# For X, want to repeat cell types 1 through 18
# For y, want to repeat cell type 1 eighteen times, then cell type 2 eighteen times, etc so that we get each cell-cell combination
plotOrderLigandCells.Mut <- rep.int(List.Mut[,1], cellTypeslength)
plotOrderLigandCells.Mut
plotOrderReceptorCells.Mut <- rep(List.Mut[,1], each = cellTypeslength)
plotOrderReceptorCells.Mut
cell.cell.interxns.mut <- paste(plotOrderLigandCells.Mut, plotOrderReceptorCells.Mut, sep = "-")

#now can use these as column labels for Xn, Yn, and XnYn
names(MutXn) <- cell.cell.interxns.mut
names(MutYn) <- cell.cell.interxns.mut
names(MutXnYn) <- cell.cell.interxns.mut

write.csv(MutXn, file = "SeV.2wk.MutXn.allcellcombos.csv")
write.csv(MutYn, file = "SeV.2wk.MutYn.allcellcombos.csv")
write.csv(MutXnYn, file = "SeV.2wk.MutXnYn.allcellcombos.csv")

# Transpose so that cell type pairs are the columns and ligand-receptor pairs are the rows - these are used to generate Figure3a graph for AT1-Car4 ECs
Table.MutXn <- t(MutXn)
Table.MutYn <- t(MutYn)
Table.MutXnYn <- t(MutXnYn)

write.csv(Table.MutXn, file = "SeV.2wk.MutXn.allcellcombos-table.csv")
write.csv(Table.MutYn, file = "SeV.2wk.MutYn.allcellcombos-table.csv")
write.csv(Table.MutXnYn, file = "SeV.2wk.MutXnYn.allcellcombos-table.csv")

# Determine outliers by applying 3 times interquartile range ####
# Values of zero skew outliers, so we decided to replace all 0 values with NA
# CONTROL OUTLIERS ####
Table.CtrlXnYn.0toNA <- Table.CtrlXnYn
Table.CtrlXnYn.0toNA[Table.CtrlXnYn.0toNA == 0] <- NA
Table.CtrlXnYn.IQR <- apply(Table.CtrlXnYn.0toNA, 2,IQR, na.rm=TRUE)
Table.CtrlXnYn.Q1 <- apply(Table.CtrlXnYn.0toNA, 2, quantile, probs=c(0.25), na.rm=TRUE)
Table.CtrlXnYn.Q3 <- apply(Table.CtrlXnYn.0toNA, 2, quantile, probs=c(0.75), na.rm=TRUE)

#Q3 + 1.5*IQR
Table.CtrlXnYn.upper1.5xIQR <- Table.CtrlXnYn.Q3 + (1.5*Table.CtrlXnYn.IQR)
#Q3 + 3*IQR
Table.CtrlXnYn.upper3xIQR <- Table.CtrlXnYn.Q3 + (3*Table.CtrlXnYn.IQR)
#Q1 - 1.5*IQR
Table.CtrlXnYn.lower1.5xIQR <- Table.CtrlXnYn.Q1 - (1.5*Table.CtrlXnYn.IQR)
#Q1 - 3*IQR
Table.CtrlXnYn.lower3xIQR <- Table.CtrlXnYn.Q1 - (3*Table.CtrlXnYn.IQR)

Table.CtrlXnYn.interquartile <- rbind(Table.CtrlXnYn.0toNA, Table.CtrlXnYn.upper1.5xIQR, Table.CtrlXnYn.upper3xIQR, Table.CtrlXnYn.lower1.5xIQR, Table.CtrlXnYn.lower3xIQR) 
write.csv(Table.CtrlXnYn.interquartile, file = "/Users/mpcain/Desktop/Harmony/Table.CtrlXnYn.interquartile.NA.csv")

# INFECTED OUTLIERS ####
Table.MutXnYn.0toNA <- Table.MutXnYn
Table.MutXnYn.0toNA[Table.MutXnYn.0toNA == 0] <- NA
Table.MutXnYn.IQR <- apply(Table.MutXnYn.0toNA, 2,IQR, na.rm=TRUE)
Table.MutXnYn.Q1 <- apply(Table.MutXnYn.0toNA, 2, quantile, probs=c(0.25), na.rm=TRUE)
Table.MutXnYn.Q3 <- apply(Table.MutXnYn.0toNA, 2, quantile, probs=c(0.75), na.rm=TRUE)

#Q3 + 1.5*IQR
Table.MutXnYn.upper1.5xIQR <- Table.MutXnYn.Q3 + (1.5*Table.MutXnYn.IQR)
#Q3 + 3*IQR
Table.MutXnYn.upper3xIQR <- Table.MutXnYn.Q3 + (3*Table.MutXnYn.IQR)
#Q1 - 1.5*IQR
Table.MutXnYn.lower1.5xIQR <- Table.MutXnYn.Q1 - (1.5*Table.MutXnYn.IQR)
#Q1 - 3*IQR
Table.MutXnYn.lower3xIQR <- Table.MutXnYn.Q1 - (3*Table.MutXnYn.IQR)

Table.MutXnYn.interquartile <- rbind(Table.MutXnYn.0toNA, Table.MutXnYn.upper1.5xIQR, Table.MutXnYn.upper3xIQR, Table.MutXnYn.lower1.5xIQR, Table.MutXnYn.lower3xIQR) 
write.csv(Table.MutXnYn.interquartile, file = "/Users/mpcain/Desktop/Harmony/Table.MutXnYn.interquartile.NA.csv")

# Apply 3 times IQR but return binary output, +1 if higher than 3 times IQR, -1 if lower than 3 times IQR ####
# Control ####
# HINT: Loop might not work if you do not read in table from .csv rather than use object already in environment
diff <- read.csv("/Users/mpcain/Desktop/Harmony/Table.CtrlXnYn.interquartile.NA.csv", header=T, row.names = 1)
cutoff <- diff[(nrow(diff)-3):nrow(diff),]
diff <- diff[1:(nrow(diff)-4),]

binary.diff <- diff
for (i in 1:ncol(diff)) {
  tmp <- diff[,i]
  low.limit <- quantile(tmp, probs=0.25, na.rm=T)-3*IQR(tmp, na.rm=T)
  up.limit <- quantile(tmp, probs=0.75, na.rm=T)+3*IQR(tmp, na.rm=T)
  binary.diff[,i] <- rep(0,nrow(diff))
  for (j in 1:nrow(diff)){
    if (!is.na(tmp[j])){
      if (tmp[j]<low.limit){binary.diff[j,i]=-1}
      if (tmp[j]>up.limit){binary.diff[j,i]=1}
    }
  }
}
write.csv(binary.diff, file = "/Users/mpcain/Desktop/Harmony/binary.Table.CtrlXnYn.interquartile.NA.csv", row.names = T)
write.csv(t(binary.diff), file = "/Users/mpcain/Desktop/Harmony/tbinary.Table.CtrlXnYn.interquartile.NA.csv", row.names = T)

# Figure 3c - heatmaps of Vegfa-Kdr and Pdgfb-Pdgfrb ####
#extract expression data for Vegfa for each cell type from control list
VEGFAexprsCtrl <- as.data.frame(List.Ctrl$Vegfa)

# carry over cell types from list to be row names associated with VEGFA expression data
rownames(VEGFAexprsCtrl) <- List.Ctrl[,1]

# unlist to make it easier to use later
VEGFAexprsCtrl.vector <- as.vector(unlist(VEGFAexprsCtrl))

# Do the same now for the receptor, Kdr
KDRexprsCtrl <- as.data.frame(List.Ctrl$Kdr)
rownames(KDRexprsCtrl) <- List.Ctrl[,1]
KDRexprsCtrl.vector <- as.vector(unlist(KDRexprsCtrl))

# get data in correct format for turning into matrix
Kdr <- cbind(rownames(KDRexprsCtrl), KDRexprsCtrl.vector)
Vegfa <- cbind(rownames(VEGFAexprsCtrl), VEGFAexprsCtrl.vector)
K3 <- as.numeric(Kdr[,2]) # tell R to treat expression values as numbers
V3 <- as.numeric(Vegfa[,2]) # tell R to treat expression values as numbers

# Create 2 matrices, one of repeating ligand expression data, one fo repeating receptor expression data, in order to multiply the two together for representation of ligand receptor interaction
# Need to make a matrix where the ligand expression data is repeated as 18 rows, once for each cell type
matrixVEGFAx <- as.matrix(t(replicate(18, V3)))
# Need to make a matrix where the receptor expression data is repeated as 18 columns, once for each cell type
matrixKDRy <- as.matrix(replicate(18, K3))

# Multiplication of ligand and receptor expression values for each cell-cell pair to make heatmap ####
VEGFA_KDR_matrix <- matrixVEGFAx*matrixKDRy

# ggplot2 uses inputed order of column names but puts rows in alphabetical order
# Need to get creative to put cell types in the order that you want them
# Row and column names are lost after matrix multiplication so need to add them back
VEGFA_KDR_df2 <- as.data.frame(VEGFA_KDR_matrix)
# below, the cell type names have to be in the same order as the expression matrices were made but can put numbers in front of names to reorder before plotting
# NOTE: start with 11 rather than 1 because ggplot will treat 11 as coming before 1 when ordering row names
LigandCluster3 <- paste(c("20Car4+ ECs","21Bulk ECs","22mature vessel ECs","11Pdgfra+ cells","12mesothelial cells","13pericytes","14Lum+ cells","15AVSM cells","16AT1 cells","17AT2 cells","18basal cells","19airway cells","23T cells","24myeloid cells","25AM","26granulocytes","27B cells","28NK cells"))
names(VEGFA_KDR_df2) <- LigandCluster3
rownames(VEGFA_KDR_df2) <- LigandCluster3
VEGFA_KDR_df2$ReceptorCluster <- LigandCluster3

# Orders rows by row name number order
test <- VEGFA_KDR_df2[order(VEGFA_KDR_df2$ReceptorCluster),]

# Orders columns by same order as row names
test2 <- test[,order(names(test))]

# set up data to use ggplot for heatmap
VEGFA_KDR_melt <- melt(test2, id.vars = "ReceptorCluster")
names(VEGFA_KDR_melt)[2:3] <- c("LigandCluster", "LR_Expression_Factor")

# creates heatmap based on "melted" matrix data
ggplot(VEGFA_KDR_melt, aes(LigandCluster, ReceptorCluster )) +
  geom_tile(aes(fill = LR_Expression_Factor), color = "white") +
  scale_fill_gradient(low = "#330066", high = "yellow") +
  ylab("KDR-expressing cell types") +
  xlab("VEGFA-expressing cell types") +
  theme(legend.title = element_text(size = 10),
        legend.text = element_text(size = 12),
        plot.title = element_text(size=16),
        axis.title=element_text(size=14,face="bold"),
        axis.text.x = element_text(angle = 90, hjust = 1)) +
  labs(fill = "L-R interaction strength")

# Figure 4c - Application of interactome to Sendia virus ####
# Multiplication of ligand and receptor values in L-R pair order, used to be called "Visualization Plot 3"
VizPlot3_Xvalues <- CtrlXn*CtrlYn
VizPlot3_Yvalues <- MutXn*MutYn

# difference: infected - control ####
ctrl.mut.diff <- VizPlot3_Yvalues - VizPlot3_Xvalues
write.csv(ctrl.mut.diff, file = "/Users/mpcain/Desktop/Harmony/difference.plot.csv")

# Transpose such that cell type pairs are the columns
Table.ctrl.mut.diff <- t(ctrl.mut.diff)
write.csv(Table.ctrl.mut.diff, file = "/Users/mpcain/Desktop/Harmony/difference.plot-table.csv")

# Determine outliers by applying 3 times interquartile range ####
# Values of zero skew outliers, so we decided to replace all 0 values with NA
# DIFFERENCE OUTLIERS ####
Table.ctrl.mut.diff.0toNA <- Table.ctrl.mut.diff
Table.ctrl.mut.diff.0toNA[Table.ctrl.mut.diff.0toNA == 0] <- NA
Table.ctrl.mut.diff.IQR <- apply(Table.ctrl.mut.diff.0toNA, 2,IQR, na.rm=TRUE)
Table.ctrl.mut.diff.Q1 <- apply(Table.ctrl.mut.diff.0toNA, 2, quantile, probs=c(0.25), na.rm=TRUE)
Table.ctrl.mut.diff.Q3 <- apply(Table.ctrl.mut.diff.0toNA, 2, quantile, probs=c(0.75), na.rm=TRUE)

#Q3 + 1.5*IQR
Table.ctrl.mut.diff.upper1.5xIQR <- Table.ctrl.mut.diff.Q3 + (1.5*Table.ctrl.mut.diff.IQR)
head(Table.ctrl.mut.diff.upper1.5xIQR)
#Q3 + 3*IQR
Table.ctrl.mut.diff.upper3xIQR <- Table.ctrl.mut.diff.Q3 + (3*Table.ctrl.mut.diff.IQR)
head(Table.ctrl.mut.diff.upper3xIQR)
#Q1 - 1.5*IQR
Table.ctrl.mut.diff.lower1.5xIQR <- Table.ctrl.mut.diff.Q1 - (1.5*Table.ctrl.mut.diff.IQR)
head(Table.ctrl.mut.diff.lower1.5xIQR)
#Q1 - 3*IQR
Table.ctrl.mut.diff.lower3xIQR <- Table.ctrl.mut.diff.Q1 - (3*Table.ctrl.mut.diff.IQR)
head(Table.ctrl.mut.diff.lower3xIQR)

Table.ctrl.mut.diff.interquartile <- rbind(Table.ctrl.mut.diff.0toNA, Table.ctrl.mut.diff.upper1.5xIQR, Table.ctrl.mut.diff.upper3xIQR, Table.ctrl.mut.diff.lower1.5xIQR, Table.ctrl.mut.diff.lower3xIQR) 
write.csv(Table.ctrl.mut.diff.interquartile, file = "/Users/mpcain/Desktop/Harmony/Table.ctrl.mut.diff.interquartile.csv")

# Apply 3 times IQR but return binary output, +1 if higher than 3 times IQR, -1 if lower than 3 times IQR ####
# Differences ####
# HINT: Loop might not work if you do not read in table from .csv rather than use object already in environment
diff <- read.csv("/Users/mpcain/Desktop/Harmony/Table.ctrl.mut.diff.interquartile.csv", header=T, row.names = 1)
cutoff <- diff[(nrow(diff)-3):nrow(diff),]
diff <- diff[1:(nrow(diff)-4),]

binary.diff <- diff
for (i in 1:ncol(diff)) {
  tmp <- diff[,i]
  low.limit <- quantile(tmp, probs=0.25, na.rm=T)-3*IQR(tmp, na.rm=T)
  up.limit <- quantile(tmp, probs=0.75, na.rm=T)+3*IQR(tmp, na.rm=T)
  binary.diff[,i] <- rep(0,nrow(diff))
  for (j in 1:nrow(diff)){
    if (!is.na(tmp[j])){
      if (tmp[j]<low.limit){binary.diff[j,i]=-1}
      if (tmp[j]>up.limit){binary.diff[j,i]=1}
    }
  }
}
write.csv(binary.diff, file = "/Users/mpcain/Desktop/Harmony/binary.Table.ctrl.mut.diff.interquartile.csv", row.names = T)
write.csv(t(binary.diff), file = "/Users/mpcain/Desktop/Harmony/tbinary.Table.ctrl.mut.diff.interquartile.csv", row.names = T)
```
