## Supplemental File 2 for "Intermediary role of lung alveolar type 1 cells in epithelial repair upon Sendai virus infection": Step3.three.timepoint.outlier.comparisons.R

```
# R code
# Margo P. Cain
# Code will generate data used to make Hernandez et al. Figure 6d. This code will pull out common and unique outlying interactions from multiple samples.

# Makes the assumption that colnames are the same and that cell type pairs are named the same.

# Load R libs ####
library(Seurat)
library(reshape2)

# 1 week & 2 week outlying interaction comparison ####
# outlying.interaxns.1 - SeV-1 week
# outlying.interaxns.2 - SeV-2 weeks
# outlying.interaxns.3 - SeV-7 weeks

outlying.interaxns.1 <- read.csv("//mdanderson.edu/MDAHomes/users37/mpcain/MPCain-server/Harmony.1wk/tbinary.Table.ctrl.mut.diff.interquartile.csv", row.names = 1)
outlying.interaxns.2 <- read.csv("//mdanderson.edu/MDAHomes/users37/mpcain/MPCain-server/Harmony/tbinary.Table.ctrl.mut.diff.interquartile.csv", row.names = 1)
outlying.interaxns.3 <- read.csv("//mdanderson.edu/MDAHomes/users37/mpcain/MPCain-server/Harmony.7wk/tbinary.Table.ctrl.mut.diff.interquartile.csv", row.names = 1)

fold.change.1 <- read.csv("//mdanderson.edu/MDAHomes/users37/mpcain/MPCain-server/Harmony.1wk/difference.plot.csv", row.names = 1)
fold.change.2 <- read.csv("//mdanderson.edu/MDAHomes/users37/mpcain/MPCain-server/Harmony/difference.plot.csv", row.names = 1)
fold.change.3 <- read.csv("//mdanderson.edu/MDAHomes/users37/mpcain/MPCain-server/Harmony.7wk/difference.plot.csv", row.names = 1)

# extracts names of the cell-cell pairs in order to isolate only cell pairs containing AT1 cells
cell.pairs.1 <- names(outlying.interaxns.1)
cell.pairs.2 <- names(outlying.interaxns.2)
cell.pairs.3 <- names(outlying.interaxns.3)

containsAT1.1 <- which(grepl("AT1", cell.pairs.1, fixed=TRUE)) # locations of cell type pairs containing AT1 cells
AT1.cell.pairs.1 <- cell.pairs.1[containsAT1.1]
#cell.pairs.1[containsAT1.1] # manual check to make sure order is correct

containsAT1.2 <- which(grepl("AT1", cell.pairs.2, fixed=TRUE)) # locations of cell type pairs containing AT1 cells
AT1.cell.pairs.2 <- cell.pairs.2[containsAT1.2]
#cell.pairs.2[containsAT1.2] # manual check to make sure order is correct

containsAT1.3 <- which(grepl("AT1", cell.pairs.3, fixed=TRUE)) # locations of cell type pairs containing AT1 cells
AT1.cell.pairs.3 <- cell.pairs.3[containsAT1.3]

match.go.in.1 <- which(AT1.cell.pairs.1 %in% AT1.cell.pairs.2) # locations where cell type pairs match each other
match.go.in.2 <- which(AT1.cell.pairs.2 %in% AT1.cell.pairs.1)
cbind(AT1.cell.pairs.1[match.go.in.1], AT1.cell.pairs.2[match.go.in.2]) # visual manual check that they match

# Now have to get correct matching locations to call outliers from correct columns (column is cell type pairs)
# below will be the locations in 'outlying.interaxns' of AT1 cell pairs matching between the two samples
x <- containsAT1.1[match.go.in.1]
cell.pairs.1[x] # manual check, should be same cell type pairs as cbind manual check above

y <- containsAT1.2[match.go.in.2]
cell.pairs.2[y] # manual check, should be same cell type pairs as cbind manual check above


z <- containsAT1.3[match.go.in.2]
cell.pairs.3[z] # manual check, should be same cell type pairs as cbind manual check above

# call locations of matching cell pairs from outlying interactions data
AT1.outlying.interaxns.matched.1 <- outlying.interaxns.1[,x]
AT1.outlying.interaxns.matched.2 <- outlying.interaxns.2[,y]
AT1.outlying.interaxns.matched.3 <- outlying.interaxns.3[,z]


# Fold Change ####

# isolate the AT1 cells only, signaling and receiving
cell.pairs.1 <- names(fold.change.1)
cell.pairs.2 <- names(fold.change.2)
cell.pairs.3 <- names(fold.change.3)

containsAT1.1 <- which(grepl("AT1", cell.pairs.1, fixed=TRUE)) # locations of cell type pairs containing AT1 cells
AT1.cell.pairs.1 <- cell.pairs.1[containsAT1.1]
#cell.pairs.1[containsAT1.1] # manual check to make sure order is correct

containsAT1.2 <- which(grepl("AT1", cell.pairs.2, fixed=TRUE)) # locations of cell type pairs containing AT1 cells
AT1.cell.pairs.2 <- cell.pairs.2[containsAT1.2]
#cell.pairs.2[containsAT1.2] # manual check to make sure order is correct

containsAT1.3 <- which(grepl("AT1", cell.pairs.3, fixed=TRUE)) # locations of cell type pairs containing AT1 cells
AT1.cell.pairs.3 <- cell.pairs.3[containsAT1.3]

match.go.in.1 <- which(AT1.cell.pairs.1 %in% AT1.cell.pairs.2) # locations where cell type pairs match each other
match.go.in.2 <- which(AT1.cell.pairs.2 %in% AT1.cell.pairs.1)
cbind(AT1.cell.pairs.1[match.go.in.1], AT1.cell.pairs.2[match.go.in.2]) # manual check that they match

# Now have to get correct matching locations to call outliers from correct columns (cell type pairs)
# Below will be the locations in outlying.interaxns of AT1 cell pairs matching between the two samples
x <- containsAT1.1[match.go.in.1]
cell.pairs.1[x] # manual check, should be same cell type pairs as cbind manual check above

y <- containsAT1.2[match.go.in.2]
cell.pairs.2[y] # manual check, should be same cell type pairs as cbind manual check above

z <- containsAT1.3[match.go.in.2]
cell.pairs.3[z] # manual check, should be same cell type pairs as cbind manual check above

AT1.fold.change.matched.1 <- fold.change.1[,x]
AT1.fold.change.matched.2 <- fold.change.2[,y]
AT1.fold.change.matched.3 <- fold.change.3[,z]

# Combine into one matrix
unlist.AT1.outlying.interaxns.matched.1 <- unlist(AT1.outlying.interaxns.matched.1)
unlist.AT1.fold.change.matched.1 <- unlist(AT1.fold.change.matched.1)
# Here, checked by comparing to AT1.outlying.interaxns.matched.1

# Remove interactions with a score of 0, which are not outlying - time point two
unlist.AT1.outlying.interaxns.matched.2 <- unlist(AT1.outlying.interaxns.matched.2)
unlist.AT1.fold.change.matched.2 <- unlist(AT1.fold.change.matched.2)
# Here, checked by comparing to AT1.outlying.interaxns.matched.2

unlist.AT1.outlying.interaxns.matched.3 <- unlist(AT1.outlying.interaxns.matched.3)
unlist.AT1.fold.change.matched.3 <- unlist(AT1.fold.change.matched.3)

rep.interaxn.names <- rownames(AT1.outlying.interaxns.matched.1)

# Combines outlying signaling interaction names, scores, and fold-chage values
AT1.df.combined.2 <- cbind(rep.interaxn.names, as.data.frame(unlist.AT1.outlying.interaxns.matched.1), as.data.frame(unlist.AT1.fold.change.matched.1), as.data.frame(unlist.AT1.outlying.interaxns.matched.2), as.data.frame(unlist.AT1.fold.change.matched.2), as.data.frame(unlist.AT1.outlying.interaxns.matched.3), as.data.frame(unlist.AT1.fold.change.matched.3))
write.csv(AT1.df.combined.2, "//mdanderson.edu/MDAHomes/users37/mpcain/MPCain-server/master.list.1wk2wk7wk.csv")

# Separates outlying interactions based on whether scores are outlying for one or both time points, and whether the score is the same or opposite

red <- subset(AT1.df.combined.2, AT1.df.combined.2[,2] != 0 & AT1.df.combined.2[,4] == 0 & AT1.df.combined.2[,6] == 0)
green <- subset(AT1.df.combined.2, AT1.df.combined.2[,2] == 0 & AT1.df.combined.2[,4] != 0 & AT1.df.combined.2[,6] == 0)
blue <- subset(AT1.df.combined.2, AT1.df.combined.2[,2] == 0 & AT1.df.combined.2[,4] == 0 & AT1.df.combined.2[,6] != 0)

yellow <- subset(AT1.df.combined.2, AT1.df.combined.2[,2] != 0 & AT1.df.combined.2[,4] != 0 & AT1.df.combined.2[,6] == 0)
magenta <- subset(AT1.df.combined.2, AT1.df.combined.2[,2] != 0 & AT1.df.combined.2[,4] == 0 & AT1.df.combined.2[,6] != 0)
cyan <- subset(AT1.df.combined.2, AT1.df.combined.2[,2] == 0 & AT1.df.combined.2[,4] != 0 & AT1.df.combined.2[,6] != 0)

gray <- subset(AT1.df.combined.2, AT1.df.combined.2[,2] != 0 & AT1.df.combined.2[,4] != 0 & AT1.df.combined.2[,6] != 0)

# Creates .csv files for each relationship
write.csv(red, "//mdanderson.edu/MDAHomes/users37/mpcain/MPCain-server/Harmony/red.csv")
write.csv(green, "//mdanderson.edu/MDAHomes/users37/mpcain/MPCain-server/Harmony/green.csv")
write.csv(blue, "//mdanderson.edu/MDAHomes/users37/mpcain/MPCain-server/Harmony/blue.csv")
write.csv(yellow, "//mdanderson.edu/MDAHomes/users37/mpcain/MPCain-server/Harmony/yellow.csv")
write.csv(magenta, "//mdanderson.edu/MDAHomes/users37/mpcain/MPCain-server/Harmony/magenta.csv")
write.csv(cyan, "//mdanderson.edu/MDAHomes/users37/mpcain/MPCain-server/Harmony/cyan.csv")
write.csv(gray, "//mdanderson.edu/MDAHomes/users37/mpcain/MPCain-server/Harmony/gray.csv")


# 2 week & 7 week outlying interaction comparison - for the 2 basal cell type interactions that are missing in the week 1 data ####
# outlying.interaxns.2 - SeV-2 week
# outlying.interaxns.3 - SeV-7 weeks

match.go.in.2 <- which(AT1.cell.pairs.2 %in% AT1.cell.pairs.3) # locations where cell type pairs match each other
match.go.in.3 <- which(AT1.cell.pairs.3 %in% AT1.cell.pairs.2)
cbind(AT1.cell.pairs.2[match.go.in.2], AT1.cell.pairs.3[match.go.in.3]) # visual manual check that they match

# Now have to get correct matching locations to call outliers from correct columns (column is cell type pairs)
# below will be the locations in 'outlying.interaxns' of AT1 cell pairs matching between the two samples
y <- containsAT1.2[match.go.in.2]
cell.pairs.2[y] # manual check, should be same cell type pairs as cbind manual check above

z <- containsAT1.3[match.go.in.3]
cell.pairs.3[z] # manual check, should be same cell type pairs as cbind manual check above

# call locations of matching cell pairs from outlying interactions data
AT1.outlying.interaxns.matched.2 <- outlying.interaxns.2[,y]
AT1.outlying.interaxns.matched.3 <- outlying.interaxns.3[,z]

# Fold Change ####

# isolate the AT1 cells only, signaling and receiving
match.go.in.2 <- which(AT1.cell.pairs.2 %in% AT1.cell.pairs.3) # locations where cell type pairs match each other
match.go.in.3 <- which(AT1.cell.pairs.3 %in% AT1.cell.pairs.2)
cbind(AT1.cell.pairs.2[match.go.in.2], AT1.cell.pairs.3[match.go.in.3]) # manual check that they match

# Now have to get correct matching locations to call outliers from correct columns (cell type pairs)
# Below will be the locations in outlying.interaxns of AT1 cell pairs matching between the two samples
y <- containsAT1.2[match.go.in.2]
cell.pairs.2[y] # manual check, should be same cell type pairs as cbind manual check above

z <- containsAT1.3[match.go.in.3]
cell.pairs.3[z] # manual check, should be same cell type pairs as cbind manual check above

AT1.fold.change.matched.2 <- fold.change.2[,y]
AT1.fold.change.matched.3 <- fold.change.3[,z]

# Combine into one matrix
unlist.not.zero.1 <- as.vector(not.zero.1)
unlist.AT1.outlying.interaxns.matched.1 <- unlist(AT1.outlying.interaxns.matched.1)
unlist.AT1.fold.change.matched.1 <- unlist(AT1.fold.change.matched.1)
AT1.df.1 <- cbind(rep.interaxn.names, as.data.frame(unlist.not.zero.1), as.data.frame(unlist.AT1.outlying.interaxns.matched.1), as.data.frame(unlist.AT1.fold.change.matched.1))
#write.csv(AT1.df.1, "/Users/mpcain/Desktop/AT1.df.1.csv")
# Here, checked by comparing to AT1.outlying.interaxns.matched.1

# Remove interactions with a score of 0, which are not outlying - time point two
not.zero.2 <- AT1.outlying.interaxns.matched.2 !=0
unlist.not.zero.2 <- as.vector(not.zero.2)
unlist.AT1.outlying.interaxns.matched.2 <- unlist(AT1.outlying.interaxns.matched.2)
unlist.AT1.fold.change.matched.2 <- unlist(AT1.fold.change.matched.2)
AT1.df.2 <- cbind(rep.interaxn.names, as.data.frame(unlist.not.zero.2), as.data.frame(unlist.AT1.outlying.interaxns.matched.2))
#write.csv(AT1.df.2, "/Users/mpcain/Desktop/AT1.df.2.csv")
# Here, checked by comparing to AT1.outlying.interaxns.matched.2

# Combines outlying signaling interaction names, scores, and fold-chage values
AT1.df.combined <- cbind(rep.interaxn.names, as.data.frame(unlist.not.zero.1), as.data.frame(unlist.AT1.outlying.interaxns.matched.1), as.data.frame(unlist.AT1.fold.change.matched.1),as.data.frame(unlist.not.zero.2), as.data.frame(unlist.AT1.outlying.interaxns.matched.2), as.data.frame(unlist.AT1.fold.change.matched.2))
AT1.df.combined.2 <- cbind(rep.interaxn.names, as.data.frame(unlist.AT1.outlying.interaxns.matched.1), as.data.frame(unlist.AT1.fold.change.matched.1), as.data.frame(unlist.AT1.outlying.interaxns.matched.2), as.data.frame(unlist.AT1.fold.change.matched.2))
```
